# Electrostatic changes enabled the diversification of an exocyst subunit via protein complex escape

**DOI:** 10.1101/2024.08.26.609756

**Authors:** Juan Carlos De la Concepcion, Héloïse Duverge, Yoonwoo Kim, Jose Julian, Haonan D. Xu, Sara Ait Ikene, Anita Bianchi, Nenad Grujic, Ranjith K. Papareddy, Irina Grishkovskaya, David Haselbach, David H. Murray, Marion Clavel, Nicholas A. T. Irwin, Yasin Dagdas

**Affiliations:** Gregor Mendel Institute, Austrian Academy of Sciences, Vienna BioCenter, Vienna, Austria; Department of Applied Genetics and Cell Biology, Institute of Molecular Plant Biology, Boku University; Vienna, 1190, Austria; Division of Molecular Cell and Developmental Biology, School of Life Sciences, University of Dundee, Dow Street, Dundee DD1 5EH, UK; Research Institute of Molecular Pathology, Vienna BioCenter, Vienna, Austria; Max-Planck-Institut für Molekulare Pflanzenphysiologie, Potsdam-Golm, Germany

## Abstract

The evolution of cellular complexity hinges on the capacity of multimeric protein complexes to diversify without compromising their ancestral functions. A central question is how individual subunits within these complexes can evolve novel functions while maintaining the integrity of the original assembly. Here, we explore this question by tracing the evolutionary trajectory of the plant exocyst, an octameric complex essential for exocytosis in eukaryotes. Remarkably, the Exo70 subunit underwent dramatic expansion and functional divergence in plants. We demonstrate that electrostatic alterations in the N-terminal region of the Exo70 subunit precipitated its dissociation from the exocyst complex. This release mitigated paralog interference, thereby facilitating the subunit’s extensive functional co-option. Our findings reveal a nuanced mechanism by which a protein subunit, ancestrally constrained within a multimeric complex, can escape those constraints and evolve novel functions, shedding light on the molecular underpinnings of cellular innovation.

**One-Sentence Summary:** Evolutionary diversification of an exocyst subunit is driven by electrostatic shifts that dissociates it from the ancestral complex.

## Main text

The evolution of protein complexes represents a pivotal event in cellular evolution, frequently driven by the duplication and subsequent specialization of pre-existing proteins (*1–4*). The emergence of multimeric protein assemblies typically occurs through the gradual accumulation of stabilizing mutations, which establish crucial protein-protein interaction interfaces and facilitate the emergence of higher-order structures (*5–9*). This trend towards increased complexity may arise from adaptive or non-adaptive processes through mechanisms like constructive neutral evolution (*6, 7, 10–14*). Therefore, the evolution of protein complexes necessitates a delicate balance between maintaining the integrity of the complex and allowing for diversification.

While the evolution of multimeric complexes contributes to cellular diversity, it simultaneously imposes evolutionary constraints on individual proteins. The entrenchment of complex subunits reduces their evolutionary flexibility (*6, 7, 9, 10*) and the duplication and diversification of a single subunit can disrupt the stoichiometry and function of the protein complex, a deleterious phenomenon known as paralog interference (*15–17*). This conflict can be mitigated through spatiotemporal segregation of paralogs, achieved via differential gene expression or altered subcellular localization (*17, 18*). However, the lack of functional autonomy among subunits, coupled with the ubiquitous expression of many conserved protein complexes (*19–24*), limits opportunities for paralog isolation. The challenge is compounded by hydrophobic entrenchment, where hydrophobic residues accumulate neutrally at interaction interfaces, inadvertently hindering subunit dissociation (*6, 9*). Although subunit duplication is observed in homo-multimeric complexes (*4, 12, 25, 26*), the mechanisms by which diversification circumvents evolutionary constraints in heteromeric complexes remain poorly understood. In particular, the diversification of protein complexes that define basic biological processes has not been functionally characterized within a robust phylogenetic framework.

The exocyst complex presents a paradoxical example of subunit evolution. The exocyst is a highly conserved hetero-oligomeric complex that plays a critical role in exocytosis across eukaryotes by orchestrating the tethering and membrane fusion of secretory vesicles (*27–32*). The exocyst is composed of eight subunits—SEC3, SEC5, SEC6, SEC8, SEC10, SEC15, Exo84, and Exo70 (*33*) —which are organized through the coiling of their N-terminal α-helices into a helical bundle, known as the CorEx (*32, 34–36*). While most eukaryotes encode single copies of these subunits, an exception is found in plants, where Exo70, the subunit responsible for targeting the exocyst to the plasma membrane, has undergone extensive diversification. This subunit has expanded into dozens of paralogs, grouped into three families: Exo70I, Exo70II, and Exo70III (*37–40*) **(Figure S1)**.

Despite the variation in expression profiles, multiple Exo70 paralogs are co-expressed within individual cells (*41, 42*) where they contribute to both canonical and non-canonical exocytic functions (*37, 43*). Paralogs of the Exo70I family, which is relatively conserved and consists primarily of Exo70 clade A, are predominantly associated with canonical exocytosis (*44–47*). In contrast, paralogs from the highly diversified Exo70II family, encompassing clades B, C, D, E, F, and H, participate in a broad array of distinct cellular processes. For instance, Exo70s from clades B and D are implicated in autophagy (*48–50*), with AtExo70D1, AtExo70D2, and AtExo70D3 specifically functioning as selective cargo receptors (*50*), a role that does not necessarily require interaction with the exocyst complex. In monocots, the Exo70 clade F is the most extensively expanded and plays a crucial role in pathogen recognition (*51, 52*), and even serve as integrated domains in immune receptors (*39, 52–54*). The involvement of other exocyst components in these diverse processes remains unclear.

While some degree of Exo70 diversification can be attributed to functional specialization, such as variations in membrane targeting (*37, 43*), the mechanisms by which the extensive radiation of Exo70s circumvented paralog interference remain elusive. To unravel how Exo70 diversification navigated the evolutionary constraints typically associated with protein complex evolution, we explored the function and evolution of Exo70, using the plant model *Marchantia polymorpha* (hereafter Marchantia) (*55*). Marchantia encodes a single copy of the exocyst subunits and only three Exo70 paralogs, each representing one of the major plant Exo70 subfamilies **(Figure S1)**. This reduced genetic redundancy makes Marchantia an ideal model for dissecting Exo70 evolution, especially when compared to species like *Arabidopsis thaliana*, which possesses 23 Exo70 paralogs (*38*).

### Exo70 paralogs are functionally diverse in *Marchantia polymorpha*

To investigate the differences between Marchantia Exo70 paralogs, we first analyzed published gene expression data derived from different developmental stages and tissues (*56*). This revealed that MpExo70I and II are highly expressed and positively correlated with one another relative to MpExo70III **(Figure S2)**, suggesting that differential expression may isolate MpExo70III, while MpExo70I and MpExo70II co-occur, which could lead to paralog interference. To understand how this potential conflict is resolved, we compared the sub-cellular localizations of the Exo70 paralogs. To this end, we used the cell plate in post-mitotic cells as a proxy for exocyst localization as the exocyst complex was shown to accumulate at the cell plate in *Arabidopsis thaliana* (*44*). As in Arabidopsis, Marchantia core exocyst subunits, MpSec6 and MpExo84, both localized to the cell plate **(Figure S3, Figure S4)**. Constitutively expressed MpExo70I and MpExo70III also accumulated at the cell plate and co-localized with MpSEC6 and MpExo84 **(Figure 1a, Figure S5, Figure S6, Figure S7)**. However, MpExo70II had a diffuse localization, distinct from MpSec6, MpExo84, MpExo70I and MpExo70III **(Figure 1a, Figure S5, Figure S6, Figure S7).** These results suggest that MpExo70III likely avoids paralog interference through differential gene expression while MpExo70I and II are spatially segregated.

**Fig. 1.**
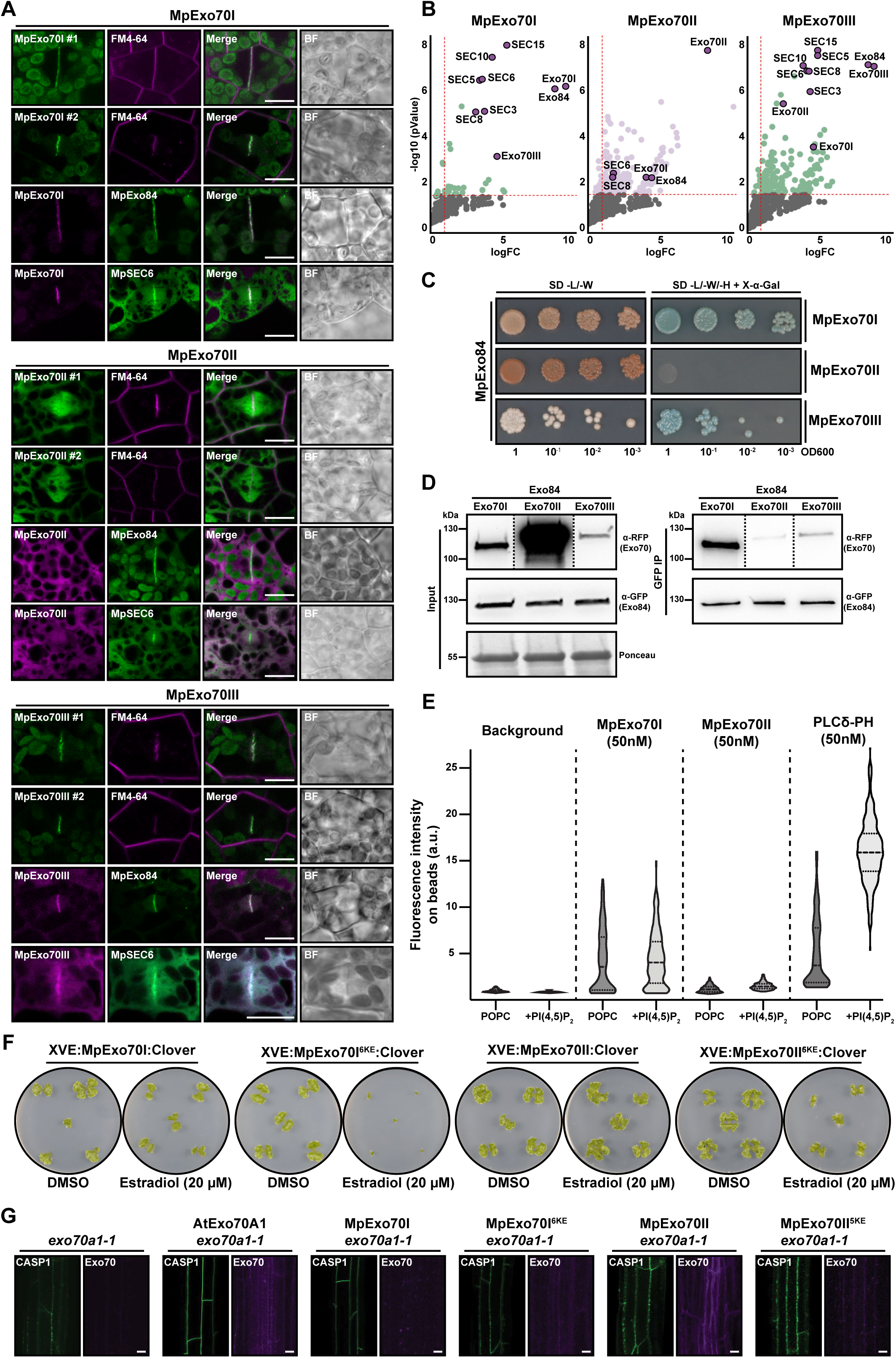
Marchantia Exo70 are functionally diverse. **(A)** Confocal micrographs of Marchantia cells from two independent cell lines stably expressing MpExo70:Clover (green) and stained with FM4-64 (magenta) and cell lines stably co-expressing MpExo70:mScarlet (magenta) with MpSEC6: Clover or MpExo84:Clover (green). The presence of the cell plate is depicted by accumulation of FM4-64 stain or the core exocyst components MpSEC6 or MpExo84. Scale bar is 10 µm. BF indicates Bright Field. For each line, a gallery of images was collected to document variability and are presented in **Figure S5** and **Figure S6** for FM4-64 treatment and co-localization, respectively. **(B)** Enrichment of exocyst protein co-purified with MpExo70I, MpExo70II or MpExo70III compared to a control expressing free GFP and represented by a volcano plot. The horizontal dashed line indicates the threshold above which proteins are significantly enriched (p value < 0.05, quasi-likelihood negative binomial generalized log-linear model) and the vertical dashed line the threshold for which proteins log2 fold change is above 1. For each plot, members of Marchantia exocyst complex are depicted by a red dot with the corresponding name. **(C)** Yeast two-hybrid assay of MpExo70 interaction with MpExo84. For each combination, 5µl of yeast at indicated OD600 were spotted and incubated in double dropout plate for yeast growth control (left) and triple dropout media supplemented with X-α-gal (right). Growth and development of blue coloration in the right panel indicates protein-protein interactions. MpExo70s were fused to the GAL4 DNA binding domain while MpExo84 was fused to the GAL4 activator domain. Accumulation of each protein in yeast cells was tested by western blot and presented in **Figure S14**. **(D)** Co-immunoprecipitation of MpExo70 proteins with MpExo84. C-terminally mScarlet-tagged MpExo70 proteins were stably co-expressed with C-terminally Clover-tagged MpExo84 in Marchantia. Immunoprecipitates (IPs) were obtained with anti-GFP magnetic beads and total protein extracts were probed with anti-RFP or anti-GFP antibodies for MpExo70 and MpExo84 proteins, respectively. Dashed lines represent a crop and assembled image from the same blot. **(E)** Evaluation of MpExo70 membrane binding at 50 nM. Fluorescence intensity of membrane associated GFP:MpExo70I, GFP:MpExo70II or Alexa488:PLCδ-PH domain were imaged by confocal microscopy. Representative confocal images are presented in **Figure S17**. Violin plots with median and quartiles are derived from 247-539 individual beads. **(F)** Macroscopic phenotypes of Marchantia transgenic lines XVE:MpExo70I:Clover, XVE:MpExo70I 6KE:Clover, XVE:MpExo70II:Clover and XVE:MpExo70II 6KE:Clover grown for 14 days on media containing B-estradiol or DMSO. Images are representative of experimental replicates presented in **Figure S20**. **(G)** CASP1:GFP accumulation at the Casparian strips of mature endodermis in Arabidopsis *exo70a1-1* (*45*) mutants lines (green) complemented with AtExo70A1, MpExo70I, MpExo70I 6KE, MpExo70II or MpExo70II 6KE (magenta). Dotted mis-localization of CASP1:GFP is representative of defect in Exo70 exocytic function (*45*). Scale bar is 10 µm.

Although spatial separation may relieve paralog interference, altered subcellular localization is likely to coincide with functional differentiation. To test whether MpExo70 paralogs have functionally diversified, we first compared their interactomes using immunoprecipitation coupled mass spectrometry (IP-MS). IP-MS analyses of MpExo70s as well as MpSec6 and MpExo84 revealed that MpExo70I only associates with a small subset of core exocyst proteins, whereas MpExo70II and MpExo70III have broader interactomes with limited overlap with each other and MpExo70I **(Figure S8, Figure S9, Figure S10, Supplemental Dataset 1, Supplemental dataset 2)**. Together, these data suggest that the Exo70 paralogs may have functionally diversified through altered protein-protein interactions.

A striking observation from these data was that whereas MpExo70I and MpExo70III strongly associated with the core exocyst components, these interactions were largely absent in MpExo70II interactome (**Figure 1b, Figure S11, Figure S12**). To further validate the apparent dissociation between MpExo70II and the exocyst complex, we directly assessed the interactions between each MpExo70 paralog and MpExo84, the core component that hetero-dimerize with Exo70 at the exocyst CorEx (*35, 36*). Indeed, a genome-wide Yeast-two-hybrid (Y2H) screen confirmed MpExo84 as the primary interactor of MpExo70I **(Figure S13)**. Similarly, pairwise Y2H identified interactions between Exo84 and MpExo70I and MpExo70III. However, MpExo70II did not interact with MpExo84 **(Figure 1c, Figure S14)**. Consistent with our Y2H and IP-MS results, in planta co-immunoprecipitation assays demonstrated that MpExo70II association with MpExo84 is far weaker than MpExo70I and MpExo70III **(Figure 1d)**. To further test whether MpExo70II can integrate into the exocyst at all, we expressed and purified Marchantia exocyst subcomplex 2 containing MpSEC10, MpSEC15, MpExo84 and either MpExo70I (SC2-I) or MpExo70II (SC2-II) from insect cells (*27, 35, 36*). Purified SC2-I had a homogeneous distribution containing all four subunits in stoichiometric amounts, whereas purified SC2-II seemed to lack MpExo70II, resulting in a heterogeneous protein population **(Figure S15)**. Altogether, these results demonstrate that MpExo70II no longer interacts with MpExo84 and the exocyst itself, implying that this protein has seemingly escaped its original context, likely alleviating any interference with other Exo70 paralogs.

The canonical function of Exo70 is recruiting the exocyst to the plasma membrane via phospholipid binding (*27, 46, 57*). To determine whether the functional diversification have arisen due to differential lipid binding abilities, we purified GFP:MpExo70I and GFP:MpExo70II from insect cells **(Figure S16)** and tested their ability to bind phospholipids. Both Exo70 paralogs bound lipids, although MpExo70I had higher affinity than MpExo70II **(Figure 1g, Figure S17)** (*27*). This indicates that, functional diversification is not due to differential lipid binding, but likely due to the dissociation of MpExo70II from the exocyst.

We next tested for functional divergence between MpExo70I and MpExo70II. As we were unable to obtain CRISPR mutants of any of MpExo70 paralogs, we tested the effects of blocking exocytosis in Marchantia using Endosidin 2 (ES2), an inhibitor that targets Exo70 to disrupt canonical exocytosis (*58*). Similar to Arabidopsis, ES2 treatment resulted in growth arrest, confirming the essentiality of Exo70-mediated exocytosis in Marchantia **(Figure S18)**. To investigate the individual contributions of MpExo70I and MpExo70II, we first generated dominant negative variants of both paralogs by replacing the conserved lysine residues (K) in the lipid binding site with glutamic acid (E) to prevent lipid binding (**Figure S19)** (*46, 57*). Next, we overexpressed MpExo70I KE or MpExo70II KE and assessed plant phenotypes. Inducible over-expression of MpExo70I KE phenocopied ES2 treatment, leading to growth arrest **(Figure 1f, Figure S20)**. In contrast, expression of MpExo70II KE only mildly effected growth and instead led to extensive proliferation of rhizoids **(Figure 1f, Figure S20)**. In agreement with our previous results, these alternative outcomes suggest that MpExo70I acts in canonical exocytosis, whereas MpExo70II functions in a different pathway.

Finally, to corroborate these functional differences, we assessed whether MpExo70 paralogs could complement exocytosis-related defects in an *exo70a1* mutant in *Arabidopsis thaliana*. AtExo70A1 is the ortholog of MpExo70I and its disruption results in multiple phenotypes (*46, 47*), including defects in AtCASP1 protein deposition at the Casparian strip in Arabidopsis roots (*45*). Consistent with a role in canonical exocytosis as part of the exocyst complex, MpExo70I complemented CASP1-GFP deposition defects, and this complementation required a functional lipid binding domain. On the contrary, neither MpExo70II nor MpExo70II KE could complement CASP1 deposition defects **(Figure 1g)**. Taken together, these results suggest that MpExo70II is not part of the exocyst complex and does not play a role in exocytosis.

### Variation in the Exo70 N-terminal domain impacts exocyst association

Next, we sought to determine the molecular and structural features underlying MpExo70II diversification. First, we took advantage of the structural similarities between Exo70s (*59*) and designed a series of chimeric proteins by swapping structurally equivalent regions between MpExo70I and MpExo70II proteins **(Figure 2a)**. Live cell imaging analysis of Marchantia lines expressing chimeras revealed that swapping the N-terminal region between MpExo70I and MpExo70II (residues Met1 to Ser154 in MpExo70I and Met1 to Ser169 in MpExo70II) was sufficient to switch their localizations. MpExo70I Chimera 1 had a diffuse localization pattern similar to MpExo70II, while MpExo70II Chimera 1 accumulated at the cell plate and co-localized with MpExo84 **(Figure 2b, Figure S21, Figure S22, Figure S23)**.

**Fig. 2.**
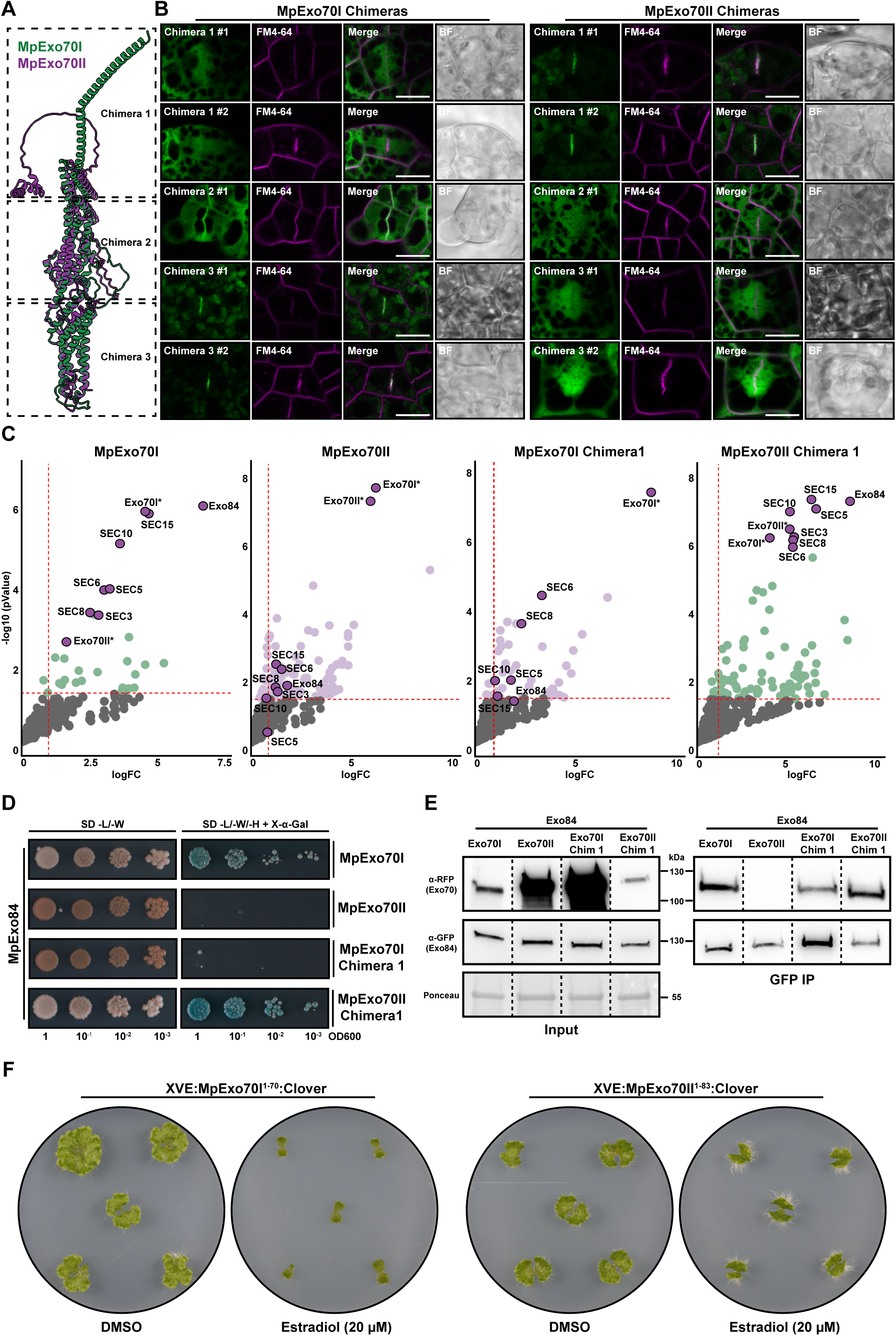
N-terminal domain determines variation between MpExo70s. **(A)** Superposition of AlphaFold2 (*77, 79*) predicted structures for MpExo70I (green) and MpExo70II (magenta) indicating the N-terminal, middle and C-terminal domains that are swapped to generate chimera 1, chimera 2 and chimera 3, respectively. Both molecules were represented as cartoon ribbons using ChimeraX (*102*). **(B)** Confocal micrographs of Marchantia cells expressing C-terminally Clover tagged chimeric MpExo70 proteins swapping domains between MpExo70I and MpExo70II:Clover (green), and stained with FM4-64 (magenta). The presence of the cell plate is depicted by accumulation of FM4-64 stain. Scale bar is 10 µm. BF indicates Bright Field. Two independent transformants were analyzed for each line, except for MpExo70I Chimera 2 and MpExo70II Chimera 2 for which only a single transformant was obtained. A gallery of images was collected to document variability in each line and are presented in **Figure S21** and **Figure S22** for MpExo70I and MpExo70II chimeras, respectively. **(C)** Enrichment of proteins co-purified with MpExo70I chimera 1 and MpExo70II chimera 1 compared to a control expressing free GFP and represented by a volcano plot. The horizontal dashed line indicates the threshold above which proteins are significantly enriched (p value < 0.01, quasi-likelihood negative binomial generalized log-linear model) and the vertical dashed line the threshold for which proteins log2 fold change is above 1. For each plot, members of Marchantia exocyst complex are depicted by a red dot with the corresponding name. Wild-type MpExo70I and MpExo70II were including as controls, obtaining a result similar to what is presented in Figure 1b. Asterisk next to the name has been added as a cautionary mark indicating challenges in assigning the peptides to a chimera or a wild-type Exo70. **(D)** Yeast two-hybrid assay of MpExo70I chimera 1 and MpExo70II chimera 1 interaction with MpExo84. For each combination, 5µl of yeast at indicated OD600 were spotted and incubated in double dropout plate for yeast growth control (left) and triple dropout media supplemented with X-α-gal (right). Growth and development of blue coloration in the right panel indicates protein-protein interactions. Wild-type MpExo70I and MpExo70II were included as controls. MpExo70s were fused to the GAL4 DNA binding domain while MpExo84 was fused to the GAL4 activator domain. Accumulation of each protein in yeast cells was tested by western blot and presented in **Figure S28**. **(E)** Co-immunoprecipitation of MpExo70I Chimera 1 and MpExo70II Chimera 1 with MpExo84. C-terminally mScarlet-tagged MpExo70 proteins were stably co-expressed with C-terminally Clover-tagged MpExo84 in Marchantia. Immunoprecipitates (IPs) were obtained with anti-GFP magnetic beads and total protein extracts were probed with anti-RFP or anti-GFP antibodies for MpExo70 and MpExo84 proteins, respectively. Wild-type MpExo70I and MpExo70II were included as controls. Dashed lines represent a crop and assembled image from the same blot. **(F)** Macroscopic phenotypes of Marchantia transgenic lines XVE:MpExo70I1-70:Clover and XVE:MpExo70II1-83 grown for 14 days on media containing B-estradiol or DMSO. Images are representative of experimental replicates presented in **Figure S31**.

To determine whether changes in localization coincided with functional shifts, we investigated how N-terminal exchange altered the interactomes of MpExo70I and MpExo70II using IP-MS. Consistent with our live cell imaging results, IP-MS experiments showed that swapping the N-terminal helical bundle altered MpExo70I and MpExo70II interactomes **(Figure S24, Figure S25, Supplemental dataset 3)**. Notably, swapping the N-terminal domain between MpExo70I and MpExo70II was sufficient to exchange their association with the exocyst complex **(Figure 2c, Figure S26, Figure S27)**, correlating with the change in cell plate localization. To determine whether the localization and interactome changes were linked to MpExo84 interaction, we also assessed Exo70 chimera and Exo84 interactions using Y2H, AlphaFold Multimer and CoIP assays. These experiments demonstrated that the N-terminal region of MpExo70I facilitates interactions and association with MpExo84 while, on the other hand, the MpExo70II N-terminus does not interact with MpExo84 in Y2H and reduces the association with MpExo84 in planta **(Figure 2d and e, Figure S28, Figure S29)**.

Given the importance of the N-terminal region in mediating the interaction between Exo70 and the exocyst complex, we further narrowed down the determinants for MpExo70 localization to the first predicted a-helix (residues Met1 to Leu70 in MpExo70I and Met1 to Leu83 in MpExo70II) **(Figure S30)**. We then hypothesized that the this α-helix would outcompete native Exo70s, causing dominant negative phenotypes. In agreement with our hypothesis, inducible over-expression of MpExo70I1-70:Clover arrested growth in Marchantia **(Figure 2f, Figure S31)**, mimicking ES2 treatment and MpExo70I KE expression. In contrast, expression of MpExo70II1-83:Clover matched the phenotype of MpExo70II KE, causing only a mild growth defect **(Figure 2f, Figure S31).** Altogether, these results suggest that functional divergence between MpExo70I and MpExo70II was driven by alterations in the N-terminal α-helix, indicating that substitutions in this region may have been sufficient to allow functional diversification of MpExo70II.

### Electrostatic divergence in the Exo70 N-terminus facilitated exocyst dissociation

To understand how variation in the N-terminus was able to isolate the Exo70 paralogs and how these changes have evolved, we sought to investigate how N-terminal diversity defines different Exo70 functions. To this end, we first identified Exo70 homologs from diverse land plants and assessed the amino acid conservation in Exo70I, Exo70II, and Exo70III paralogs. In agreement with our experimental results, the N-terminal domain of Exo70II was significantly more diverse than the rest of the protein and both Exo70I and Exo70III **(Figure 3a)**. Given this diversity, we attempted to determine whether amino acid content could differentiate the Exo70 paralogs using linear discriminant analysis **(Figure 3b)**. We found that amino acid content accurately predicted ortholog identity (accuracy ≈ 95%), largely due to an increased abundance of negatively charged residues (D, E) and a decrease in positively charged residues (K, R) in the N-terminal domain of Exo70II compared to Exo70I and Exo70III **(Figure 3c)**. Accordingly, the average net charge of the Exo70II N-terminal domain is more negative than other Exo70s **(Figure 3d)**. Notably, we observed a similar phenomenon in Exo84, where the N-terminal domain of Exo84c is more negatively charged relative to the N-termini of Exo84a and Exo84b. These results suggest that electrostatic tuning could be sufficient to impact the interactions between different Exo70 and Exo84 paralogs **(Figure S32a, b)**.

**Fig. 3.**
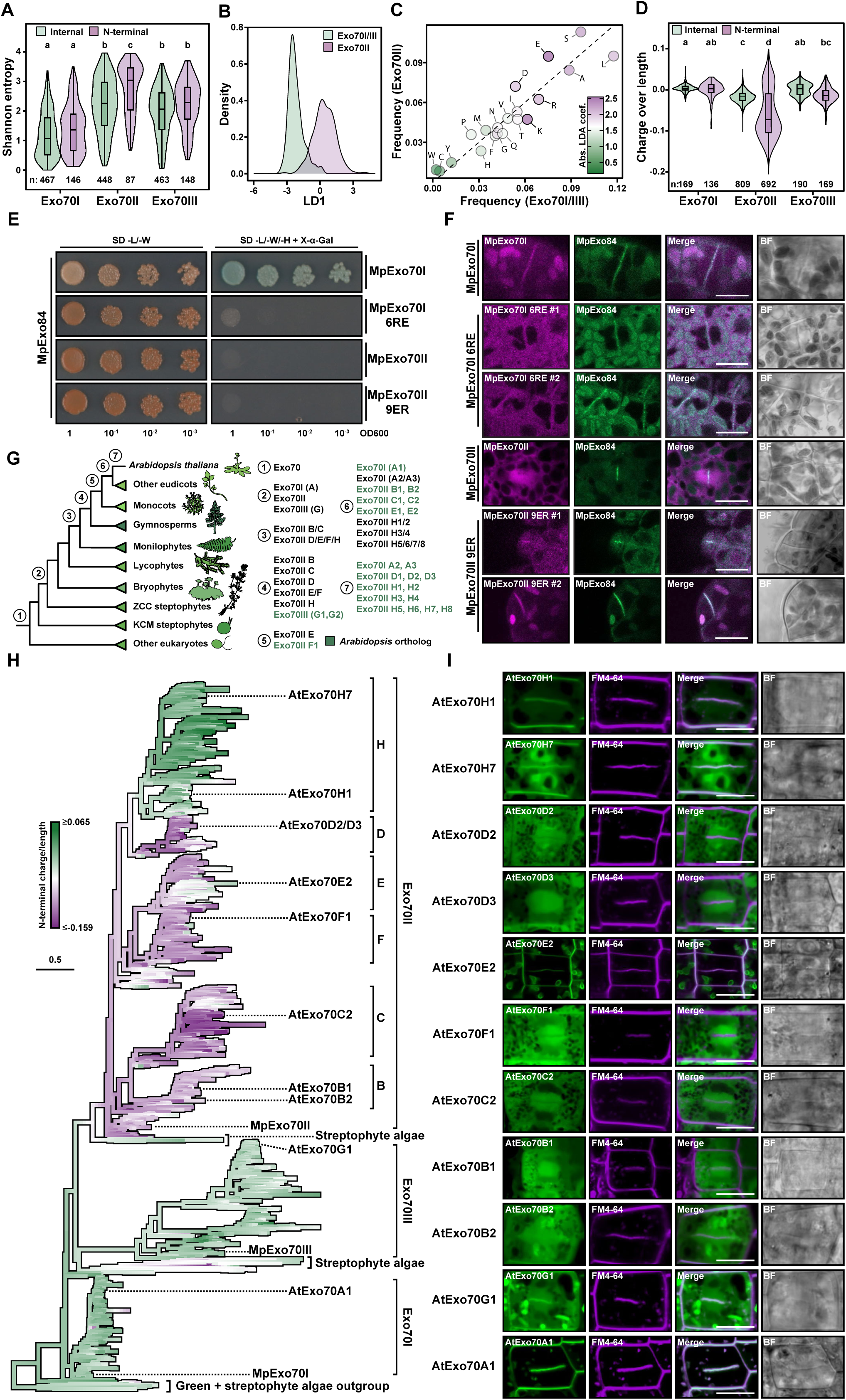
Electrostatic charges underpin functional differences in MpExo70. **(A)** Shannon entropy for aligned sites (*n*) in the N-terminal helix and the remainder of the protein (internal) of land pant Exo70I, Exo70II, and Exo70III paralogs. The center line of the boxplots denotes the median and the upper and lower borders span from the first to the third quartiles, with whiskers extending 1.5 times the interquartile range. Distributions were compared using pairwise Tukey honestly significant difference (HSD) tests. Significance groups are denoted using compact letter display (P < 0.01 after Bonferroni multiple test correction). **(B)** Linear discriminant analysis (LDA) differentiating Exo70 paralogs based on N-terminal α-helix amino acid composition. **(C)** Amino acid frequencies in the N-terminal α-helices of Exo70 paralogs. Points have been colored based on their contribution to the LDA. **(D)** Normalized electrostatic charge of the N-terminal α-helix compared to the rest of the protein for land plant Exo70 paralogs. The number of analyzed proteins is noted (*n*). **(E)** Yeast two-hybrid assay of MpExo70I 6RE and MpExo70II 9ER interaction with MpExo84. For each combination, 5µl of yeast at indicated OD600 were spotted and incubated in double dropout plate for yeast growth control (left) and triple dropout media supplemented with X-α-gal (right). Growth and development of blue coloration in the right panel indicates protein-protein interactions. Wild-type MpExo70I and MpExo70II were included as controls. MpExo70s were fused to the GAL4 DNA binding domain while MpExo84 was fused to the GAL4 activator domain. Accumulation of each protein in yeast cells was tested by western blot and presented in **Figure S33**. **(F)** Confocal micrographs of Marchantia cells from two independent cell lines stably co-expressing C-terminally tagged mScarlet MpExo70I 6RE or MpExo70II 9ER (magenta) with MpExo84:Clover (green). The presence of the cell plate is depicted by accumulation of MpExo84. Wild-type MpExo70I and MpExo70II are included as controls. Scale bar is 10 µm. BF indicates Bright Field. For each line, a gallery of images was collected to document variability and are presented in **Figure S34** and **Figure S35** for MpExo70I 6RE and MpExo70II 9ER, respectively. **(G)** Relative timing of the emergence of each of the *Arabidopsis thaliana* Exo70 paralogs inferred using the Exo70 phylogeny in **Figure S37**. Taxonomic groups are noted with cartoons obtained from Phylopic.org. **(H)** A maximum likelihood phylogeny of the Exo70 family overlaid with ancestral state reconstructions of the normalized electrostatic charge of the N-terminal α-helices. Arabidopsis thaliana and Marchantia polymorpha paralogs used in this study, as well as Exo70 subfamilies, have been noted. The scale bar represents the average number of substitutions per site. Full phylogenies are available in **Figure S37** and from iTOL (*96*) (https://itol.embl.de/shared/OK75j4e8edHZ). **(I)** Confocal micrographs of Arabidopsis root cells stably expressing C-terminally GFP tagged AtExo70 proteins (green) stained with FM4-64 (magenta). The presence of the cell plate is depicted by accumulation of FM4-64 stain. Scale bar is 10 µm. BF indicates Bright Field. Two additional images were collected to document variability in each line and are presented in **Figure S38**.

To experimentally test whether the N-terminal charge defines the association of Exo70I and Exo70II with the exocyst complex, we generated charge substitution mutants in which we replaced MpExo70I residues Arg13, Arg18, Arg41, Arg50, Arg55 and Arg60 with Glu (MpExo70I 6RE), recreating the charge of MpExo70II. Likewise, we neutralized the negative net charge of MpExo70II by replacing residues Glu7, Glu25, Glu30, Glu40, Glu47, Glu54, Glu60, Glu64 and Glu66 with Arg (MpExo70II 9ER). To assess the functional impact of charge inversion, we tested MpExo84 interaction and cell plate localization. Charge inversion in MpExo70I was sufficient to abrogate the interaction with MpExo84 **(Figure 3e, Figure S33)**. Consistently, MpExo70I 6RE lost cell plate localization, adopting a diffuse localization, reminiscent of MpExo70II **(Figure 3f, Figure S34**). In the case of MpExo70II 9ER, charge reversion was not sufficient for MpExo84 interaction **(Figure 3e, Figure S33)** however, it was sufficient for partial re-localization of MpExo70II 9ER to the cell plate **(Figure 3f, Figure S35)**. To ensure these functional changes in MpExo70I 6RE are underpinned by charge reversion rather than alteration of specific residues, we also mutated MpExo70I Arg residues to Ala (MpExo70I 6RA) or Lys (MpExo70I 6RK). In contrast with MpExo70I 6RE, both MpExo70I 6RA and MpExo70I 6RK retained MpExo84 binding **(Figure S36a, b)** and cell plate localization **(Figure S36c)**. Taken together, our results suggest that electrostatic change in the Exo70 N-terminal domain, mediated by few amino acid substitutions, underpins the differential association of Exo70 paralogs with the exocyst complex, highlighting how paralog interference could be mitigated through electrostatic alterations.

### An electrostatic shift at the N-terminal domain predates the expansion of Exo70II

Given our results, we then hypothesized that circumventing paralog interference through complex dissociation mediated by electrostatic changes could have facilitated Exo70 diversification and evolution. To test our hypothesis, we first conducted a comprehensive phylogenetic analysis of the Exo70 family in plants to identify the relationships between Exo70 subfamilies (**Figure S37**). As suggested previously (*38, 40*), the three Exo70 families (I, II, and III) arose early in the streptophyte lineage and, in contrast to Exo70I and III, Exo70II radiated extensively in vascular plants (**Figure 3g, Figure S37, Supplemental dataset 4**). Based on this phylogenetic context, we next sought to examine the evolutionary history of Exo70 N-terminal charge diversification using ancestral state reconstruction. This analysis revealed that the ancestral Exo70 likely harbored an uncharged N-terminus, which was retained in the Exo70I and Exo70III families. However, the emergence of Exo70II coincided with an electrostatic shift towards negative charge in the N-terminal region which preceded the Exo70 radiation in vascular plants **(Figure 3h)**. Interestingly, during the diversification of Exo70II, N-termini in subclades E and H seem to have reverted to a neutral or positive overall charge. Exo84 showed a similar pattern where N-terminal charge diversification coincided with Exo84 duplication early in land plant evolution **(Figure S32c, d)**. Based on these data, we hypothesized that dissociation of Exo70II from the exocyst underpinned by alteration of N-terminal charge, allowed for the expansion and subsequent functional diversification of these proteins. To experimentally link N-terminal charge with exocyst association across the phylogenetic tree, we generated a collection of Arabidopsis lines overexpressing representative members of Exo70 subclades A to H and assessed their localization at the cell plate **(Figure 3i, Figure S38)**. Similar to Marchantia, Arabidopsis Exo70s from clades I and III with positively charged N-terminal domains (Subclade A and G, respectively), localized at the cell plate. On the contrary, Exo70II paralogs with a predicted negative charge (Subclades B, C, D and F) had a diffuse localization reminiscent of Marchantia MpExo70II. However, Exo70II paralogs exhibiting charge reversion such as AtExo70E2, AtExo70H1 and AtExo70H7, all localized to the cell plate **(Figure 3i, Figure S38**). Overall, our data indicate that electrostatic tuning played a crucial role in Exo70 evolution, providing a straightforward mechanism for dissociation from the exocyst complex. Exo70II escape from the exocyst complex could have alleviated the evolutionary constraints of being in a multimeric protein assembly and prevented paralog interference, allowing rapid diversification and neofunctionalization.

## Discussion and conclusion

Given that most of the protein machinery governing essential cellular processes was already present in the Last Common Eukaryotic Ancestor (*60*), deciphering the mechanisms underlying their diversification throughout eukaryotic evolution remains a fundamental question in evolutionary cell biology. In this study, we reveal how a core subunit of the exocyst complex, Exo70, diversified in plants through a process of dissociation from the exocyst complex. This separation was driven by electrostatic changes in the N-terminal region of Exo70, which are crucial for its interaction with Exo84 and subsequent association with the exocyst. This disengagement from the complex alleviated evolutionary constraints, particularly those imposed by paralog interference, ultimately facilitating a significant expansion and diversification of Exo70 in vascular plants **(Figure 4a)**.

**Fig. 4.**
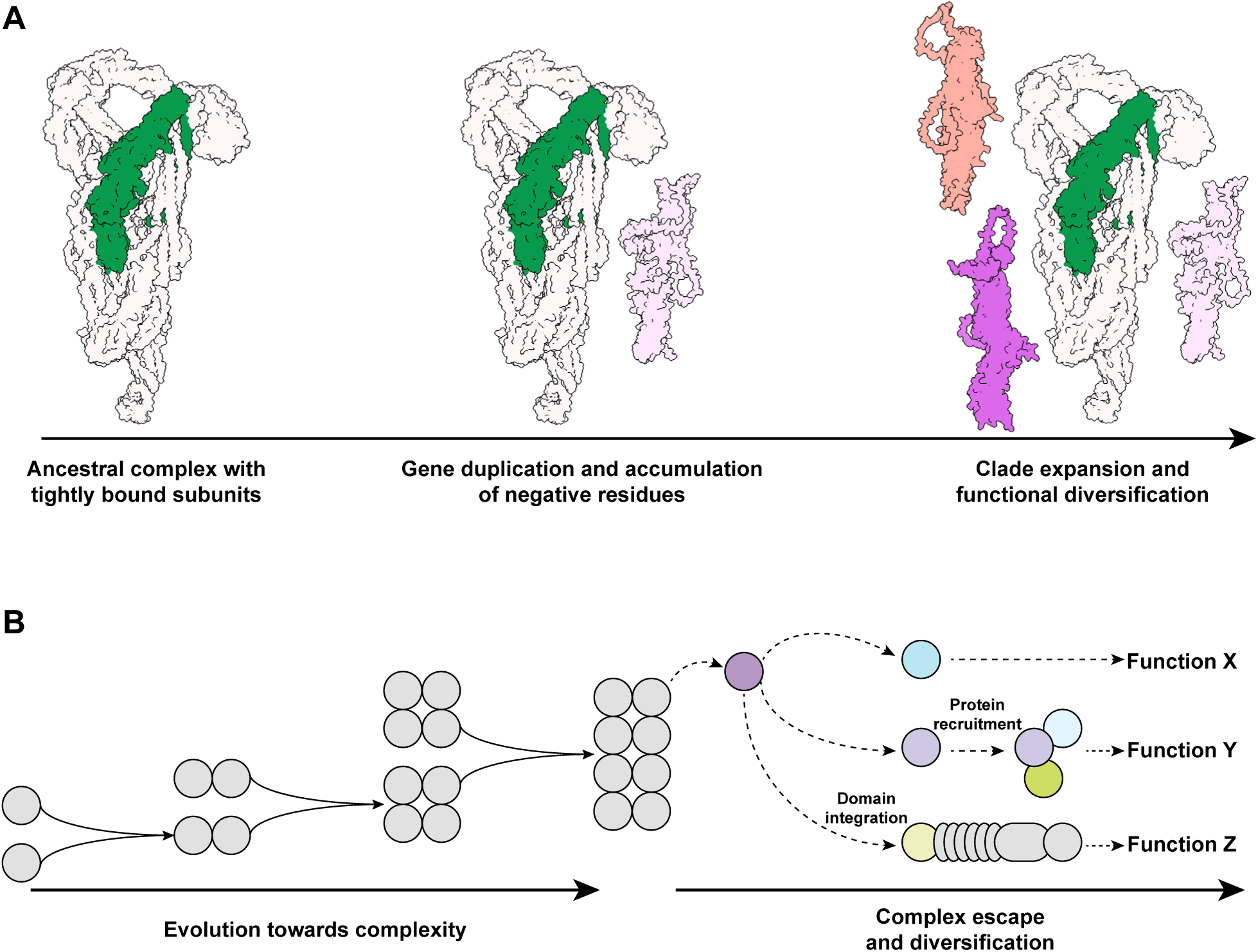
Graphical summary of plant Exo70 diversification by protein complex escape. **(A) From left to right:** The exocyst is an heteromeric complex composed by tightly bound subunits required to perform its function. Gene duplication and accumulation of negative residues at the N-terminal domain led to dissociation of the Exo70 subunit. Dissociation from the exocyst complex relieved paralog interference and allowed for Exo70 expansion and functional diversification. **(B)** Protein complexes can evolve towards complexity by gene duplication and neofunctionalization. In a protein complex escape scenario, a single subunit dissociates from the complex to perform a novel function (function X), to be recruited by molecular interactions with other proteins (function Y) or to get integrated as a domain within other proteins (function Z).

However, complex dissociation alone does not fully account for the evolutionary trajectory of Exo70. Escape of Exo70II from the exocyst complex opened avenues for novel interactions, allowing these paralogs to integrate into other cellular processes. Supporting this hypothesis, MpExo70I Chimera1 exhibited reduced association with the exocyst complex in our IP-MS analysis and, in turn, developed a distinct interactome separate from MpExo70II. We propose that, over evolutionary timescales, the novel interactions that emerged following Exo70II dissociation resulted in the recruitment of Exo70 into diverse pathways unrelated to exocytosis. For instance, Exo70s have been reported to acquire short linear motifs that mediate interactions with the core autophagy protein ATG8 (*61, 62*), linking them to autophagy (*63*), particularly within the Exo70II clades B (*48, 49*) and D (*50*). Specifically, AtExo70D paralogs have been shown to act as cargo receptors, mediating the degradation of type-A ARRs proteins and thereby modulating cytokinin signaling (*50*). Another example of neofunctionalization is observed in the role of certain Exo70 paralogs in plant immunity (*39*). In particular, Exo70II paralogs in clade F are involved in pathogen recognition in monocots (*51, 52, 64*) and have even been incorporated as integrated domains within intracellular immune receptors (*53, 54*) to detect pathogens (*65*). None of these functions are associated with exocytosis, making it difficult to envisage a role for the full exocyst complex in these processes. Our findings suggest that the most parsimonious explanation for these activities is that Exo70II proteins function independently of the exocyst. Consistent with this, some members of the Exo70 F clade have lost the entire N-terminal domain, likely precluding their participation in forming the CorEx and associating with the exocyst (*52*). Another significant implication of our findings concerns the accuracy of protein function predictions based on domain analysis. Our results, along with previous studies, indicate that Exo70II paralogs may function independently of the exocyst complex. However, homology-based functional predictions often erroneously assume canonical Exo70 functions for these paralogs. This suggests that current predictive tools may underestimate the functional diversity of proteins predicted to be embedded within multimeric assemblies.

Previous studies, primarily focused on homomeric complexes, have elucidated how proteins evolve into multimeric assemblies (*4, 8, 10, 25*) through mechanisms such as the formation of hydrophobic interfaces (*6, 8, 9*) and processes leading to molecular entrenchment (*5, 7, 10*). These mechanisms typically constrain individual subunits from gaining functional independence, while paralog interference further limits their evolutionary diversification (*15, 17*). In contrast, our research uncovers a simple yet effective mechanism by which polymorphisms in a single subunit of a heteromeric complex can bypass both molecular entrenchment and paralog interference. This allows the subunit to dissociate from the original complex and be co-opted into alternative cellular pathways **(Figure 4b)**. The extent to which this ‘protein complex escape’ phenomenon occurs in other complexes likely depends on specific factors, such as the ease of subunit dissociation, the potential to form new interactions, and the ability to function as an independent module. We anticipate that further research will uncover overlooked cases of subunit duplication in other complexes, revealing unexpected instances of functional diversification and contributing to a deeper understanding of cellular complexity in eukaryotic evolution.

## Acknowledgements

We thank Dr. David Mackey and Dr. Roger W. Innes for providing plasmids that served as template to clone Arabidopsis Exo70s, Dr. Khong-Sam Chia and Dr. Philip Carella for providing the plasmid pMpGWB168 (XVE::GW) and Dr. Niko Geldner for providing the Arabidopsis exo70a1/CASP1:GFP line. We also thank Dr. Yan Ma for assistance on imaging Arabidopsis Casparian Strips and Dr. Hannes Maib for providing the initial biosensor reagent. This work was supported by the BioOptics facility, The Plant Sciences Facility, the Protein Technologies Facility and the Mass Spectrometry Facility of the Vienna BioCenter Core Facilities. We gratefully acknowledge the Dundee Imaging Facility and the University of Dundee for their support in this work. Thanks to Dr. Gautam Dey, Dr. Chung Hyun Cho and Prof. Sophien Kamoun for critical reading of the manuscript.

## Author contributions

Conceptualization: J.C., N.A.T.I. and Y.D. Methodology: J.C., H.D., Y.K., J.J., H.D.X., S.A.I., A.B., N.G, I.G., D.H.M., M.C., N.A.T.I. Investigation: J.C., H.D., Y.K., J.J., H.D.X.,S.A.I., A.B., D.H.M., M.C., N.A.T.I. Visualization: J.C., H.D., Y.K., J.J., H.D.X., D.H.M., M.C., N.A.T.I. Funding acquisition: D.H., D.H.M., N.A.T.I., Y.D. Project administration: Y.D. Supervision: J.C., D.H., D.H.M., Y.D. Writing – original draft: J.C., N.A.T.I., Y.D. Writing – review & editing: J.C., H.D., Y.K., J.J., H.D.X., S.A.I., A.B., N.G, R.P., I.G., D.H., D.H.M., M.C., N.A.T.I., Y.D.

## Data and materials availability

Source data and raw images used to generate the main and supplementary figures are deposited to Zenodo (https://zenodo.org/records/13374123). The mass spectrometry proteomics data have been deposited to the ProteomeXchange Consortium via the PRIDE partner repository (*66*) with the dataset identifier PXD051853.

## Materials and Methods

### Molecular cloning

For protein production in *E. coli*, coding sequences for MpExo70I and MpExo70I 6KE were synthesized as double-stranded DNA fragments (Twist Bioscience). DNA fragments spanning amino acids 70 to 649 were amplified by PCR and subsequently cloned into the pOPIN-GG vector pPGN-C (*67*) with an N-terminal 6xHis:SUMO tag via Golden Gate cloning (*68*). For protein production in insect cells, GFP:MpExo70I99-649 and GFP:MpExo70II111-685 were synthesized as codon-optimized DNA fragments and subsequently cloned in a baculovirus expression vector system (BEVS), GoldenBac (*69*), by the ProTech service at Vienna BioCenter. For production of Marchantia exocyst subcomplex IIa (MpSEC15:MpSEC10:MpExo84:MpExo70I) and exocyst subcomplex IIb (MpSEC15:MpSEC10:MpExo84:MpExo70II), codon-optimized full-length coding sequences were synthesized and subsequently cloned for co-expression in BEVS by GoldenBac.

For Y2H assays, coding sequences for MpExo70 proteins and mutants, as well as MpExo84, were synthesized as double-stranded DNA fragments (Twist Bioscience). The fragments were subsequently inserted via BsaI in a pGADT7 (MpExo84) or pGBKT7 (MpExo70) vectors adapted for Golden Gate cloning (*68*) provided by the SynBio service at The Sainsbury Laboratory, Norwich.

Plasmids for stable transformation in *Marchantia polymorpha* and *Arabidopsis thaliana*, coding sequences were assembled with the indicated promoter, fluorescence tag and terminators into pGGsun via BsaI-mediated GreenGate cloning (*70*). MpExo70I:Clover, MpExo70I 6KE:Clover, MpExo70I1-70:Clover, MpExo70II:Clover, MpExo70II 6KE:Clover and MpExo70II1-83:Clover β-estradiol inducible expression constructs were generated by Gateway cloning (*71*) into pMpGWB168 (XVE::GW) (*72, 73*).

### Plant growth and transformation

#### Marchantia polymorpha

*Marchantia polymorpha* accession Takaragaike-1 (Tak-1) were axenically maintained and asexually propagated on 0.5 Gamborg B5 + MES medium (1.5 g/L B5 Gamborg, 0.5 g/L MES hydrate, 1% sucrose, pH adjusted to 5.5) containing 1% (w/v) agar at 21 °C under continuous white light (50 µM m-² s-1).

Stable transformants of Marchantia were generated by gemmae transformation. Tak-1 gemmae were grown for two days in 0.5 Gamborg B5 + MES medium plates containing 1% (w/v) agar. A solution of *Agrobacterium tumefaciens* strain GV3101 (C58 [RifR] Ti pMP90 [pTiC58DT-DNA] [gentR] Nopaline [pSoup-tetR]) containing the desired construct resuspended in transformation media (1.5 g/L B5 Gamborg, 0.5 g/L MES hydrate, 1 g/L casein hydrolysate, 0.3 g/L of L-glutamine, 2% sucrose, pH adjusted to 5.5) supplemented with 150 µM acetosyringone was added to the plates and incubated at 28 °C in dark. After incubation, Marchantia gemmae were washed with sterile water, scrapped out from the transformation plate and transfer to 0.5 Gamborg B5 + MES plates containing Ticarcillin and the selective antibiotic. Transformant were recovered from the plates after 2-4 weeks and protein expression was assessed by western blot. A list of Marchantia lines generated here can be found in **Table S1**.

#### Arabidopsis thaliana

*Arabidopsis thaliana* lines used in this study were sown on water-saturated soil for standard plant growth under a 16 h light/8 h dark photoperiod with 165 µM m-² s-1 light intensity. For in vitro seedling growth, Arabidopsis seeds were surface sterilized in 70% ethanol 0.05% SDS for 15 minutes, rinsed in ethanol absolute and dried on sterile paper. Seeds were plated in ½ MS salts (Duchefa), 1% sucrose containing 1% (w/v) agar and stratified for 48 hours at 4°C in dark. Plates were then grown under LEDs with 50 µM m-2 s-1 and a 16 h light/8 h dark photoperiod.

Stable transformants of Arabidopsis thaliana were generated by delivering the desired constructs via plant transformation with *Agrobacterium tumefaciens* strain GV3101 (C58 [RifR] Ti pMP90 [pTiC58DT-DNA] [gentR] Nopaline [pSoup-tetR]) using the floral dip method (*74*). A list of Arabidopsis lines generated here can be found in **Table S1**.

### Confocal microscopy

For imaging of *Marchantia polymorpha* cells using confocal microscopy, asexual gemmae were incubated in liquid 0.5 Gamborg B5 + MES medium (1.5 g/L B5 Gamborg, 0.5 g/L MES hydrate, 1% sucrose, pH adjusted to 5.5) for two days at 21 °C under continuous white light (50 µM m-² s-1). The thalli were then placed on a microscope slide with deionized water and covered with a coverslip. The meristem region was used for image acquisition. For co-localization of Marchantia exocyst proteins with FM4-64, plants were treated for 10 min with 13.3 µM FM4-64 prior imaging.

For confocal microscopy of *Arabidopsis thaliana* seedling roots, seed sterilization was performed using Sterilization Solution I (70% ethanol, 0.01% Triton X-100) for 10min followed by Sterilization solution II (50% Sodium hypochlorite) for 5 min. Removal of the sterilization solution II was done rinsing the seeds 4 times with milli-Q water. Seeds were sown on Murashige–Skoog (MS) media right after sterilization. Stratification was conducted incubating at 4 °C in dark for 3 days and after it, the plates were incubated in controlled-environment growth chamber at 22 °C under long day photoperiod conditions (16-h-light/8-h-dark photoperiod) at 80-100 µM m-² s-1. Plates were placed vertically to let the roots elongate along the media surface.

3 days old Arabidopsis seedlings were treated for 5 min with 13.3 µM FM4-64, placed on a microscope slide with water and covered with a coverslip. The epidermal cells of root meristem zone were used for AtExo70 and FM4-64 colocalization imaging. For imaging of the Arabidopsis casparian strip, we imaged the tenth root cell above the elongation region.

Confocal imaging was performed using a downright point laser scanning confocal microscope Zeiss LSM 800 AxioImager.Z2 (Carl Zeiss) equipped with plan-Apochromat 63x corrective water immersion objective and ZEN software (blue edition, Carl Zeiss). FM 4-64 fluorescence was excited at 561 nm and detected between 656 and 700 nm. GFP fluorescence was excited at 488 nm and detected between 488 and 545 nm. mScarlet fluorescence was excited at 561 nm and detected between 570 and 617 nm. Pinholes were adjusted to one Air Unit for each wavelength. For each experiment, all replicate images were acquired using identical confocal microscopic parameters. Confocal images were processed with Fiji (version 1.52, Fiji).

### Chemical treatments in *Marchantia polymorpha*

Chemical treatments to test the effect of Endosisin2 (ES2) (*58*) in plant growth or to induce protein expression using β-estradiol were performed by culturing Marchantia gemmae directly into 0.5 Gamborg B5 + MES medium (1.5 g/L B5 Gamborg, 0.5 g/L MES hydrate, 1% sucrose, pH adjusted to 5.5) containing 1% (w/v) supplemented with different concentrations of ES2 or 20 µM β-estradiol, respectively.

### Expression and purification of proteins for in vitro studies

#### Expression and purification of MpExo70s in *E. coli*

Heterologous expression and purification of SUMO-tagged MpExo70 proteins was performed as previously described (*59*). SUMO:MpExo70I70-649 or SUMO:MpExo70I70-649 KE were produced in *E. coli* RosettaTM (DE3). Cell cultures were grown in terrific broth at 37°C for 5–7 h and then at 16°C overnight after induction with 1 mM IPTG. Cells were harvested by centrifugation and re-suspended in 20 mM HEPES pH 8, 500 mM NaCl, 5% (vol/vol) glycerol, and 20 mM imidazole supplemented with cOmpleteTM EDTA-free protease inhibitor tablets (Roche). Cells were sonicated and, following centrifugation at 40,000xg for 30 min, the clarified lysate was applied to a HisTrapTM Ni2+-NTA column connected to an ÄKTA pure™ protein purification system (Cytiva). Proteins were step-eluted with elution buffer (20 mM HEPES pH 8, 500 mM NaCl, 5% (vol/vol) glycerol, and 500 mM imidazole).

Fraction with the eluted proteins were collected and injected onto a HiLoad® 16/600 Superdex® 200 pg gel filtration column (Cytiva) pre-equilibrated with 20 mM HEPES pH 7.5, 150 mM NaCl and 5% (vol/vol) glycerol supplemented with 1mM Tris 2-carboxyethyl phosphine hydrochloride (TCEP, Sigma). Elution fractions were collected and evaluated by SDS-PAGE and relevant fractions with purified proteins were concentrated as appropriate for further studies.

#### Expression and purification of MpExo70s in insect cells

Expression and production of GFP:MpExo70I99-649 and GFP:MpExo70II111-685 in insect cells, source plasmids containing the constructs were transformed into DH10EMBacY cells. Colonies containing recombinant baculoviral shuttle vectors (bacmids) were selected by blue-white selection on LB agar plates containing X-Gal and IPTG. Bacmid DNA was extracted by alkaline lysis and isopropanol precipitation and confirmed by PCR. To generate the viral stock (V0), bacmids were then transfected into adherent *Spodoptera frugiperda* (Sf9) insect cells in six-well plates, using transfection medium (Expression systems) and PEI. Successful transfection was tracked by monitoring a yellow fluorescent protein encoded by the bacmid backbone. The plate was incubated for 1 week and the virus was harvested for amplification.

Recombinant protein was produced in *Trichoplusia ni* High FiveTM cells (Thermo Fisher) infected at a density of 1× 106 ml with the appropriate virus, and grown at 21 °C and 120 r.p.m. Cells were harvested 4 days after infection by centrifugation at 1000 g for 15 min, and pellets were stored at –70 °C. All insect cell culture works were performed using ESF 921 serum-free growth medium (Expression Systems) without antibiotic supplementation.

Cell pellets were resuspended in 20 mM HEPES pH 8, 300 mM NaCl, 5% (vol/vol) glycerol, and 20 mM imidazole supplemented with cOmpleteTM EDTA-free protease inhibitor tablets (Roche) and Benzonase (IMP Molecular Biology Service). Resuspended cells were then disrupted using a glass Dounce homogenizer. After centrifugation at 40,000 g for 30 min the clarified lysate was processed as described above for protein purification.

#### Expression and purification of Marchantia exocyst subcomplex II

For expression and production of Marchantia exocyst subcomplex IIa (MpSEC15:MpSEC10:MpExo84:MpExo70I) and exocyst subcomplex IIb (MpSEC15:MpSEC10:MpExo84:MpExo70II), source plasmids containing the four proteins were transformed into DH10EMBacY cells. Viral stocks and *Trichoplusia ni* High FiveTM cells expressing the subcomplexes were produced as described above.

Once harvested, the pellets were re-suspended in 20 mM HEPES pH 8, 150 mM NaCl, and 20 mM imidazole supplemented with cOmpleteTM EDTA-free protease inhibitor tablets (Roche) and Benzonase (IMP Molecular Biology Service). Resuspended cells were then disrupted using a glass Dounce homogenizer. After centrifugation at 40,000 g for 30 min the clarified lysate was applied to a HisTrapTM Ni2+-NTA column connected to an ÄKTA pure™ protein purification system (Cytiva). Proteins were step-eluted with elution buffer (20 mM HEPES pH 8, 150 mM NaCl, and 500 mM imidazole).

Fraction with the eluted proteins were collected and injected onto a HiLoad® 16/600 Superose 6 pg preparative gel filtration column (Cytiva) pre-equilibrated with 20 mM HEPES pH 7.5, 150 mM NaCl and supplemented with 1mM TCEP. Elution fractions were collected and evaluated by SDS-PAGE. Relevant fractions with purified proteins were then concentrated as appropriate.

### Analysis of purified proteins by mass photometry

Mass photometry analysis of proteins was performed at room temperature on a OneMP photometer instrument (Refeyn) with the data acquisition software AcquireMP (Refeyn). High precision cover glasses (24 × 50 mm, VWR) were rinsed thoroughly with isopropanol, cleared using double-distilled water, and dried under a clean nitrogen stream. One drop of type F immersion oil (Olympus) was applied to the photometer lens and a cleaned cover glass was settled on the lens. A gasket (Grace Bio-labs) was then adhered to the cover glass and 9 µL of the buffer was pipetted into the gasket wall without touching the cover glass. The buffer drop was focused in a 5 µm × 10 µm field to a stable sharpness value around 5.5 before gently pipetting 1 µL of diluted protein sample (∼0.1 µM) into the drop. The counting events were recorded for 60s at a 1kHz frame rate and images were processed using DiscoverMP (Refeyn).

### Protein-lipid interaction: Membrane overlay assays

Membrane overlay assay were performed with slight variation of a previously published protocol (*46*). In brief, Membrane lipid strips (Echelon Bioscience, P-6002) were blocked with 10 mL blocking buffer (5% bovine serum albumin, 10 mM Tris-HCl pH 7.5, 150 mM NaCl, 0.1% Tween 20) at room temperature for 1 h in gentle agitation. After incubation, 5 ml of the blocking buffer was exchanged with 5 ml of blocking buffer containing ∼130 nM concentration of purified SUMO:MpExo70I70-649 or SUMO:MpExo70I70-649 KE. Strips were then incubated at room temperature for 2 h under gentle agitation and then washed three times for 15 min with TBST (10 mM Tris-HCl pH 7.5, 150 mM NaCl, 0.1% Tween 20). A solution of SUMO-tag Monoclonal Antibody (4G11E9, Thermo Fischer Scientific) at 1:1000 dilution in blocking buffer was added to the lipid strips and incubated 1 h at room temperature. The membranes were then washed with TBST again 3 times for 15 min before adding Goat Anti-Mouse IgG (H + L)-HRP antibody (1706516, Bio-Rad) 1:10000 in blocking buffer. After 1 h incubation, membranes were washed again 3 times with TBST, and images were captured using iBright Imaging System (Invitrogen) with SuperSignal West Pico PLUS Chemiluminescent Substrate (Thermo Fisher Scientific).

### Protein-lipid interaction: Membrane coated beads binding assay

Liposomes and membrane coated beads were generated as described previously (*27*). In brief, liposomes containing PI(4,5)P2 were made by mixing 85 mol % 1-palmitoyl-2-oleoyl-glycero-3-phosphocholine (POPC, Avanti Polar Lipids), 10 mol % phosphatidylserine (POPS, Avanti Polar Lipids), 5 mol % PI(4,5)P2 (Echelon Biosciences), and 0.1% Atto647N-DOPE (ATTO-TEC). Control liposomes without phosphoinositides were made by mixing 90 mol % POPC, 10 mol % POPS, and 0.1% Atto647N-DOPE. The mixtures were then evaporated under clean nitrogen stream and dried overnight under vacuum. Dried lipids were resuspended in 20 mM HEPES, 150 mM NaCl, 0.5 mM Tris 2-carboxyethyl phosphine hydrochloride (TCEP, Sigma) at 37 °C. The mixtures were then freeze-thawed for 6 cycles in liquid nitrogen and extruded to 0.1 µm via track-etched membranes (Whatman). The homogenized liposomes were ready to use, aliquoted, and frozen in liquid nitrogen and stored at - 20 °C.

Membrane coated beads were made in 100 µL reactions by incubating 100 µM liposomes and 5 × 105 of silica beads (Whitehouse Scientific) in 250 mM NaCl on a rotator at room temperature for 30 min. Beads were then washed twice with 20 mM HEPES, pH 7.4, and resuspended in 20 mM HEPES, pH 7.4, 250 mM NaCl, 0.5 mM TCEP. Membrane coated beads were kept rotating while the binding assay was set up.

#### Protein-membrane binding assay was set up by mixing 7 µL of the membrane coated beads and 7 µL of protein samples in gradient concentrations in uncoated µ-Slide 8 well chambers (Ibidi)

### Protein-protein interaction: Yeast-2-hybrid assay

Yeast-two-hybrid assays of the interaction between MpExo84 and MpExo70 alleles and mutants were adapted from a previously described protocol (*59, 75*). In brief, a pGADT7 plasmid encoding MpExo84 was co-transformed into chemically competent Y2HGold cells (Takara Bio, USA) with a pGBKT7 plasmid encoding MpExo70 alleles and mutants using the Frozen-EZ Yeast Transformation Kit (Zymo research).

After growing in selection plates, single co-transformants were inoculated in liquid SD -Leu -Trp media for two days at 30 °C. Saturated culture was then used to make serial dilutions of OD600 1, 0.1, 0.01, and 0.001 and 5 µl of each dilution was spotted on a SD -Leu -Trp plate as a growth control, and on a SD -Leu -Trp -His plate containing X-α-gal (Takara Bio, USA). Plates were imaged after incubation for 72 h at 30 °C. Each experiment was repeated a minimum of three times, with similar results.

To assay the accumulation of MpExo84 and MpExo70 proteins in yeast cells, total yeast extracts were produced by harvesting cells from the liquid media and incubation for 10 min at 95°C after resuspending them in Laemmli buffer. Samples were then centrifugated and the supernatant was subjected to SDS-PAGE and western blot. The membranes were probed with anti-GAL4 DNA Binding domain (Sigma) antibody for the MpExo70s proteins in pGBKT7 and with the anti-GAL4 activation domain (Sigma) antibody for MpExo84 in pGADT7.

### Protein-protein interaction: In planta Co-immunoprecipitation

For protein extraction, Marchantia gemmae from 10 gemmae cups were placed in 40 ml of liquid 0.5 Gamborg B5 + MES medium and incubated at 21 °C under continuous white light (50 µM m-² s-1). Plant tissue was then collected and briefly dried on paper before grind to fine powder in liquid nitrogen using a pestle and mortar. Plant powder was mixed with two times weight/volume ice-cold extraction buffer (10% glycerol, 25 mM Tris pH 7.5, 1 mM EDTA, 150 mM NaCl, 2% w/v PVPP, 10 mM DTT, 1× tablet of cOmplete™, EDTA-free Protease Inhibitor Cocktail [Roche], 0.1% Tween 20 [Sigma] per 50 ml), centrifuged at 4200 × g at 4°C for 20–30 min, and the supernatant was passed through a 0.45 µm Minisart syringe filter. The presence of proteins in the input was determined by SDS-PAGE/western blot and probing the membranes with anti-GFP (11814460001/54732800, Roche) or anti-RFP (AB_2631395, Chromotek) antibodies for protein tagged with Clover and mScarlet, respectively.

For immunoprecipitation, 1 ml of filtered plant extract was incubated with 30 µl of GFP-Trap® Magnetic beads (Chromotek) in a rotatory mixer at 4 °C. After 1 h, the beads were pelleted on a magnetic rack and the supernatant removed. The pellet was then washed and resuspended in 1 ml of IP buffer (10% glycerol, 25 mM Tris pH 7.5, 1 mM EDTA, 150 mM NaCl, 0.1% Tween 20 [Sigma]) and pelleted again as before. Washing steps were repeated five times. Finally, 30 µl of Laemmli buffer was added to the agarose and incubated for 10 min at 70°C. The beads were pelleted again by centrifugation, and the supernatant loaded on SDS-PAGE gels prior to western blotting. Membranes were probed with anti-GFP antibody (11814460001/54732800, Roche) to detect MpExo84:Clover and anti-RFP (AB_2631395, Chromotek) to detect MpExo70:mScarlet proteins.

### Protein-protein interaction: Affinity purification-mass spectrometry

#### Affinity purification

For affinity purification of Marchantia exocyst proteins, stable lines overexpressing MpSEC6, MpExo84 or MpExo70s with a C-terminal Clover tag were grown and processed for total protein extraction as detailed above.

Immunoprecipitation of the proteins was performed by adding 30 µl of GFP-Trap® Magnetic beads (Chromotek) into 1 ml of crude extract and incubating in LoBind® Eppendorf tubes for 1 h in a rotatory mixer at 4 °C. The beads were washed twice with IP buffer (10% glycerol, 25 mM Tris pH 7.5, 1 mM EDTA, 150 mM NaCl, 0.1% Tween 20 [Sigma]) by pelleting them on a magnetic rack and removing the supernatant. The beads were then washed four more times with IP buffer without Tween 20. After washing, the magnetic beads were resuspended in 100 µl of IP buffer without Tween 20. 10 µl was drained and denatured in 20 µl of Laemmli buffer for quality check by western blot. The remaining 90 µl was pelleted by centrifugation and, after removing the excess of buffer, the tubes were stored at -20 °C before on-bead Trypsin digestion.

#### Trypsin digestion

Beads were resuspended in 40µl of 100 mM ammonium bicarbonate (ABC), supplemented with 400 ng of lysyl endopeptidase (Lys-C, Fujifilm Wako Pure Chemical Corporation) and incubated for 4 h on a Thermo-shaker with 1200 rpm at 37°C. The supernatant was transferred to a fresh tube and reduced with 0.5mM Tris 2-carboxyethyl phosphine hydrochloride (TCEP, Sigma) for 30 min at 60°C and alkylated in 4 mM methyl methanethiosulfonate (MMTS, Fluka) for 30 min at room temperature. Subsequently, the sample was digested with 400 ng trypsin (Trypsin Gold, Promega) at 37°C overnight. The digest was acidified by addition of trifluoroacetic acid (TFA, Pierce) to 1%. A similar aliquot of each sample was analyzed by LC-MS/MS.

#### Mass spectrometry data acquisition

The nano HPLC system (UltiMate 3000 RSLC nano system) was coupled to an Orbitrap ExplorisTM 480 mass spectrometer equipped with a Nanospray Flex ion source (all parts Thermo Fisher Scientific).

Peptides were loaded onto a trap column (PepMap Acclaim C18, 5 mm × 300 µm ID, 5 µm particles, 100 Å pore size, Thermo Fisher Scientific) at a flow rate of 25 µl/min using 0.1% TFA as mobile phase. After loading, the trap column was switched in line with the analytical column (PepMap Acclaim C18, 500 mm × 75 µm ID, 2 µm, 100 Å, Thermo Fisher Scientific). Peptides were eluted using a flow rate of 230 nl/min, starting with the mobile phases 98% A (0.1% formic acid in water) and 2% B (80% acetonitrile, 0.1% formic acid) and linearly increasing to 35% B over the next 120 min. This was followed by a steep gradient to 95%B in 5 min, stayed there for 5 min and ramped down in 2 min to the starting conditions of 98% A and 2% B for equilibration at 30°C.

The Orbitrap Exploris 480 mass spectrometer was operated in data-dependent mode, performing a full scan (m/z range 350-1200, resolution 60,000, normalized AGC target 300%) at 3 different compensation voltages (CV -45 V, -60 V and -75 V), followed by MS/MS scans of the most abundant ions for a cycle time of 0.9 s for CVs -45 V and - 60 V, and 0.7 s for CV -75 V. MS/MS spectra were acquired using an isolation width of 1.2 m/z, normalized AGC target 200%, minimum intensity set to 25,000, HCD collision energy of 30 %, maximum injection time of 100 ms and resolution of 30,000. Precursor ions selected for fragmentation (include charge state 2-6) were excluded for 45 s. The monoisotopic precursor selection (MIPS) mode was set to peptide and the exclude isotopes feature was enabled.

#### Analysis of mass spectrometry results

The total number of MS/MS fragmentation spectra was used to quantify each protein (Tables S21 to S27). The data matrix of Peptide-Spectrum Match (PSM) was analyzed using the R package IPinquiry4 (https://github.com/hzuber67/IPinquiry4) that calculates log2 fold change and P values using the quasi-likelihood negative binomial generalized loglinear model implemented in the edgeR package (*76*). Only proteins identified with at least 3 PSM were considered. Each sample was triplicated per experiment. Candidate proteins for each bait were filtered by pairwise comparison to GFP (empty vector) control. For Figure S12, pairwise comparison between the bait of interest and MpExo70II was performed. For all pairwise comparisons performed for WT exocyst subunits threshold for candidate selection was set at log2 fold change>0 and pvalue<0.05. For the experiment performed with MpExo70 chimeras, threshold for candidate selection was set at log2 fold change>1 and pvalue<0.01. Annotations were retrieved for each protein detected in both experiments using the ID mapping web tool of UniProt (https://www.uniprot.org/id-mapping, access December 2023). Venn diagram was built using the Venny 2.1.0 online tool and then redrawn manually. Protein abundance was plotted in a heatmap as log2(meanPSM+1) and normalized to the mean value per protein using R. Rows and columns were clustered using Euclidean distance.

### Protein-protein interaction: AlphaFold2 Multimer

We used AlphaFold2 Multimer (*77–79*) to predict protein-protein interaction between Exo70s and candidate proteins as described in (*80, 81*). Protein sequences were extracted in FASTA format from marchantia.info and processed using mmseqs (*82*) to generate local multiple sequence alignments. Subsequent structural predictions were performed with ColabFold (*79*). The confidence in protein-protein interaction predictions was assessed using two metrics: the Interface Predicted TM-score (ipTM) and a custom PEAK score. The PEAK score represents the Predicted Aligned Error (PAE) between chains, excluding intra-molecular interactions. For each interaction, five independent models were generated by Alphafold2 using default settings. The average or maximum ipTM and PEAK scores from these models were used in the analysis.

### Bioinformatic analyses: Comparative genomics and phylogenetics

Exo70 and Exo84 paralogs were identified across eukaryotes using a diverse eukaryotic dataset comprised of genome-predicted proteomes obtained from Uniprot (n = 174, downloaded 15 March 2023) (*83*). To taxonomically balance the dataset, we selected the best two proteomes per genus based on BUSCO (Benchmarking Universal Single Copy Orthologues) completeness (*84*). For metazoans, fungi, and embryophytes, more strict taxonomic criteria were set by selecting the best proteome per phyla for metazoans (n = 17), the best proteome per class for fungi (max two per phylum, n = 8), and the best proteome per order for embryophytes (max three per class, n = 12). The dataset was also supplemented with transcriptome-predicted proteomes from two species of CRuMs (*Rigifila ramosa*: SRR5997435, *Diphylleia rotans*: SRR5997435). Each individual proteome was then clustered at 100% sequence identity using CD-HIT v4.8.1 and the resulting proteomes were combined and compiled into a searchable database using Diamond v2.0.9 (*85, 86*).

To identify Exo70 homologs, the database was searched using Diamond BLASTp with *Arabidopsis thaliana* Exo70A1 (UniProt accession: Q9LZD3) and Exo84A (UniProt accession: F4I4B6) as initial queries (query coverage ≥ 50%, E < 10-5, sensitive mode). The resulting hits were aligned using MAFFT v7.520, trimmed using trimAl v1.4.rev15 with a gap threshold of 50%, and a maximum-likelihood phylogeny was inferred with IQ-Tree v2.2.6 using the LG4M substitution model with topology support assessed using Shimodaira-Hasegawa approximate likelihood ratio tests (SH-aLRT, n = 1,000) (*87–90*). The phylogenies were inspected manually in FigTree v1.4 (http://tree.bio.ed.ac.uk/software/figtree/) and non-homologous sequences were excluded. This search was repeated twice iteratively, and the resulting homologs were aligned using the L-INS-i algorithm of MAFFT and used to generate a profile hidden Markov model (HMM). The proteomes were then searched a final time with the HMM using HMMER v3.4 (E < 10-5) and the hits were screened phylogenetically as described (*91*). Lastly, to check for proteins that were missed due to genomic mis-annotation, proteins identified from the predicted proteomes were used as queries for tBLASTn (E-value < 10−5) searches against eukaryotic genomes, downloaded from NCBI, and protein predictions were generated using Exonerate v2.2 (see https://github.com/nickatirwin/Phylogenomic-analysis) (*92, 93*). The final set of Exo70 and Exo84 homologs were finally aligned and used to generate an HMM.

To expand the identification of Exo70 and Exo84 homologs in the viridiplantae, a second dataset of predicted-proteomes from plants (n = 81), streptophyte algae (n = 13), and chlorophytes (n = 21) was assembled from a combination of sources, including UniProt and EukProt v3 (*94*). The resulting dataset was surveyed as described above, using two iterative rounds of HMMER searches. Lastly, to remove non-homologous sites, the HMMs were mapped to each protein using HMMER HMMScan (E < 10-5, domE < 10-5) and homologous regions were extracted. These regions were aligned, trimmed with a gap-threshold of 50%, and sequences with less than 50% (Exo70) or 25% (Exo84) trimmed alignment coverage were excluded. The final phylogenies were generated with IQ-Tree using the LG+C50+F+R10 (Exo70) and LG+C50+F+R7 (Exo84) substitution models, selected using ModelFinder (*95*). Phylogenies were visualized in IToL v6 (*96*) (https://itol.embl.de/shared/OK75j4e8edHZ). Raw data with datasets, sequences, alignments and phylogenies can be found in **Supplemental Dataset 4**.

### Bioinformatic analyses: Sequence analysis and ancestral state reconstruction

To characterize N-termini of Exo70 and Exo84, the secondary structure of each viridiplantae homolog was predicted using PSSPred v4 (*97*). Amino acid sequences were then recoded with secondary structure predictions and were subsequently aligned using MAFFT. The alignments were visualized using AliView v1.28 (*98*) and the N-termini were identified and extracted (untrimmed sites: 0 to 9870 for Exo70, and 0 to 2906 for Exo84). The N-terminal and C-terminal regions were then recoded back to amino acids, clustered at 95% identity using CD-Hit, and aligned using MAFFT L-INS-i. Plant Exo70 (I, II, and III) and Exo84 (A, B, C) paralogs were then separated, and the individual alignments were trimmed with a gap-threshold of 75% using trimAl. Shannon entropy was calculated for each site using the bio3D package in R v4.3.2 (*99*). Linear discriminant analysis (LDA) was conducted using the LDA implementation in the MASS package in R, based on amino acid composition of Exo70 N-terminal sequences. Model training was conducted using 30% of the data. To analyse electrostatics, charge at pH 7 was predicted using BioPython (Bio.SeqUtils) on N-terminal helices greater than 100 amino acids in length (*100*). Lastly, ancestral state reconstructions were conducted using the fastAnc function in PhyTools v2 based on the rooted Exo70 and Exo84 phylogenies and the length-normalized N-terminal charge of each sequence (*101*). Raw data for the N-terminal analysis can be found in **Supplemental Dataset 4**.

---

**Fig. S1.**
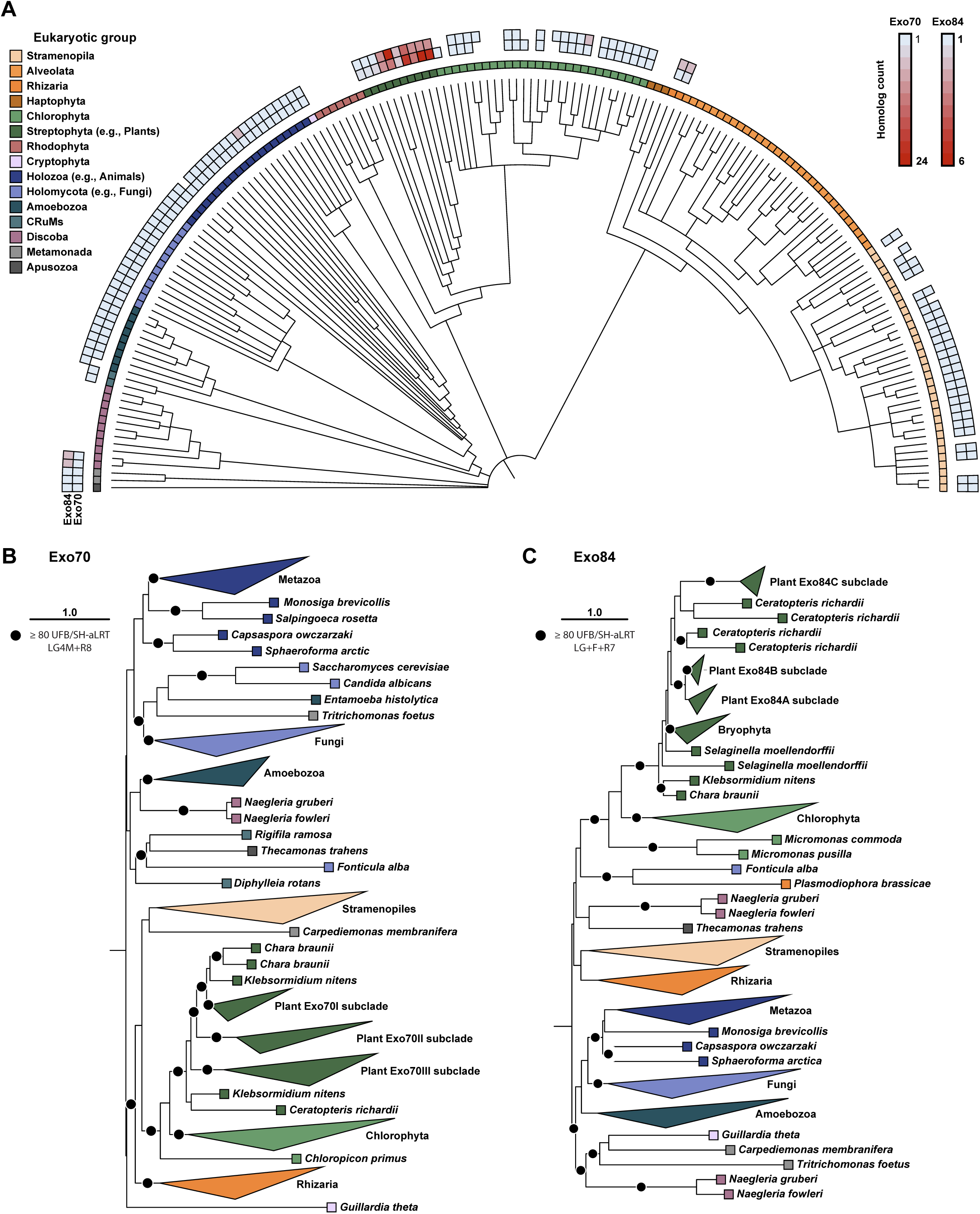
**(A)** A eukaryotic phylogeny depicting the frequency and abundance of Exo70 and Exo84 paralogs across representative species. Eukaryotic supergroups have been denoted with a colored ribbon. **(B, C)** Maximum likelihood phylogenies of eukaryotic Exo70 **(B)** and Exo84 **(C)** families. Statistical support was inferred using ultrafast bootstrap (UFB) and Shimodaira-Hasegawa approximate likelihood ratio tests (SH-aLRT). The scale bars represent the average number of substitutions per site. Full phylogenies are available in iTOL (*96*) (https://itol.embl.de/shared/OK75j4e8edHZ).

**Fig. S2.**
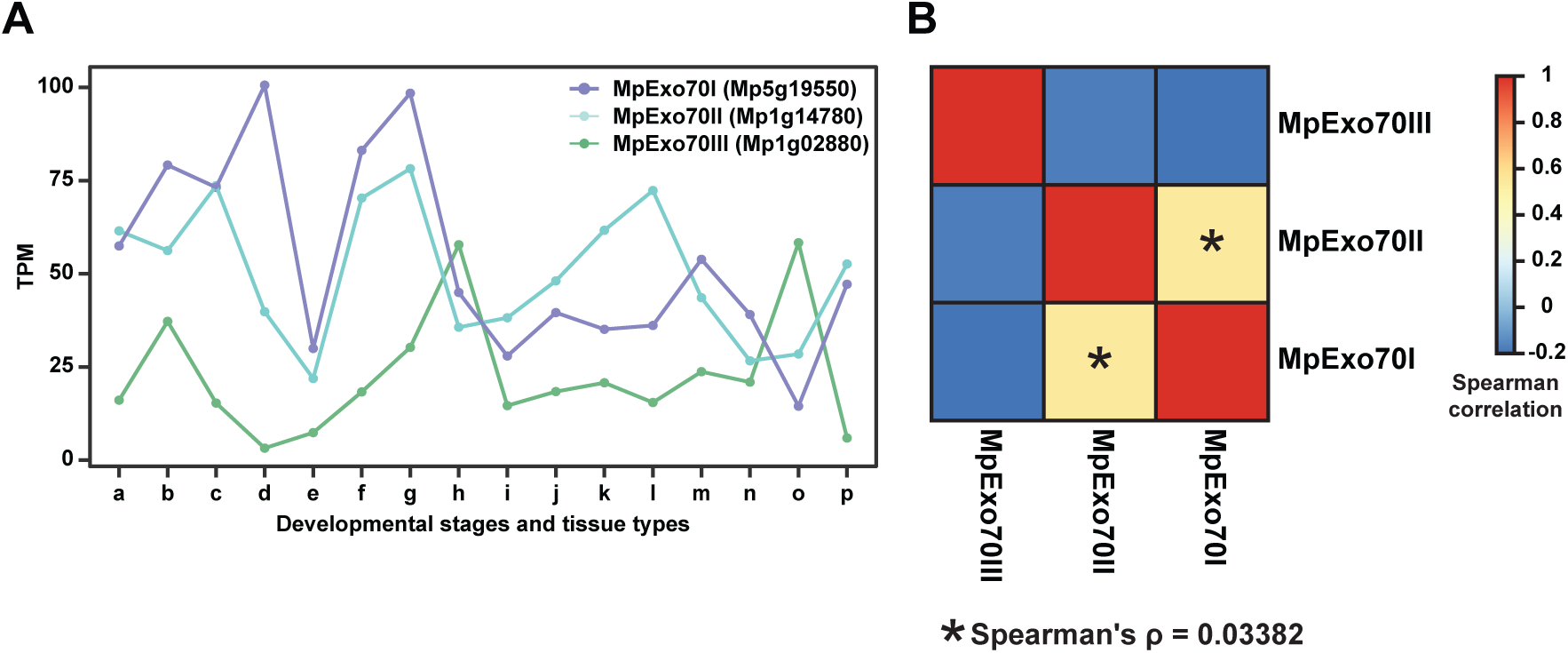
(A) Expression levels of Exo70 paralogs in Marchantia polymorpha across developmental stages and tissue types. Data was obtained from MarpolBase (Accessed July 2024). a: Tak-1 thallus 9-day male Higo et al., b: Tak-1 thallus 14-day male Karaaslan et al., c: Tak-1 antheridiophore mature male Higo et al., d: Tak-1 antheridium male Higo et al., e: Tak-1 sperm cell Julca et al., f: Tak-2 archegoniophore mature female Higo et al., g: Tak-2 archegonia female Hisanaga et al., h: Cam1 x Cam2 Spores 0 h Bowman et al., i: Cam1 x Cam2 Spores 24 h Bowman et al., j: Cam1 x Cam2 Spores 48 h Bowman et al., k: Cam1 x Cam2 Spores 72 h Bowman et al., l: Cam1 x Cam2 Spores 96 h Bowman et al., m: Tak-1 gemma cup 21 day male Ishizaki et al., n: Tak-1 midrib 21 day male Ishizaki et al., o: BC3 x Tak-1 gametophytic apical cell 9-day Frank et al., p: BC3 x Tak-1 young sporophyte 13-day Frank et al. **(B)** Spearman correlation between expression data for MpExo70I, MpExo70II and MpExo70III.

**Fig. S3.**
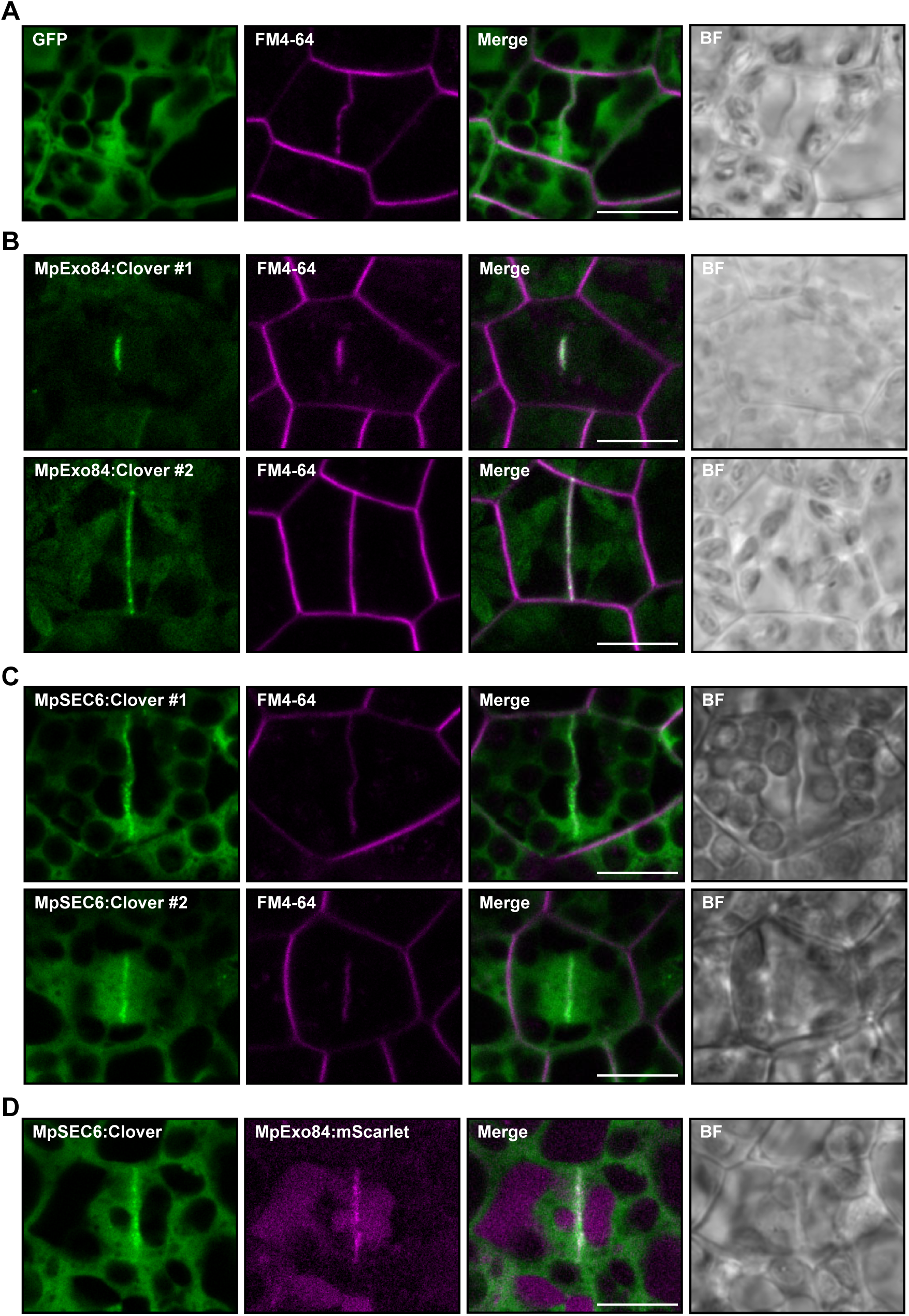
Marchantia exocyst localizes at the cell plate. Confocal micrographs of Marchantia cells stably expressing **(A)** free GFP, **(B)** MpExo84:Clover (two independent lines) and **(C)** MpSEC6:Clover (two independent lines) stained with FM4-64 (magenta). The presence of the cell plate is depicted by accumulation of FM4-64 stain. **(D)** Confocal micrographs of Marchantia cells stably co-expressing MpSEC6: Clover (green) or MpExo84:mScarlet (magenta). Scale bar is 10 µm. BF indicates Bright Field.

**Fig. S4.**
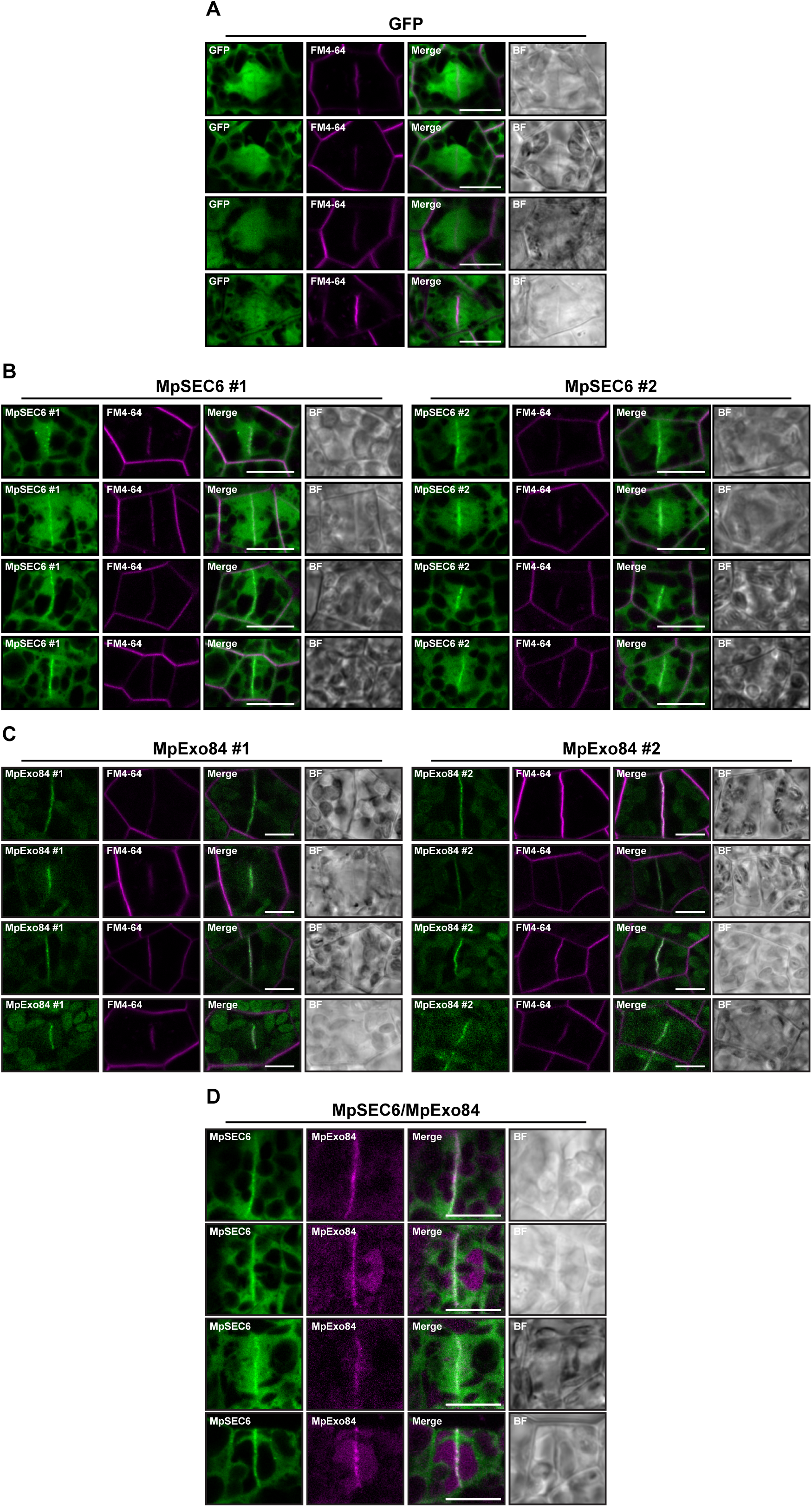
Image gallery for MpExo84 and MpSEC6 localization in Marchantia. Micrograph collection of Marchantia cells stably expressing **(A)** free GFP, **(B)** MpExo84:Clover (two independent lines) and **(C)** MpSEC6:Clover (two independent lines) stained with FM4-64 (magenta). The presence of the cell plate is depicted by accumulation of FM4-64 stain. **(D)** Micrograph collection of Marchantia cells stably co-expressing MpSEC6: Clover (green) or MpExo84:mScarlet (magenta). Scale bar is 10 µm. BF indicates Bright Field.

**Fig. S5.**
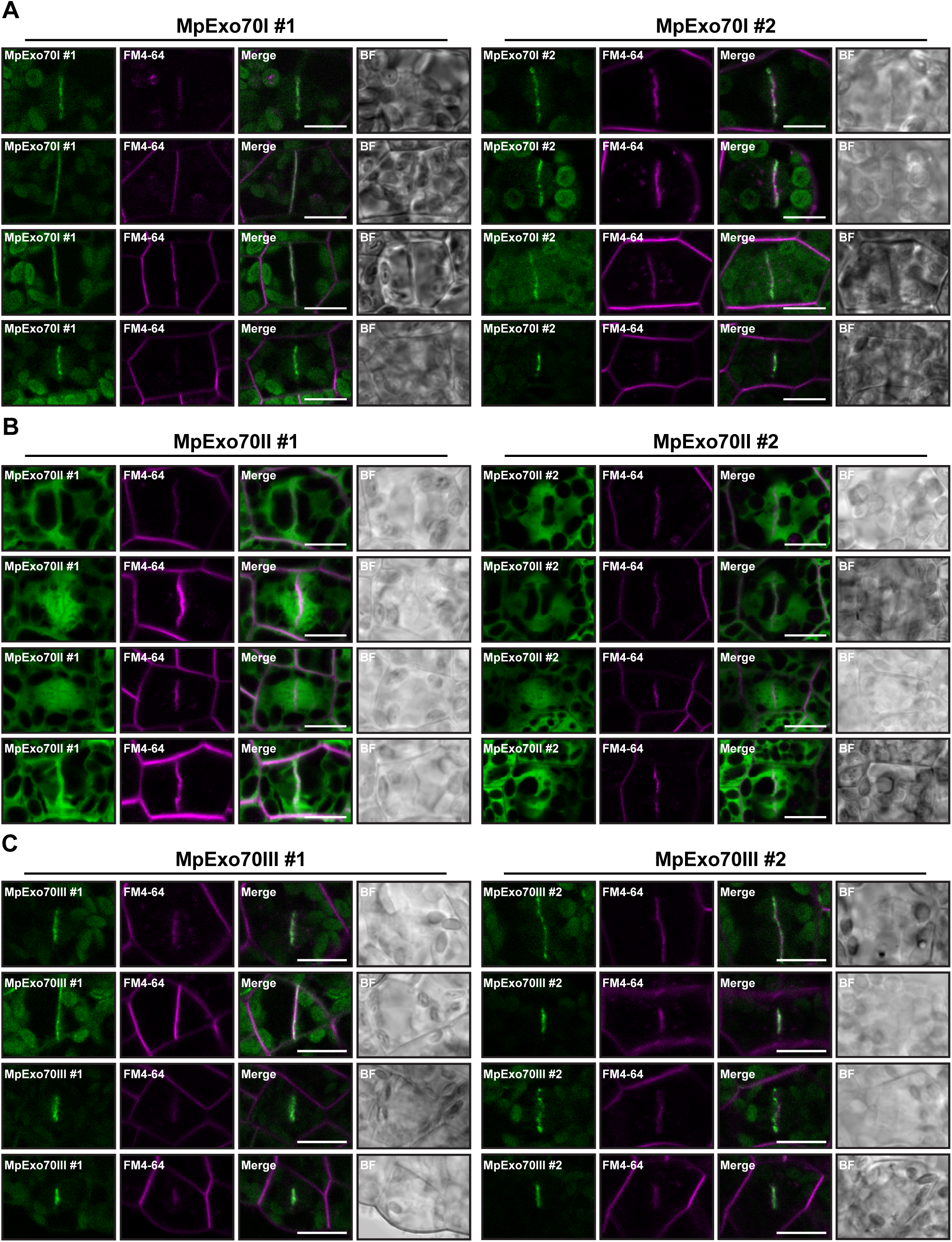
Image gallery for MpExo70 localization in Marchantia. Micrograph collection of Marchantia cells from two independent lines stably expressing **(A)** MpExo70I:Clover, **(B)** MpExo70II:Clover and **(C)** MpExo70III:Clover stained with FM4-64 (magenta). The presence of the cell plate is depicted by accumulation of FM4-64 stain. Scale bar is 10 µm. BF indicates Bright Field.

**Fig. S6.**
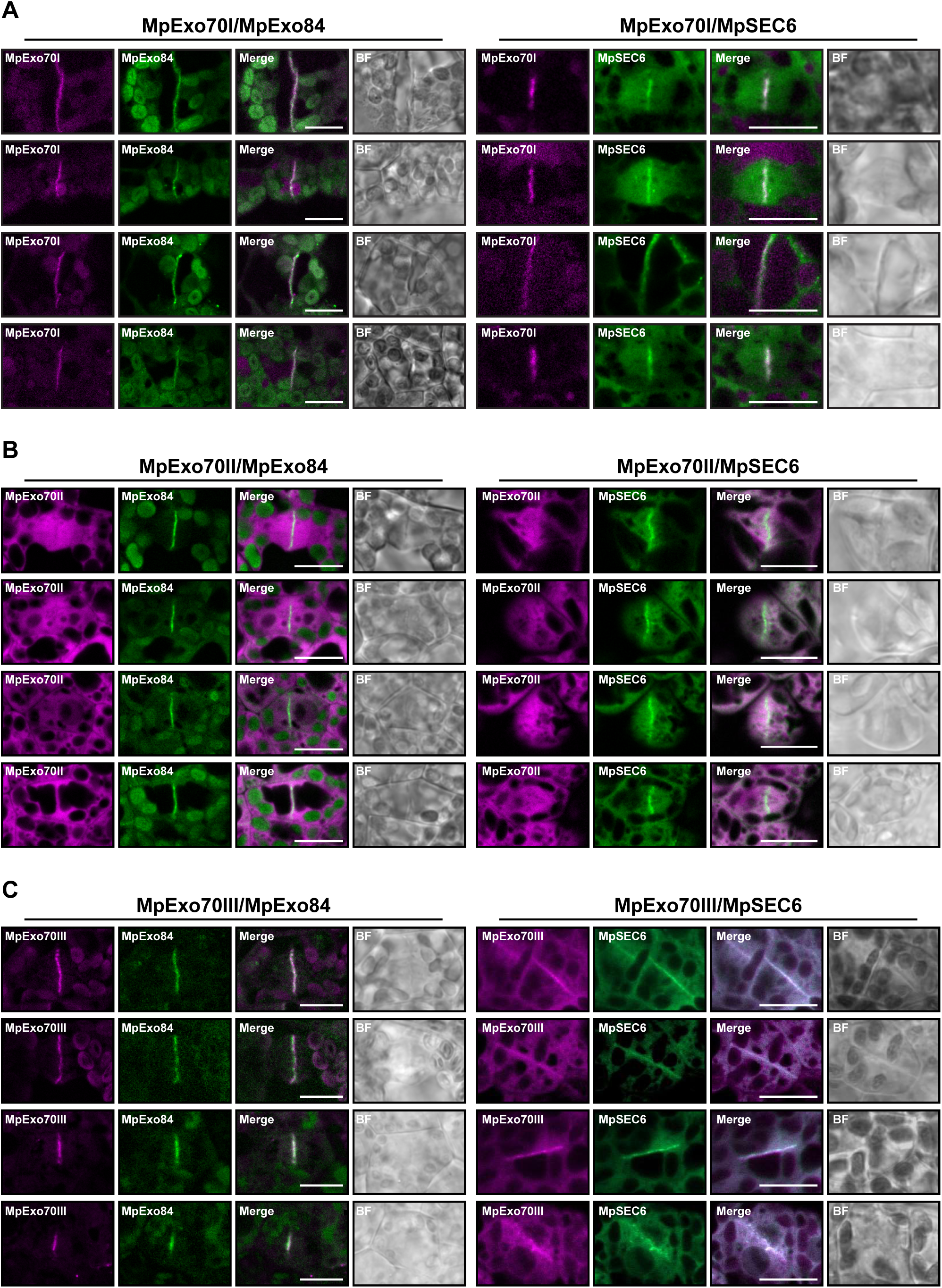
Image gallery for MpExo70 co-localization with MpExo84 and MpSEC6 in Marchantia. Micrograph collection of Marchantia cells stably co-expressing MpExo84:Clover or MpSEC6:Clover (green) with **(A)** MpExo70I:mScarlet, **(B)** MpExo70II:mScarlet and **(C)** MpExo70III:mScarlet (magenta). The presence of the cell plate is depicted by accumulation of either MpExo84:Clover or MpSEC6:Clover. Scale bar is 10 µm. BF indicates Bright Field.

**Fig. S7.**
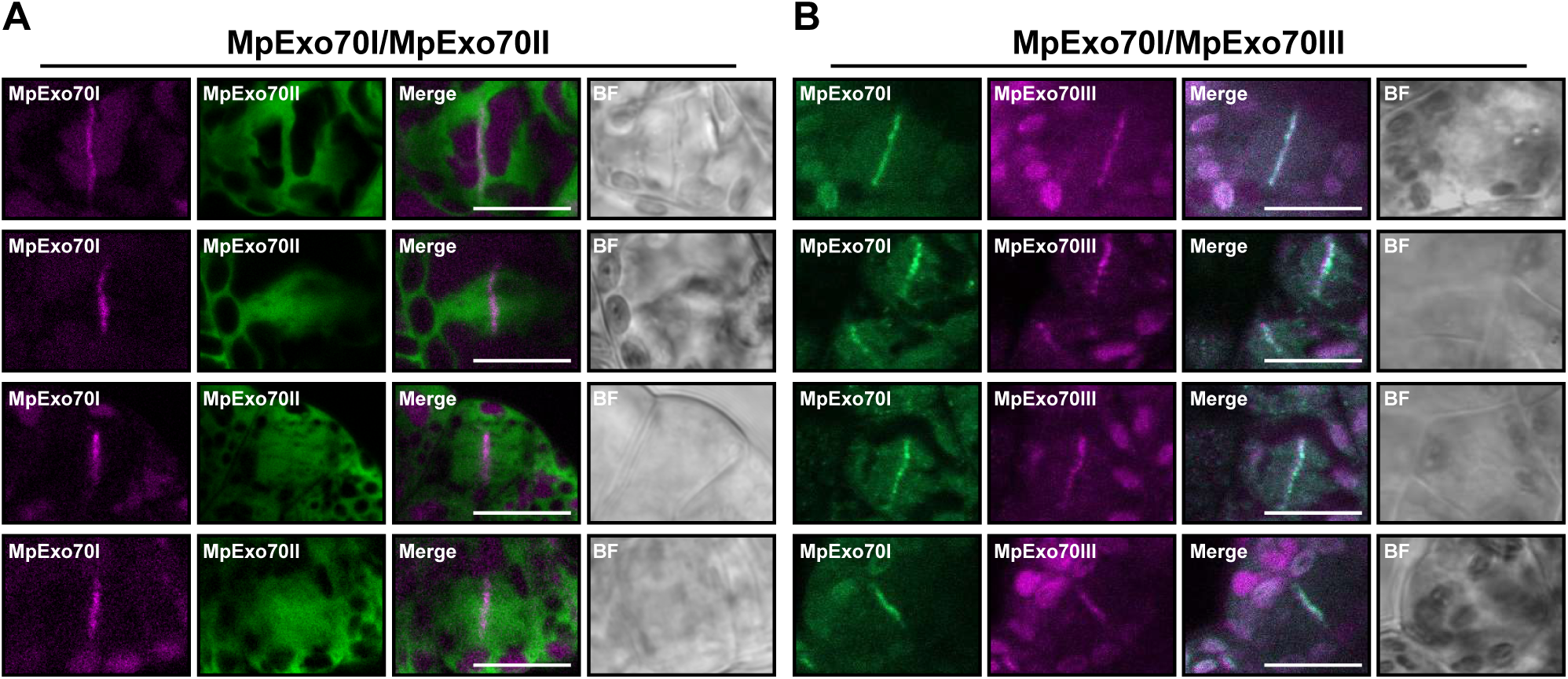
Co-localization of MpExo70 proteins in Marchantia cells. Micrograph collection of Marchantia cells stably co-expressing **(A)** MpExo70I:mScarlet (magenta) and MpExo70II:Clover (green) or **(B)** MpExo70I:Clover (green) and MpExo70III:mScarlet (magenta). Scale bar is 10 µm. BF indicates Bright Field.

**Fig. S8.**
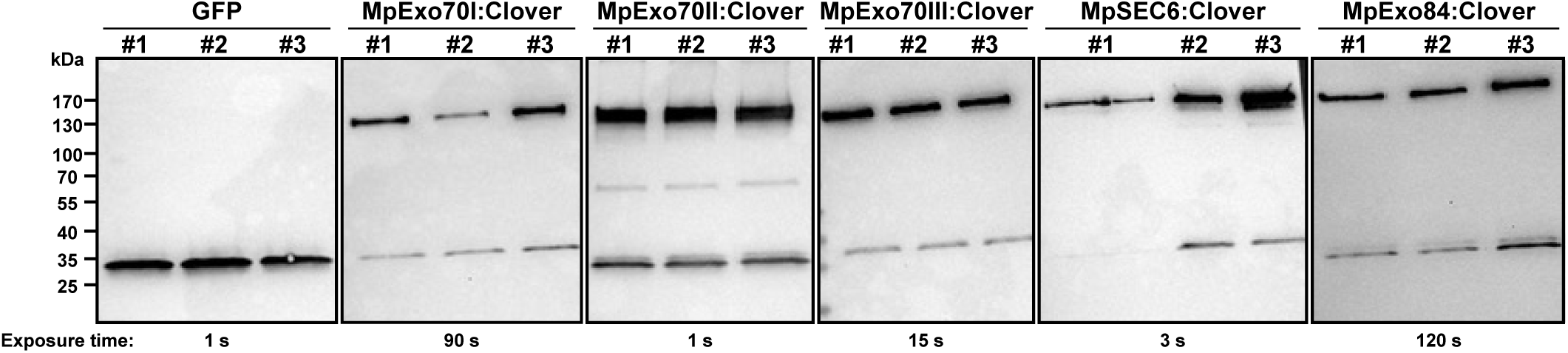
Protein accumulation in IP-MS samples analyzed by Western blot. Prior to mass-spectrometry analysis, immunoprecipitates obtained with anti-GFP magnetic beads were probed for the presence of free GFP, MpExo70I:Clover, MpExo70II:Clover, MpExo70III:Clover, MpSEC6:Clover or MpExo84:Clover using anti-GFP antibody. Exposure time is indicated below each panel as the accumulation of different proteins varied consistently.

**Fig. S9.**
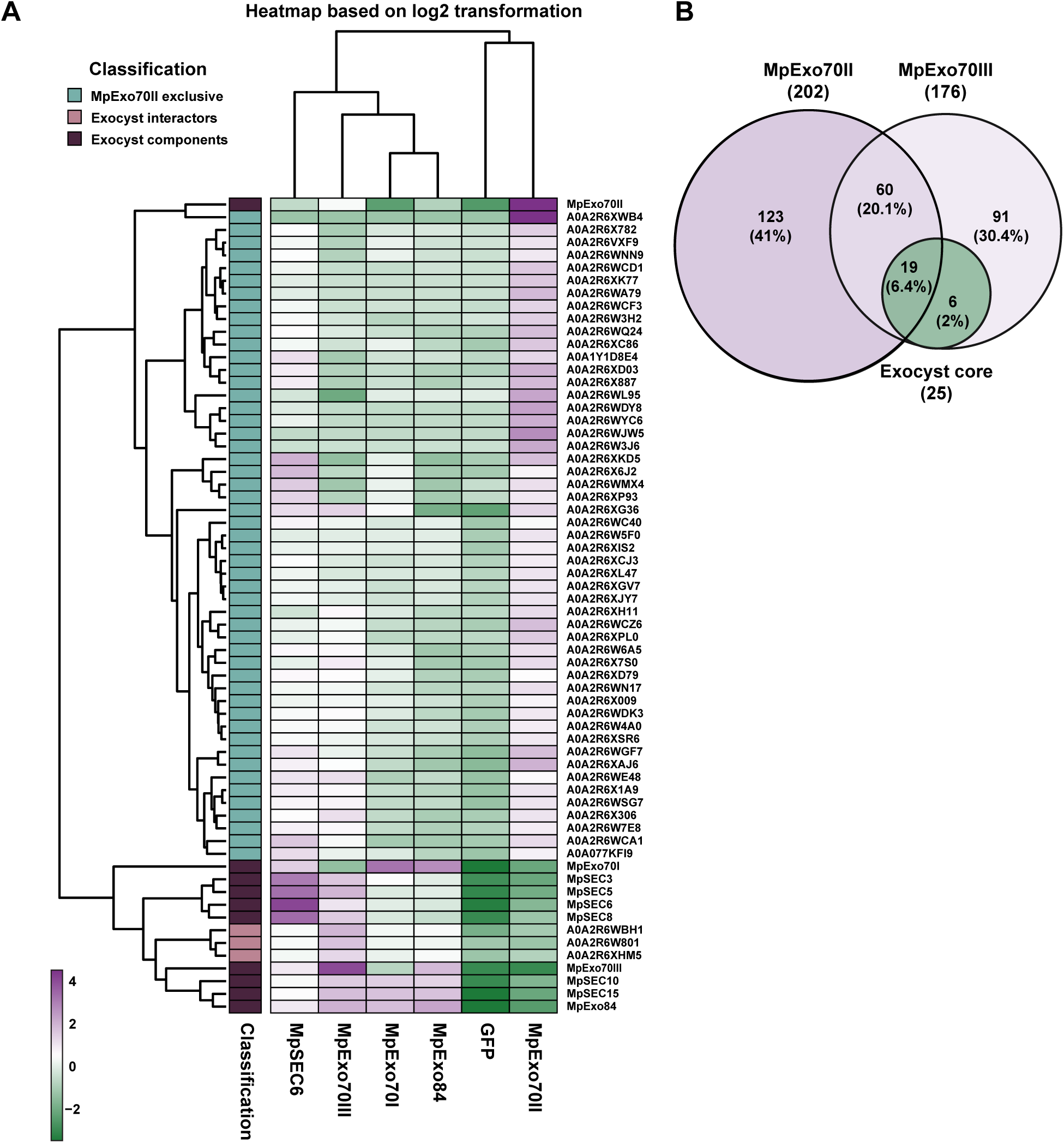
MpExo70 have different interactomes. **(A)** Protein abundance pattern represented by a heatmap (Log2(PSM+1) – meanPSM per protein, with values clipped between -3.5 and 4.5) for the exocyst components and the proteins identified as uniquely enriched in MpExo70II:Clover vs. GFP control dataset. **(B)** Venn diagram of three overlapping pairwise comparisons for AP-MS conducted in Marchantia: MpExo70I:Clover vs. GFP control (green circle), MpExo70II:Clover vs. GFP control (Purple circle) and MpExo70III:Clover vs. GFP control (Light purple circle). Total number of interactors for each pairwise comparison is indicated in brackets under the protein name. Number and percentage of shared interactors between pairwise comparisons are indicated in each overlapping area. Results represented are the mean from three independent replicates.

**Fig. S10.**
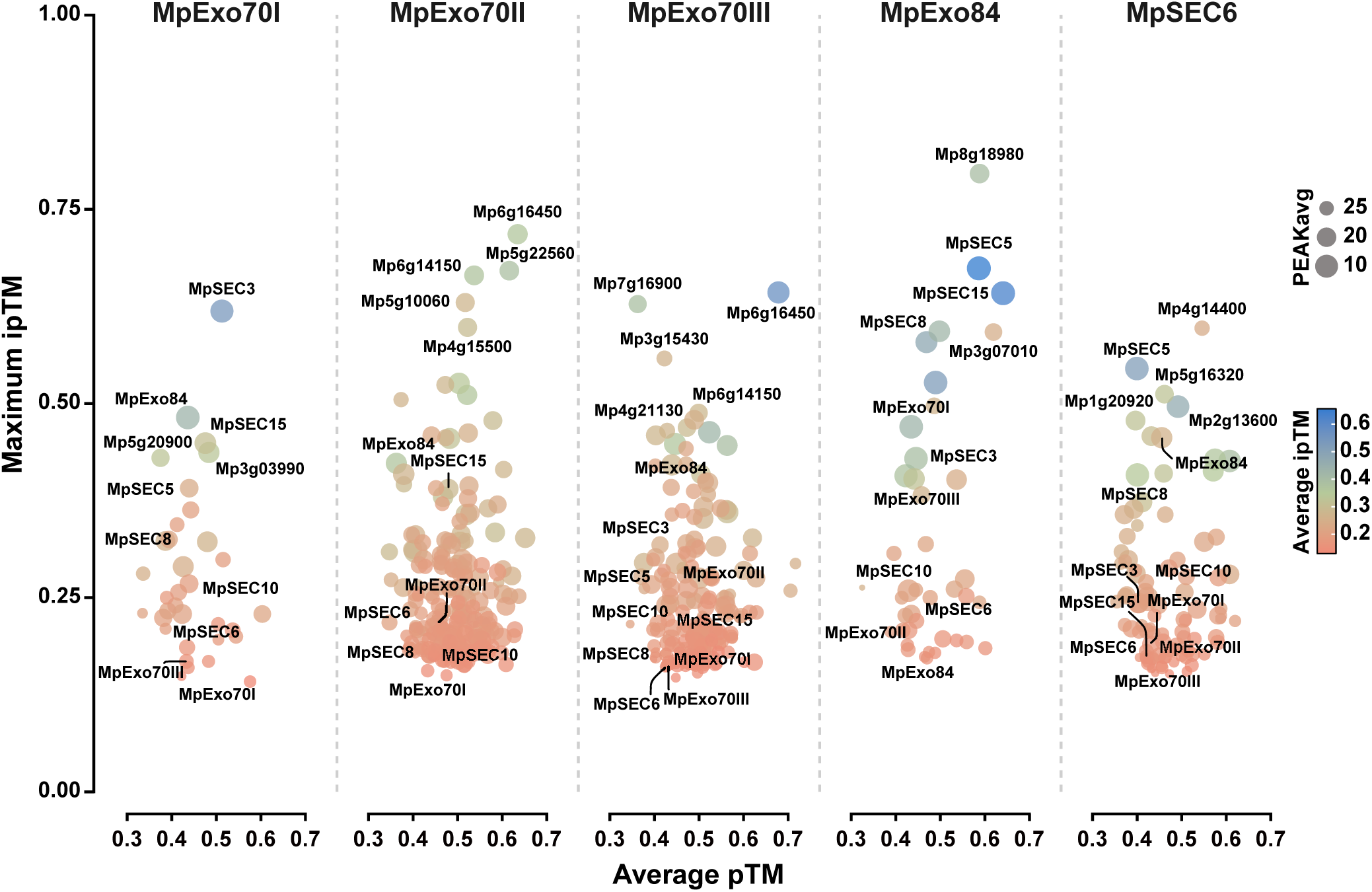
Analysis of MpExo70 interactors by AlphaFold2. Scatterplot of AlphaFold-Multimer (*78*) predicted ipTM versus pTM scores (ipTM: interface predicted Template Modeling score; pTM: predicted Template Modeling score) for MpExo70I, MpExo70II, MpExo70III, MpExo84 and MpSEC6 vs. their respective interactomes obtained by IP-MS. The maximum ipTM value from 5 independent predictions are use in the Y axis, while the average of pTM values from the 5 predictions is used in the X axis. The average ipTM is represented by the color of the dot. Dot size correlates to PEAK value average, where PEAK score represents average minimum predicted aligned error between protein chains excluding intra-molecular interactions. Top 5 candidates and exocyst components are indicated.

**Fig. S11.**
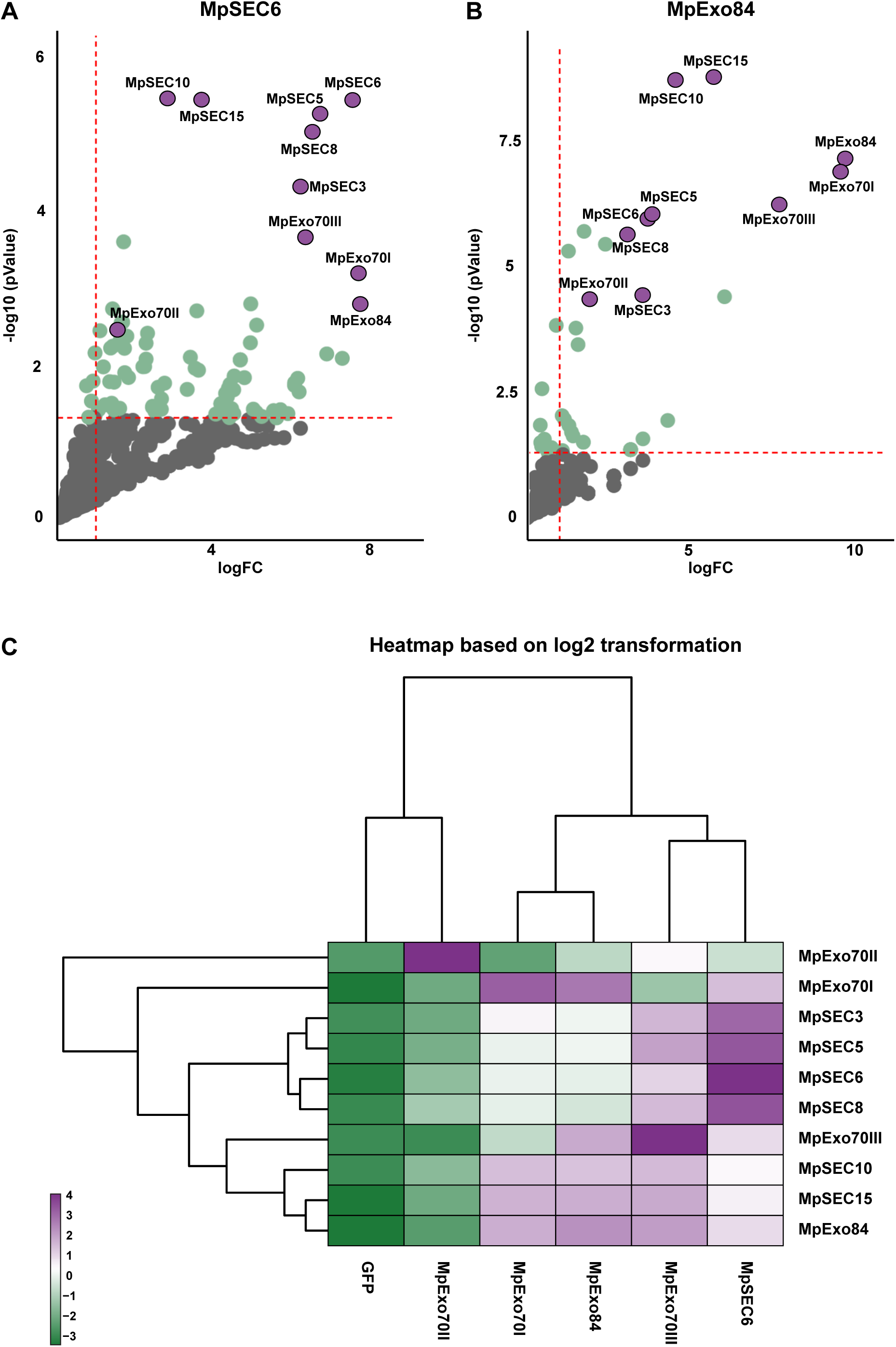
Abundance of exocyst components in the IP-MS datasets. Enrichment of proteins co-purified with **(A)** MpSEC6 and **(B)** MpExo84 compared to a control expressing free GFP and represented by a volcano plot. The horizontal dashed line indicates the threshold above which proteins are significantly enriched (p value < 0.05, quasi-likelihood negative binomial generalized log-linear model) and the vertical dashed line the threshold for which proteins log2 fold change is above 1. For each plot, members of Marchantia exocyst complex are depicted by a red dot with the corresponding name. **(C)** Protein abundance pattern represented by a heatmap (Log2(PSM+1) – meanPSM per protein, with values clipped between -3.5 and 4) for the exocyst components in each IP-MS dataset. Results represented are the mean from three independent replicates.

**Fig. S12.**
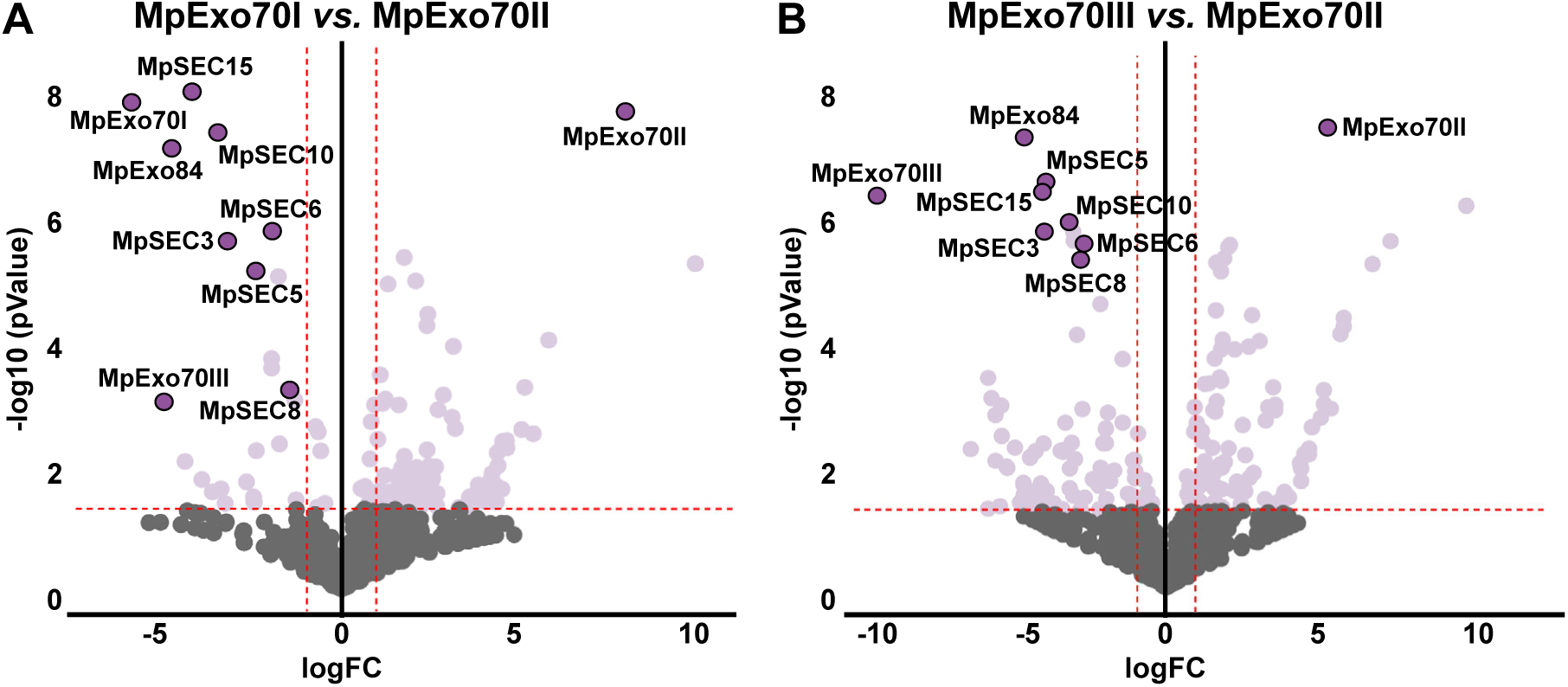
Exocyst components are less abundant in MpExo70II dataset. Enrichment of proteins co-purified with **(A)** MpExo70I and **(B)** MpExo70III compared to MpExo70II and represented by a volcano plot. The horizontal dashed line indicates the threshold above which proteins are significantly enriched (p value < 0.05, quasi-likelihood negative binomial generalized log-linear model) and the vertical dashed line the threshold for which proteins log2 fold change is above 1. For each plot, members of Marchantia exocyst complex are depicted by a red dot with the corresponding name.

**Fig. S13.**
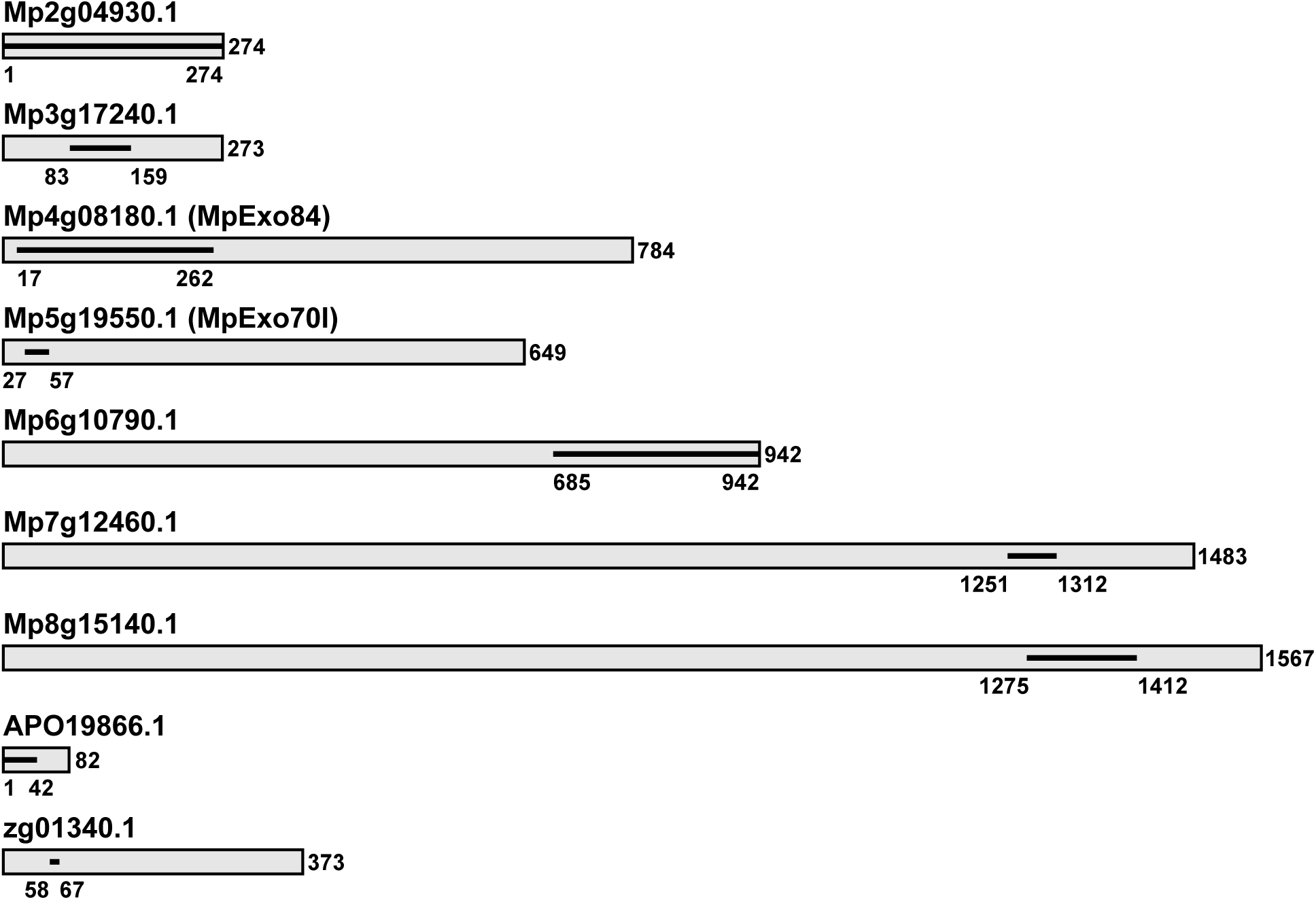
Schematic representation of MpExo70I interactors obtained by genome-wide yeast-two-hybrid. For each candidate, full-length protein is represented by a grey box. The line inside the box represents the domain for which an interacting clone was found.

**Fig. S14.**
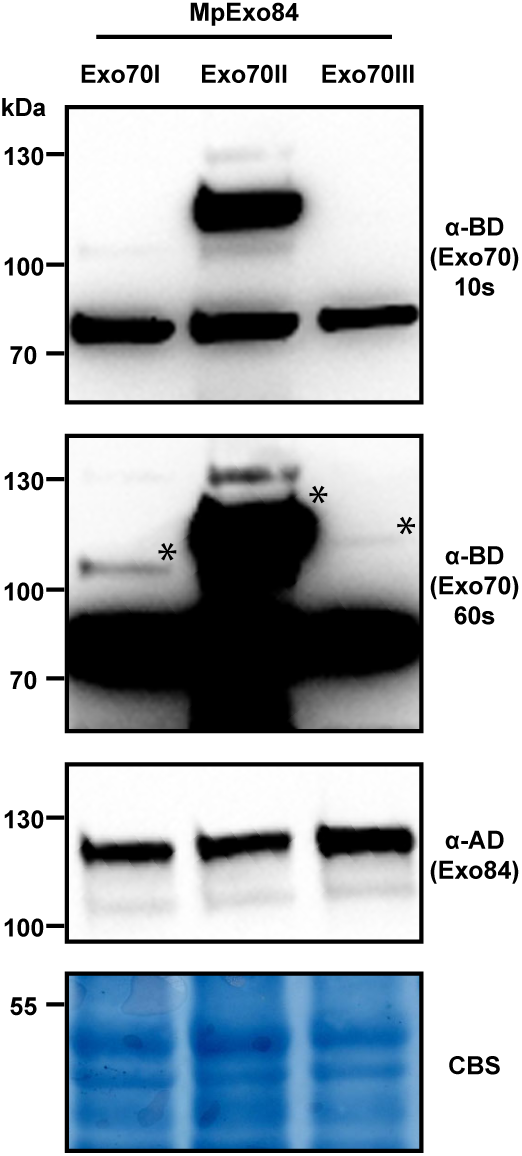
Protein accumulation in Yeast-Two-Hybrid assay analyzed by Western blot. Yeast lysate was probed for the presence of MpExo70I, MpExo70II and MpExo70III using anti-GAL4 binding domain (BD); and MpExo84 using anti-GAL4 DNA activation domain (AD) antibodies. Total protein extracts were stained with Coomassie Blue Stain (CBS). Accumulation of MpExo70II was consistently higher and two panels with indicated exposure times are included. Asterisks indicate the bands corresponding to the proteins of interest.

**Fig. S15.**
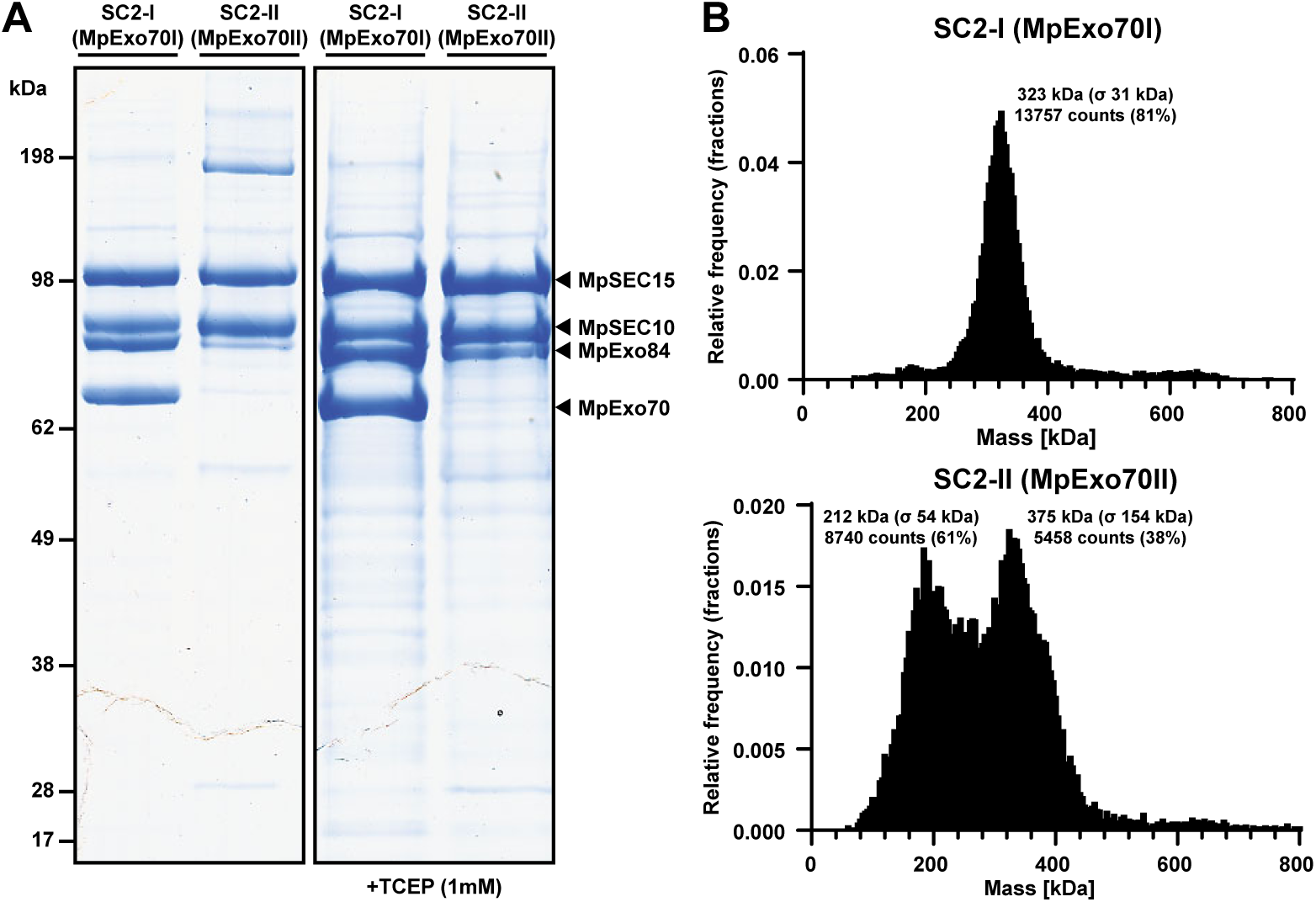
MpExo70II does not associate with exocyst proteins in purified subcomplexes. **(A)** SDS-PAGE analysis of purified Marchantia exocyst subcomplex 2-I (MpSEC15:MpSEC10:MpExo84:MpExo70I) and exocyst subcomplex 2-II (MpSEC15:MpSEC10:MpExo84:MpExo70II) produced in insect cells. The bands corresponding to MpSEC15:GFP:6XHis (MW: 118,7 kDa), MpSEC10 (92,2 kDa), MpExo84 (86,4 kDa) and MpExo70 (∼73 kDa) are indicated by arrows **(B)** Mass distribution of purified exocyst subcomplex 2-I (MW: 370 kDa) and exocyst subcomplex 2-II (MW: 374,3 kDa) estimated by mass photometry. For each peak, an estimated mass, standard deviation and number of counts is indicated.

**Fig. S16.**
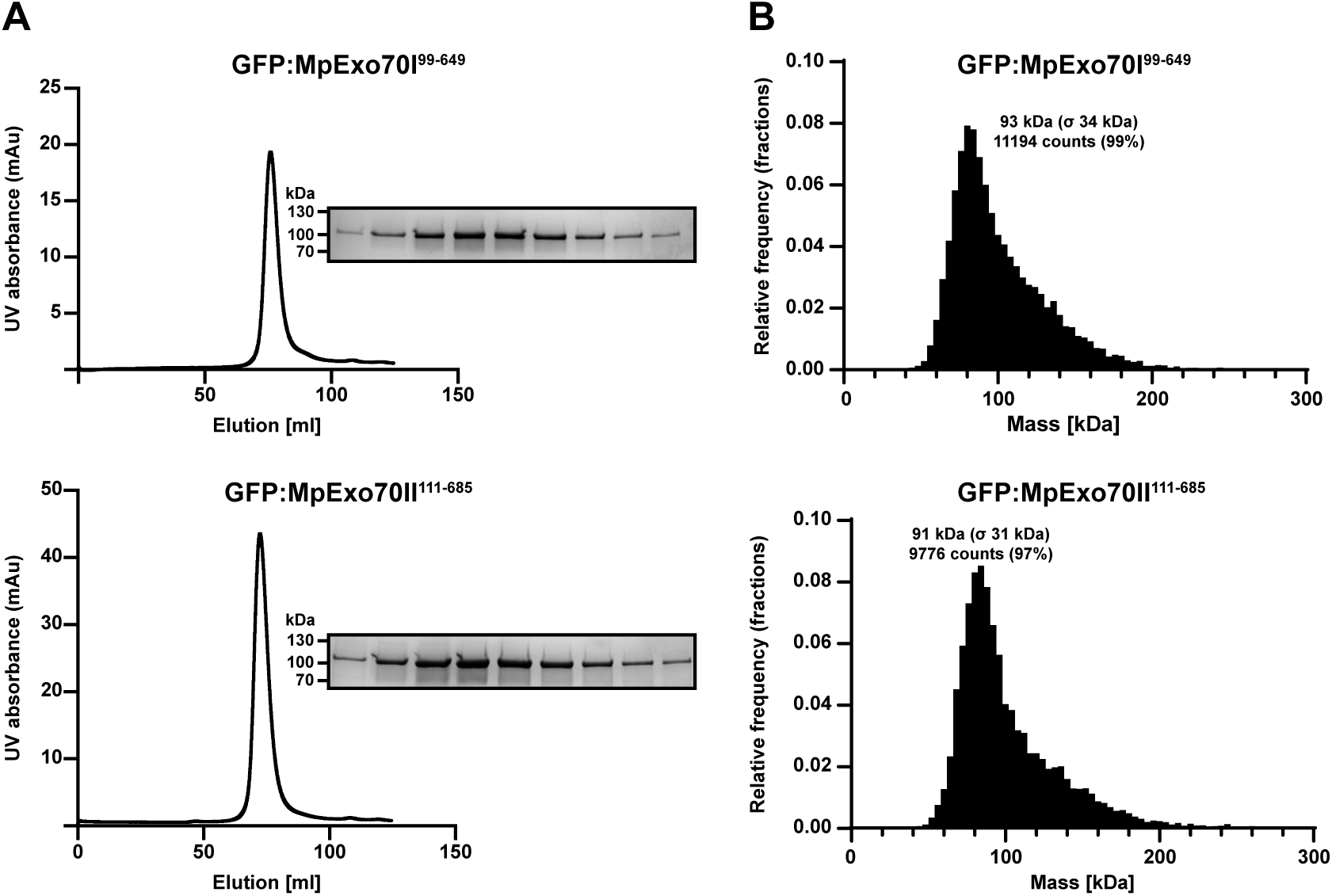
Purification of GFP:Exo70 proteins from insect cells. **(A)** Elution trace of GFP:MpExo70I99-649 and GFP:MpExo70II111-685 after gel filtration and SDS-PAGE analysis of relevant elution peak. **(B)** Mass distribution of purified GFP:MpExo70I99-649 (MW: 90 kDa) and GFP:MpExo70II111-685 (MW: 93,6 kDa) estimated by mass photometry. For each peak, an estimated mass, standard deviation and number of counts is indicated.

**Fig. S17.**
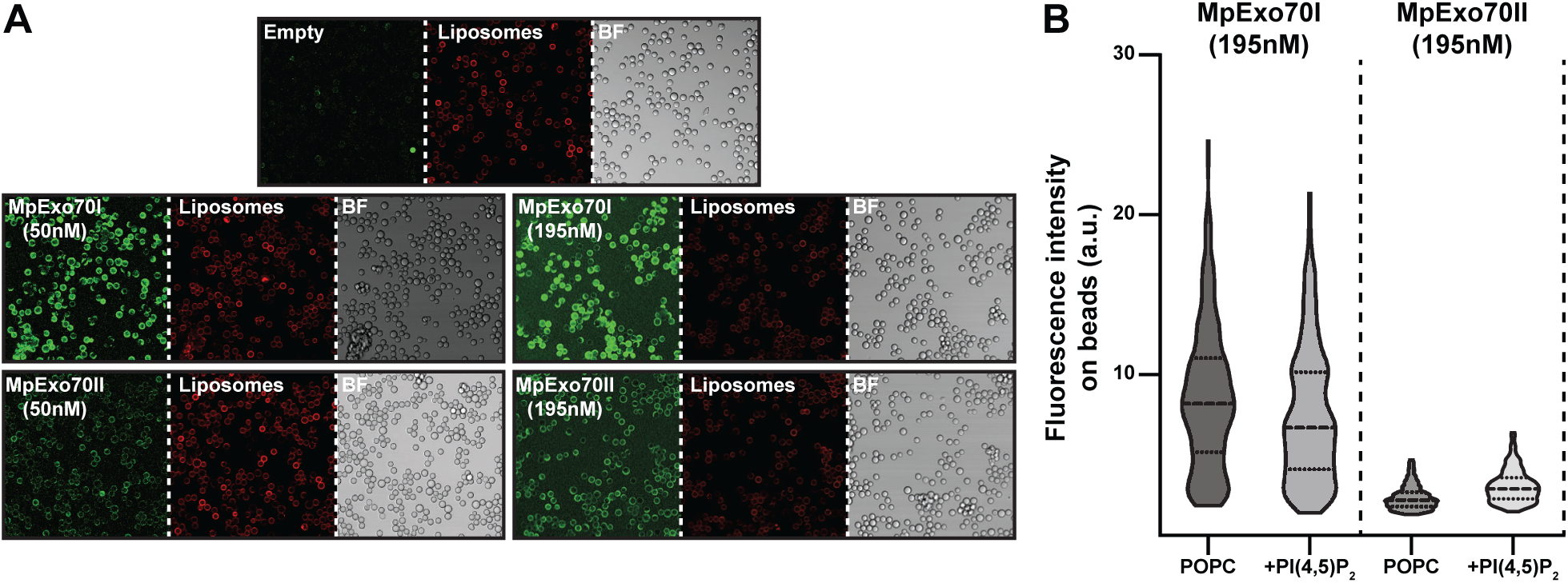
MpExo70I and MpExo70II binds liposome with different affinity. **(A)** Representative confocal microscopy images of purified GFP:MpExo70I99-649 and GFP:MpExo70II111-685 (green) added to liposomes (red) at 50 nM and 195 nM concentrations. BF indicates Bright Field. **(B)** Violin plots with median and quartiles of GFP:MpExo70I99-649 and GFP:MpExo70II111-685 fluorescence intensity in liposomes assayed at 195 nM concentration.

**Fig. S18.**
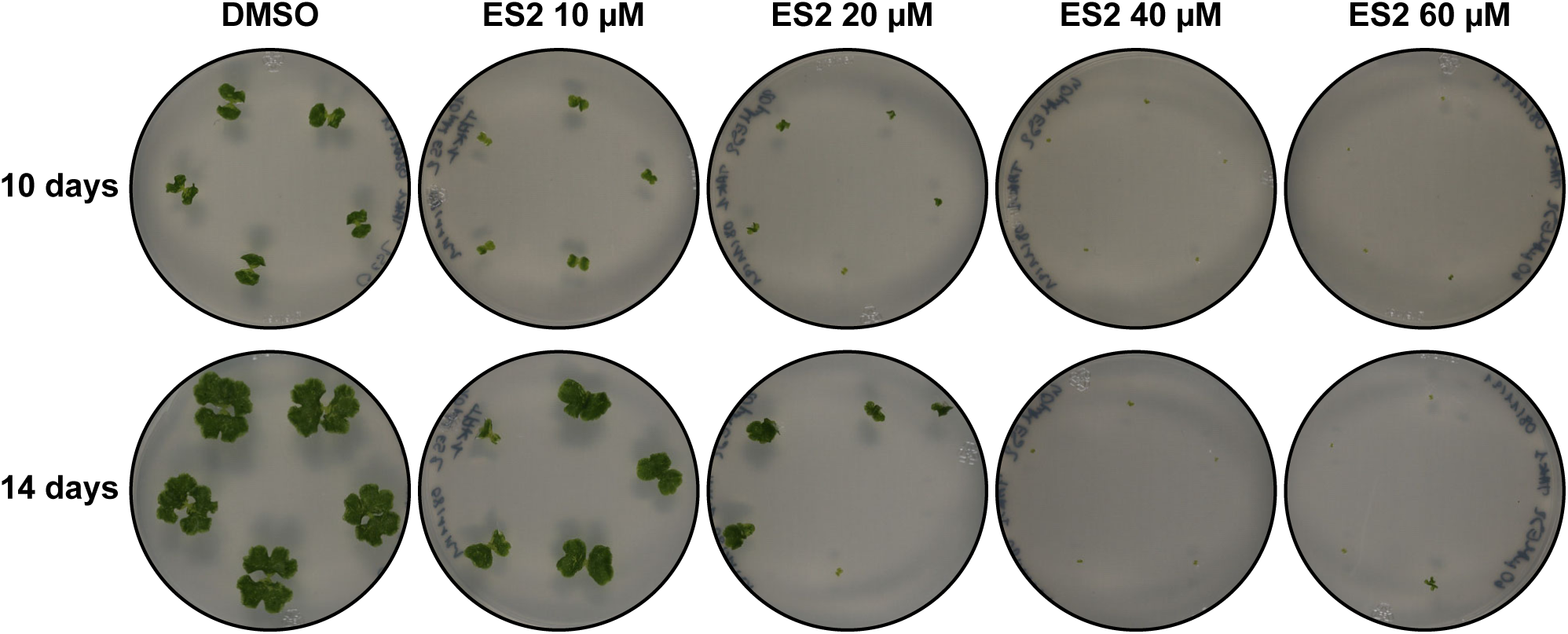
Endosidin 2 (ES2) inhibits Marchantia growth. Growth defect phenotypes of Marchantia Tak-1 grown for 10 or 14 days on media containing increasing concentrations of ES2 (0, 10, 20, 40 and 60 μM).

**Fig. S19.**
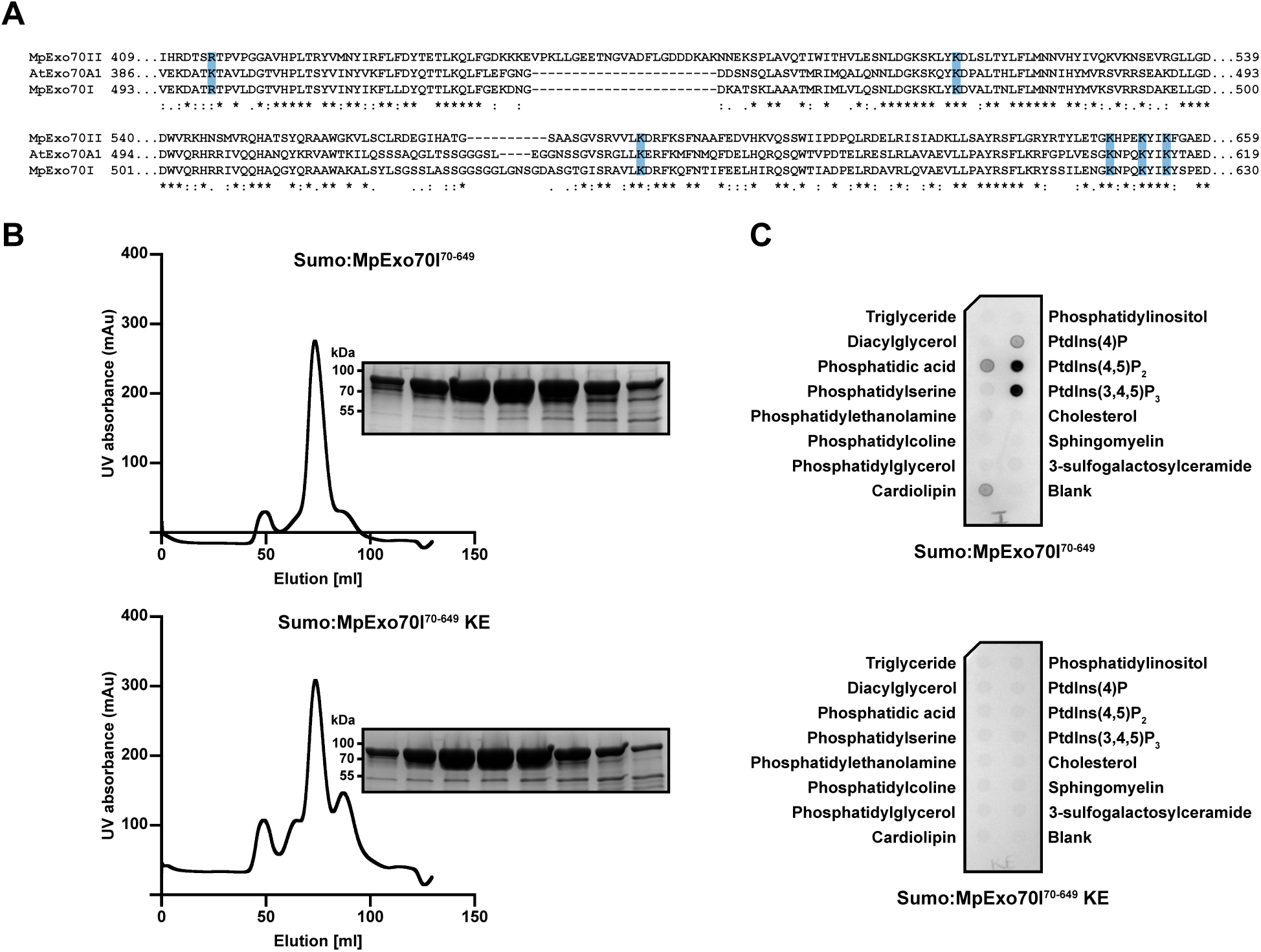
Lys to Glu (KE) mutations in MpExo70 abrogate binding to phospholipids. **(A)** Protein sequence alignment with Clustal Omega (*103*) for AtExo70A1, MpExo70I and MpExo70II at the C-terminal region containing the lipid binding domain. Conserved residues involved in lipid binding previously described for AtExo70A1 (*46*) are highlighted in blue. **(B)** Elution trace of SUMO:MpExo70I70-649 and SUMO:MpExo70I70-649 KE (MW: 78,5 kDa) produced in *E. coli* after gel filtration and SDS-PAGE analysis of relevant elution peak. **(C)** Protein–lipid overlay assays with purified SUMO:MpExo70I70-649 and SUMO:MpExo70I70-649 KE. Bound proteins were visualized using anti-SUMO antisera.

**Fig. S20.**
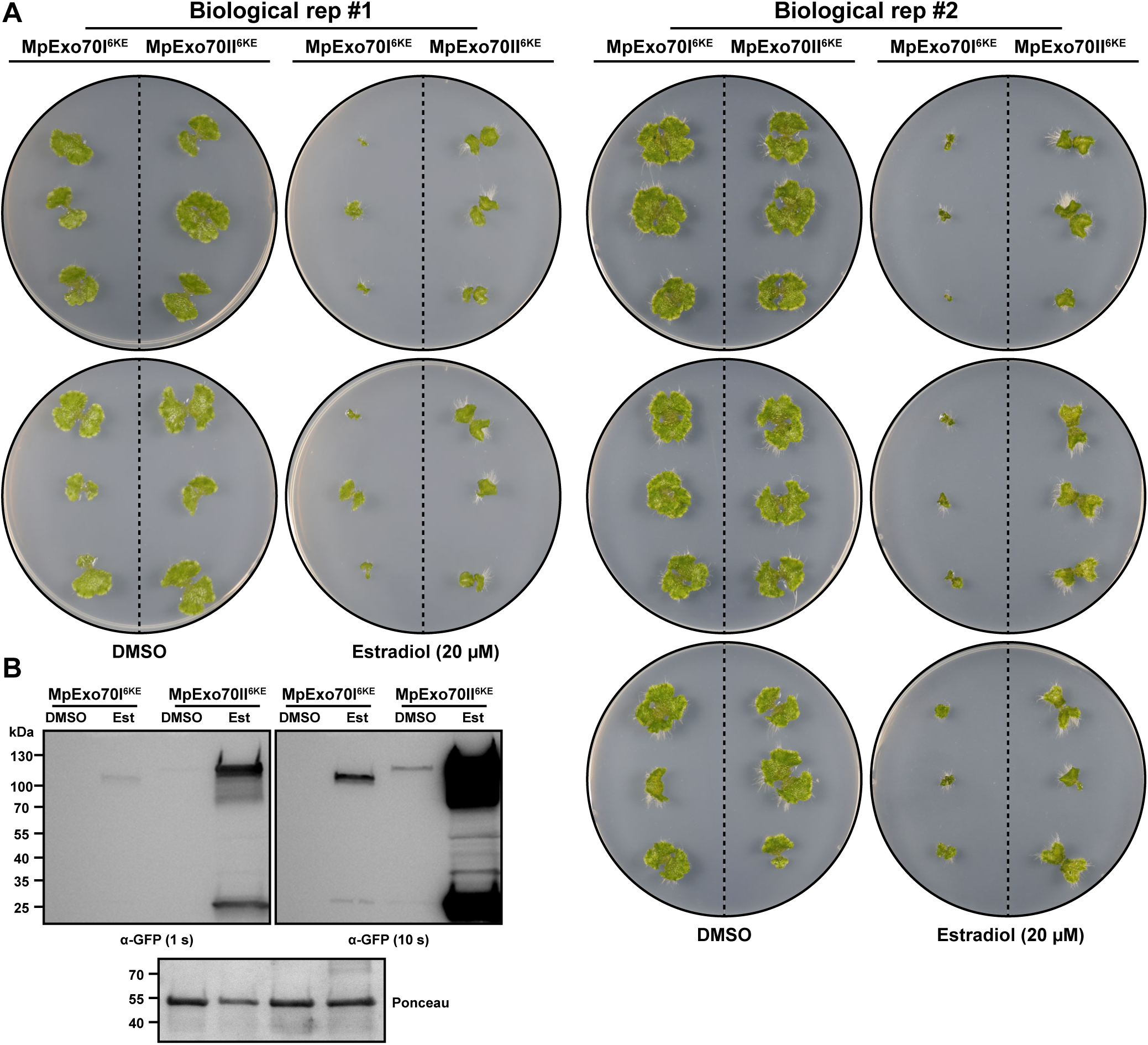
Experimental replicates of inducible expression of MpExo70 KE mutants in Marchantia. **(A)** Macroscopic phenotypes of Marchantia transgenic lines XVE:MpExo70I:Clover, XVE:MpExo70I 6KE:Clover, XVE:MpExo70II:Clover and XVE:MpExo70II 6KE:Clover grown for 14 days on media containing B-estradiol (20 µM) or DMSO. **(B)** Accumulation of MpExo70I 6KE:Clover and MpExo70II 6KE:Clover after estradiol-mediated induction visualized by western blot.

**Fig. S21.**
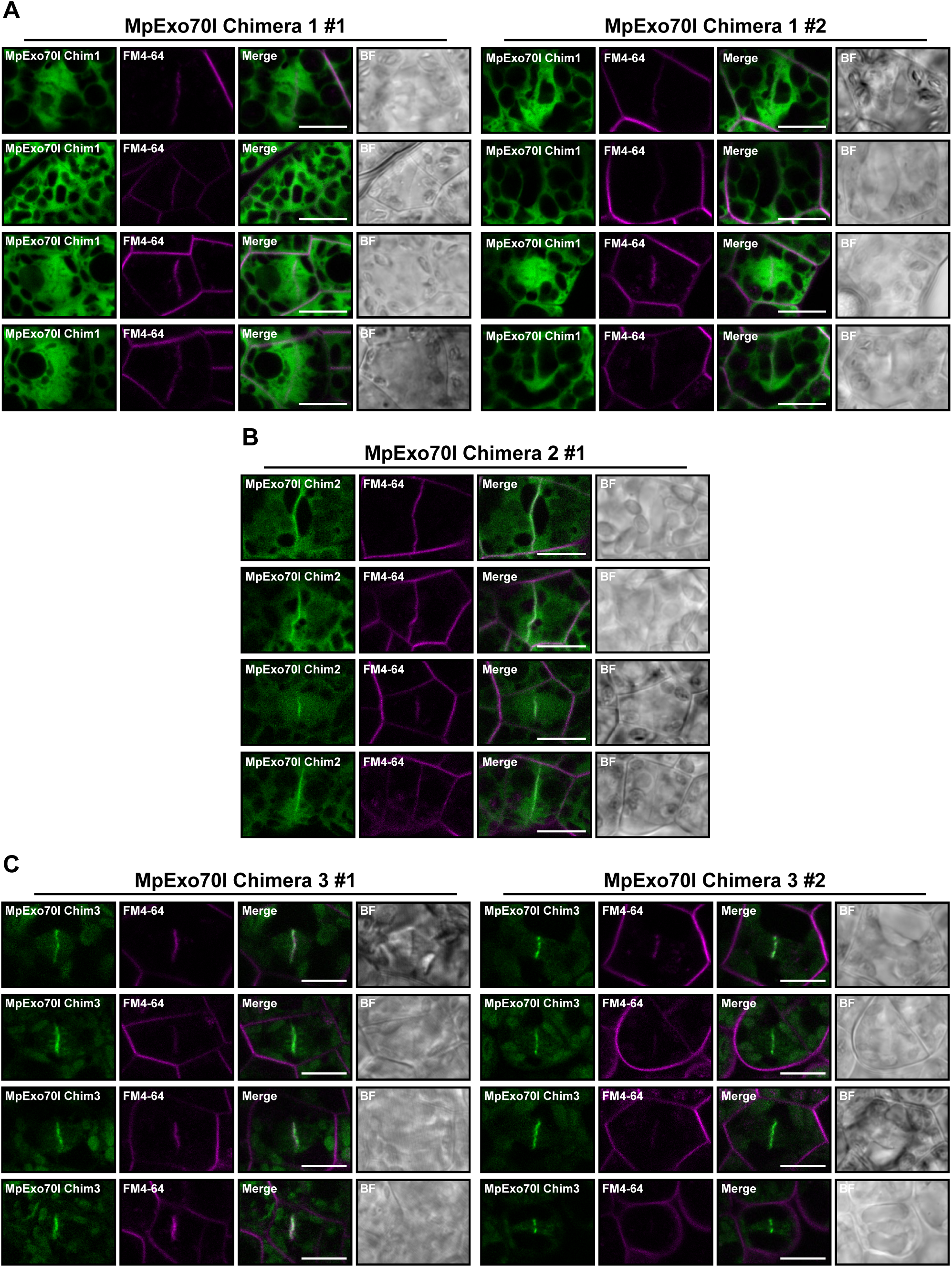
Image gallery for localization of MpExo70I chimeras in Marchantia. Micrograph collection of Marchantia cells stably expressing **(A)** MpExo70I chimera 1:Clover, **(B)** MpExo70I chimera 2:Clover and **(C)** MpExo70I chimera 3:Clover stained with FM4-64 (magenta). Two independent lines were imaged except for MpExo70I chimera 2. The presence of the cell plate is depicted by accumulation of FM4-64 stain. Scale bar is 10 µm. BF indicates Bright Field.

**Fig. S22.**
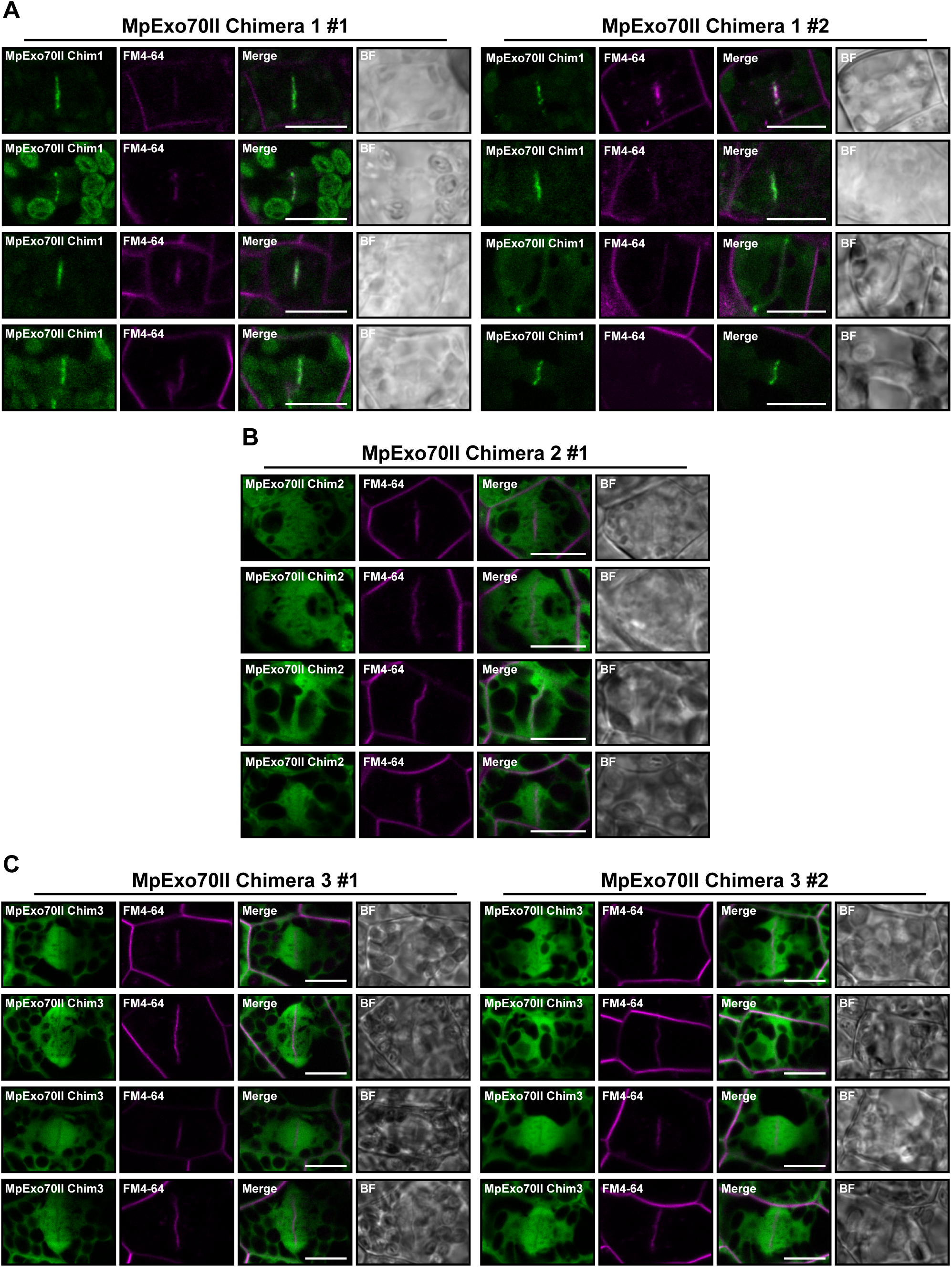
Image gallery for localization of MpExo70II chimeras in Marchantia. Micrograph collection of Marchantia cells stably expressing **(A)** MpExo70II chimera 1:Clover, **(B)** MpExo70II chimera 2:Clover and **(C)** MpExo70II chimera 3:Clover stained with FM4-64 (magenta). Two independent lines were imaged except for MpExo70II chimera 2. The presence of the cell plate is depicted by accumulation of FM4-64 stain. Scale bar is 10 µm. BF indicates Bright Field.

**Fig. S23.**
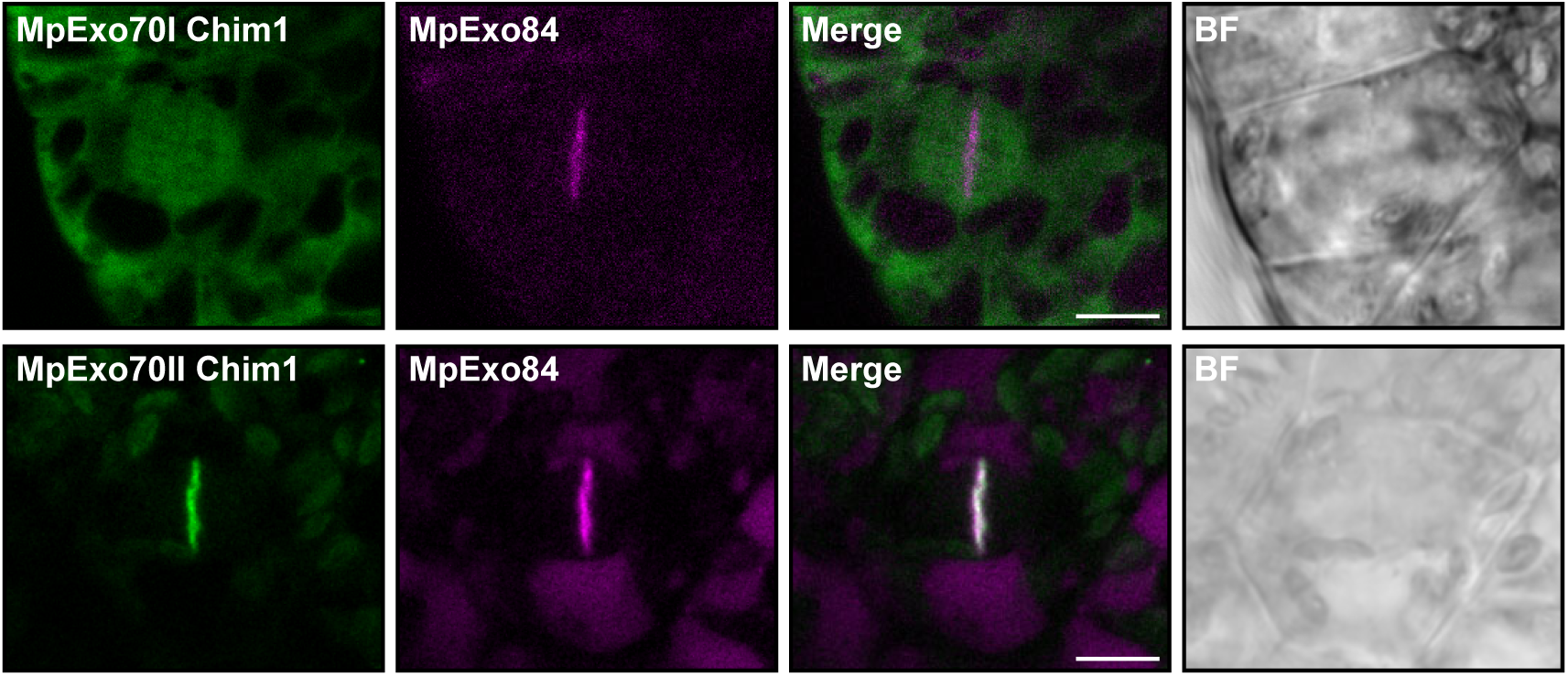
Co-localization of MpExo70 chimera 1 proteins with the exocyst complex. Confocal micrographs of Marchantia cells stably co-expressing **(A)** MpExo70I chimera 1:Clover or **(B)** MpExo70II chimera 1:Clover (green) and MpExo84:mScarlet (magenta). The presence of the cell plate is depicted by accumulation of MpExo84:mScarlet. Scale bar is 10 µm. BF indicates Bright Field.

**Fig. S24.**
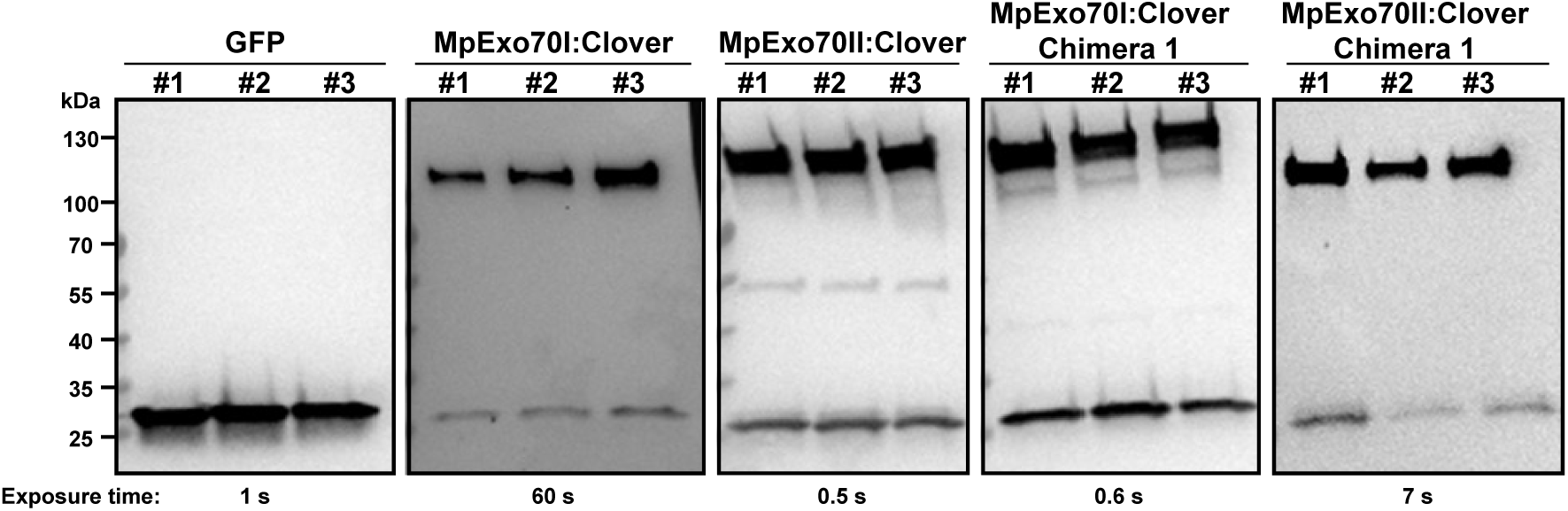
Protein accumulation in IP-MS samples analyzed by Western blot. Prior to mass-spectrometry analysis, immunoprecipitates obtained with anti-GFP magnetic beads were probed for the presence of free GFP, MpExo70I:Clover, MpExo70II:Clover, MpExo70I chimera 1:Clover or MpExo70II chimera 1:Clover using anti-GFP antibody. Exposure time is indicated below each panel as the accumulation of different proteins varied consistently.

**Fig. S25.**
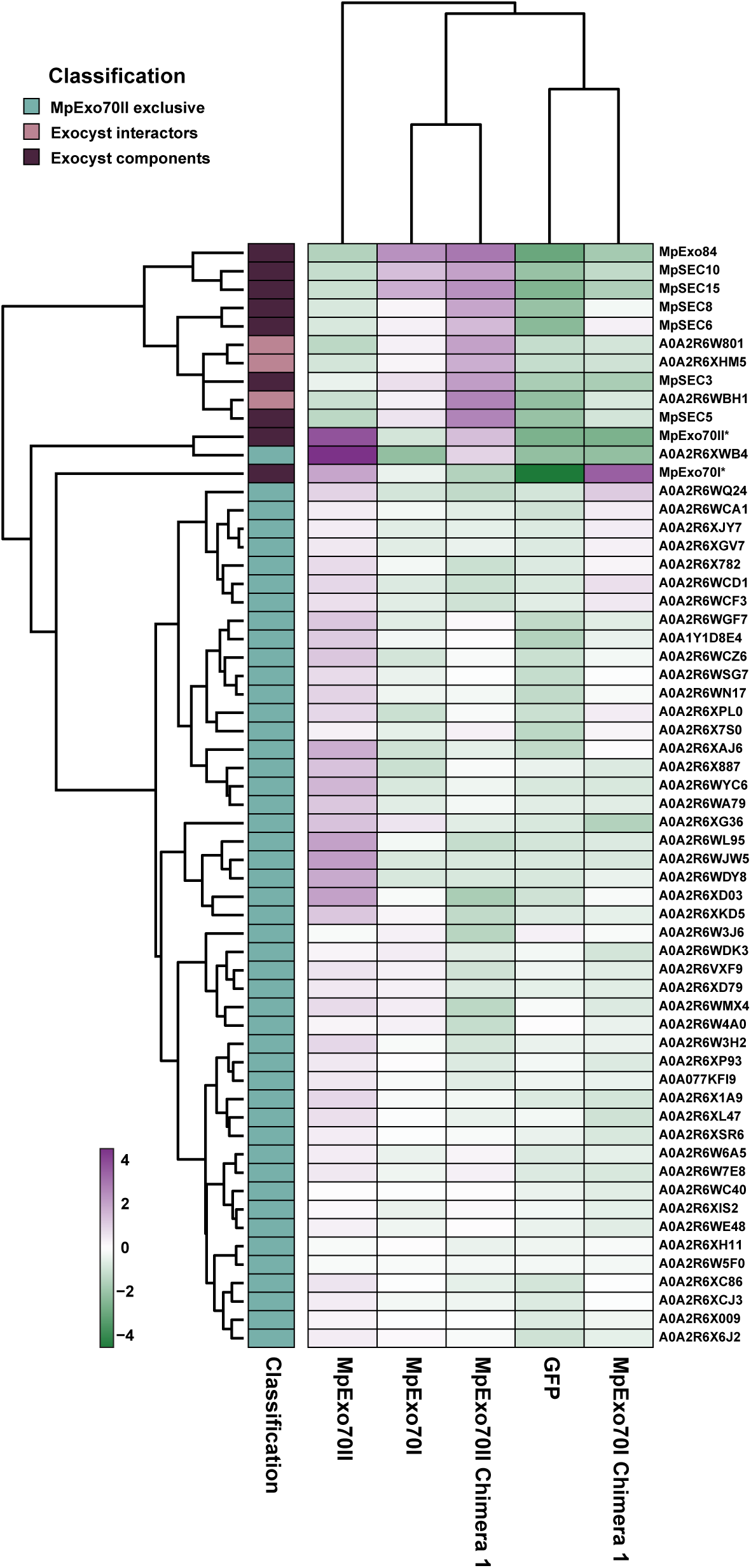
Replacement of N-terminal domain alter MpExo70s interactomes. Protein abundance pattern represented by a heatmap (Log2(PSM+1) – meanPSM per protein) for the exocyst components and the proteins identified as uniquely enriched in MpExo70II:Clover vs. GFP control dataset. Results represented are the mean from three independent replicates. Asterisk next to the name has been added as a cautionary mark indicating challenges in assigning the peptides to a chimera or a wild-type Exo70.

**Fig. S26.**
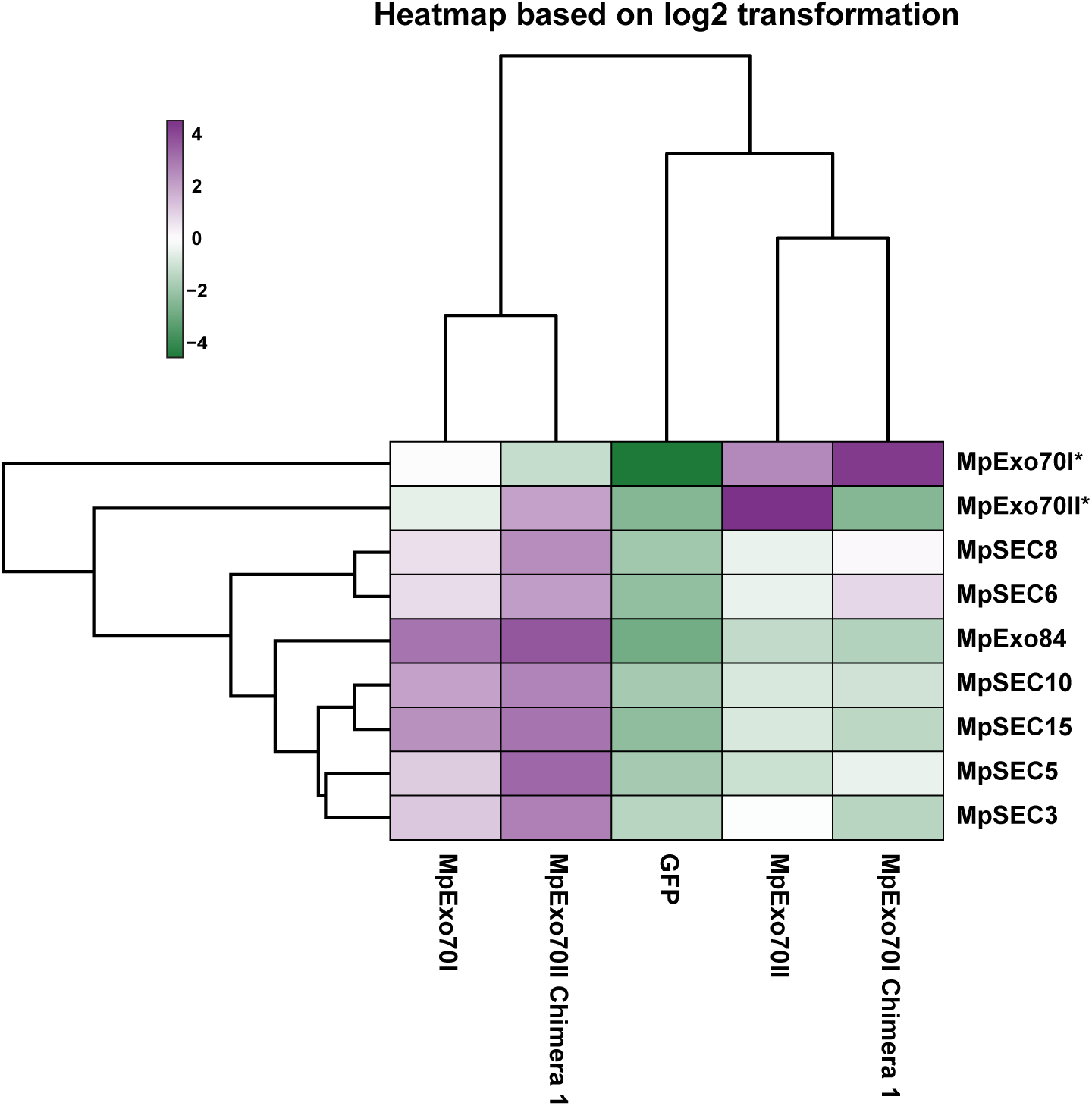
Abundance of exocyst components in the IP-MS datasets from MpExo70 chimeras. Protein abundance pattern represented by a heatmap (Log2(PSM+1) – meanPSM per protein) for the exocyst components in each IP-MS dataset. Results represented are the mean from three independent replicates. Asterisk next to the name has been added as a cautionary mark indicating challenges in assigning the peptides to a chimera or a wild-type Exo70.

**Fig. S27.**
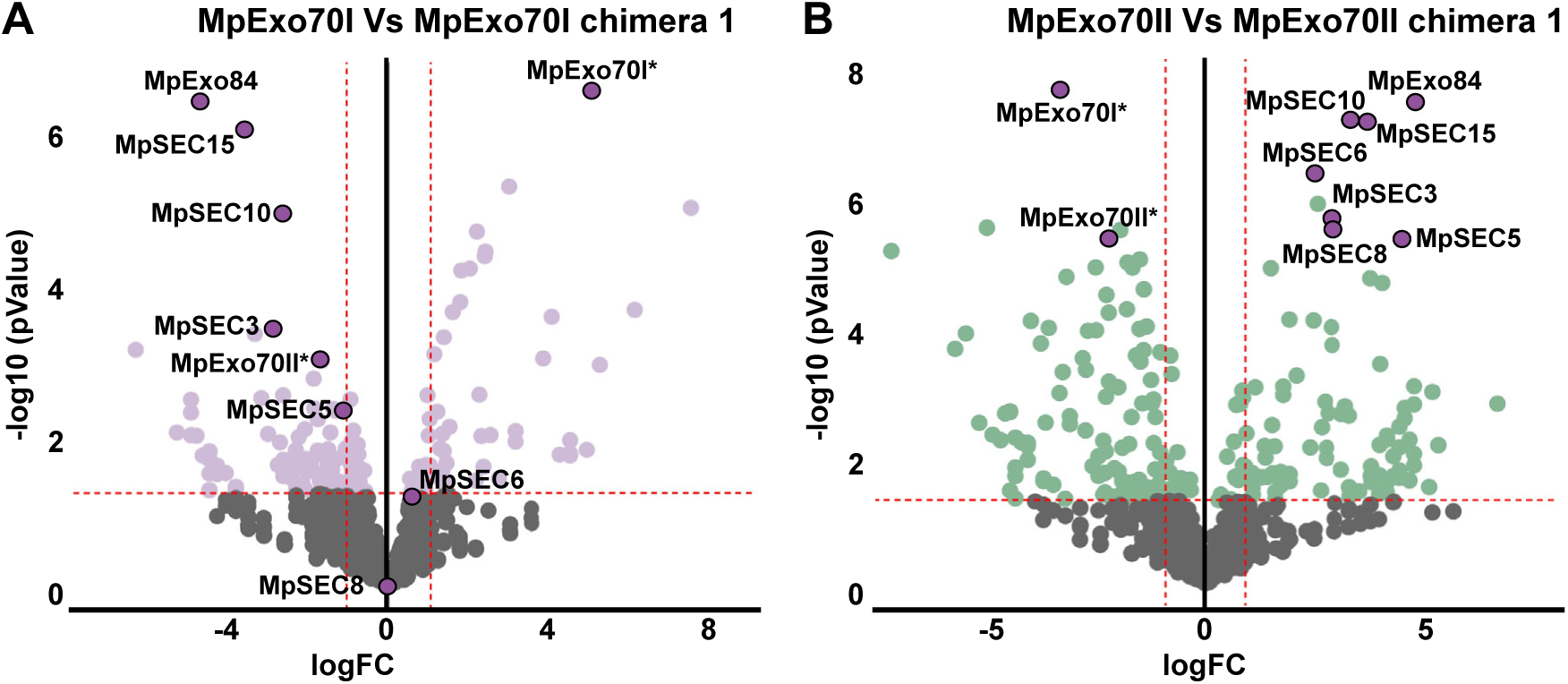
N-terminal domain swaps change MpExo70 association to exocyst complex. Enrichment of proteins co-purified with **(A)** MpExo70I vs. MpExo70I chimera 1 and **(B)** MpExo70II vs. MpExo70II chimera 1 represented by a volcano plot. The horizontal dashed line indicates the threshold above which proteins are significantly enriched (p value < 0.01, quasi-likelihood negative binomial generalized log-linear model) and the vertical dashed line the threshold for which proteins log2 fold change is above 1. For each plot, members of Marchantia exocyst complex are depicted by a red dot with the corresponding name. Asterisk next to the name has been added as a cautionary mark indicating challenges in assigning the peptides to a chimera or a wild-type Exo70.

**Fig. S28.**
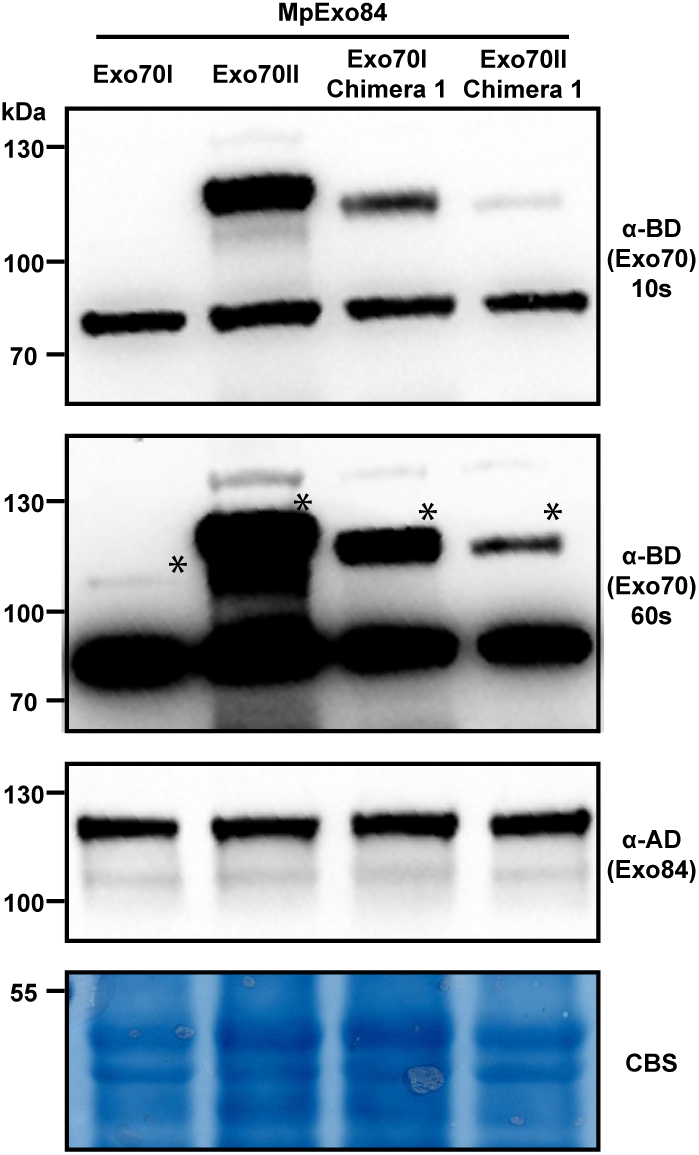
Protein accumulation in Yeast-Two-Hybrid assay analyzed by Western blot. Yeast lysate was probed for the presence of MpExo70I, MpExo70II, MpExo70I chimera 1 and MpExo70II chimera 1 using anti-GAL4 binding domain (BD); and MpExo84 using anti-GAL4 DNA activation domain (AD) antibodies. Total protein extracts were stained with Coomassie Blue Stain (CBS). Accumulation of MpExo70II and MpExo70I chimera 1 were consistently higher and two panels with indicated exposure times are included. Asterisks indicate the bands corresponding to the proteins of interest.

**Fig. S29.**
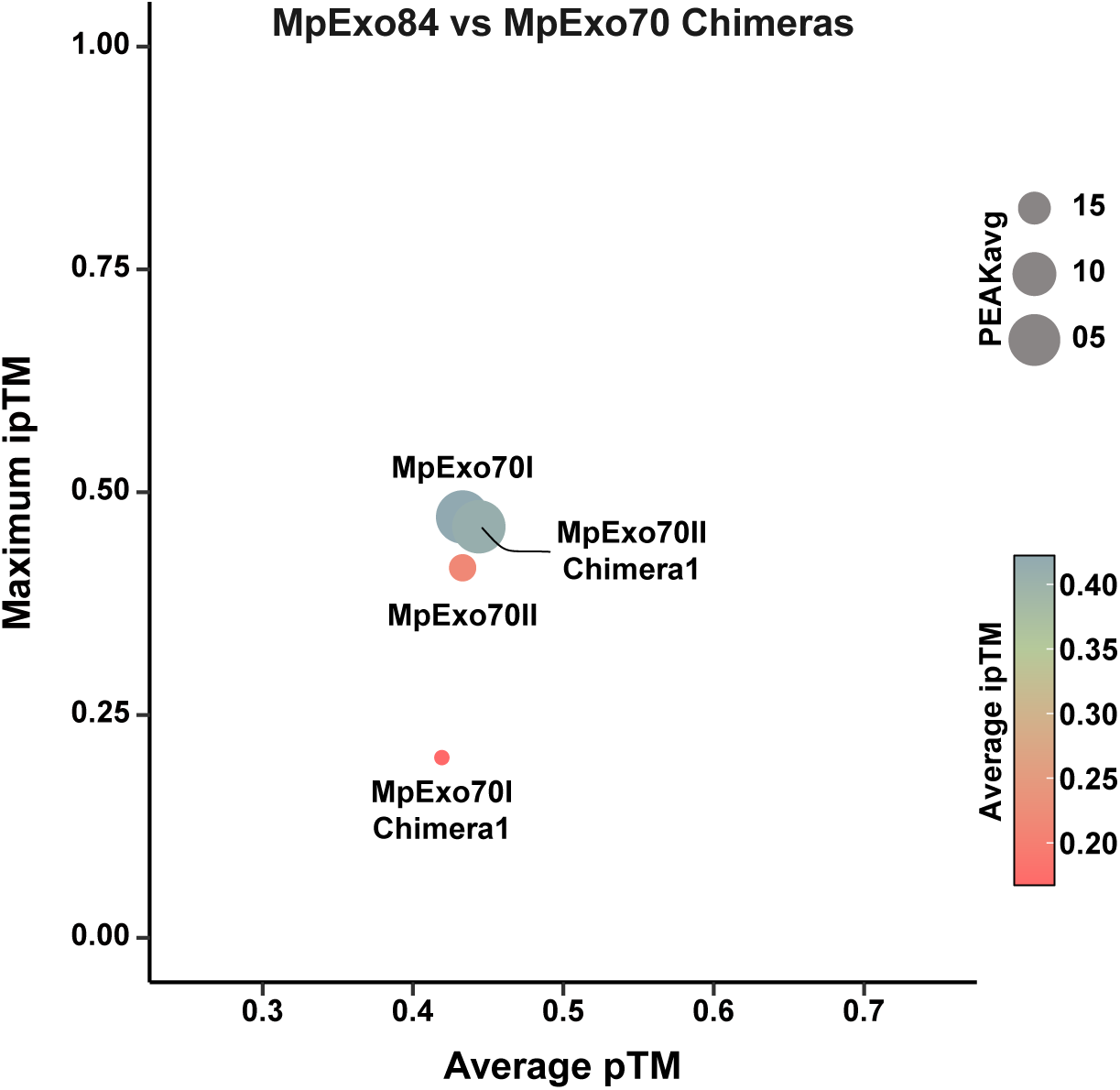
Analysis of MpExo70 chimera interaction with MpExo84 by AlphaFold2. Scatterplot of AlphaFold-Multimer (*78*) predicted ipTM versus pTM scores (ipTM: interface predicted Template Modeling score; pTM: predicted Template Modeling score) of the interaction between MpExo84 and MpExo70I, MpExo70II, MpExo70I chimera 1 or MpExo70II chimera 1. The maximum ipTM value from 5 independent predictions are use in the Y axis, while the average of pTM values from the 5 predictions is used in the X axis. The average ipTM is represented by the color of the dot. With the dot size correlating with the PEAK average. Dot size correlates to PEAK value average, where PEAK score represents average minimum predicted aligned error between protein chains excluding intra-molecular interactions.

**Fig. S30.**
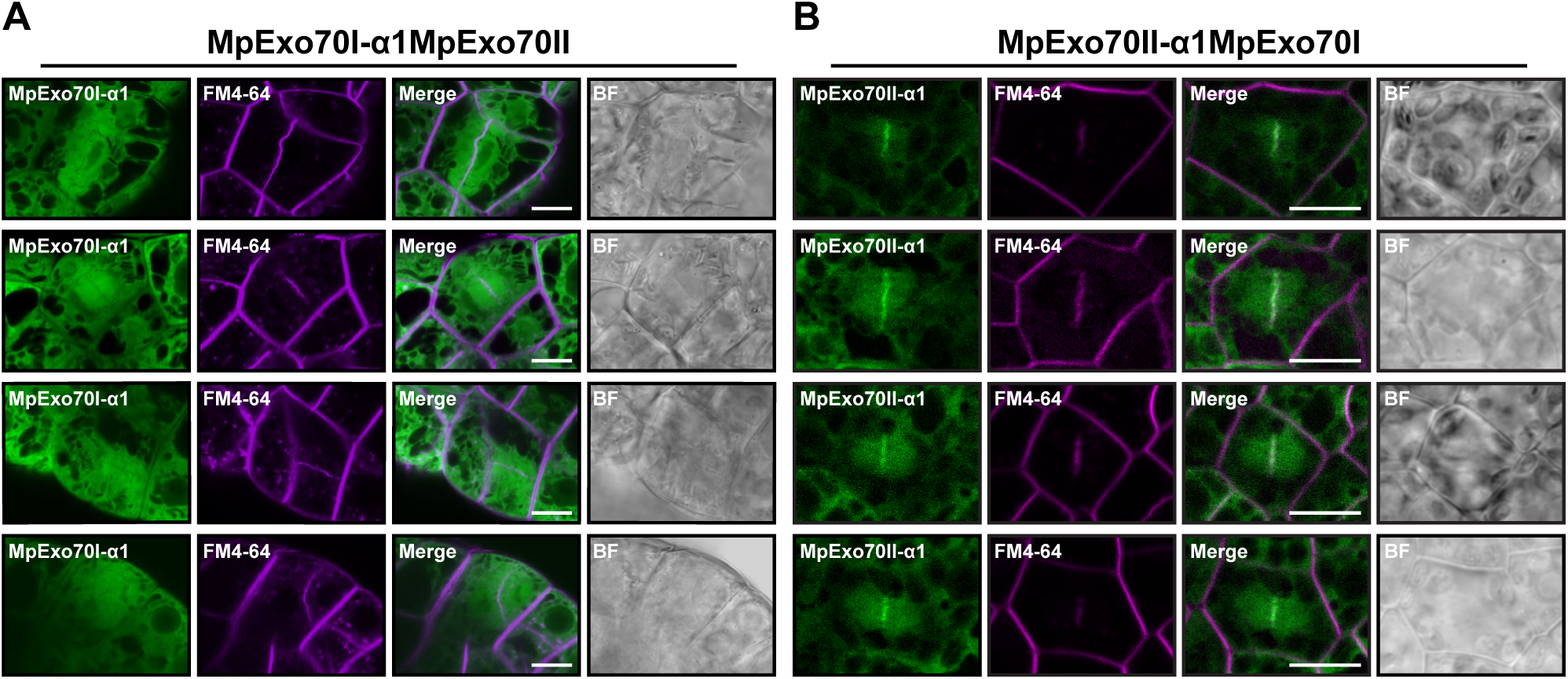
Localization of MpExo70 α1 chimeras in Marchantia. Micrograph collection of Marchantia cells stably expressing **(A)** MpExo70I-α1MpExo70II chimera:Clover and **(B)** MpExo70II-α1MpExo70I chimera:Clover stained with FM4-64 (magenta). The presence of the cell plate is depicted by accumulation of FM4-64 stain. Scale bar is 10 µm. BF indicates Bright Field.

**Fig. S31.**
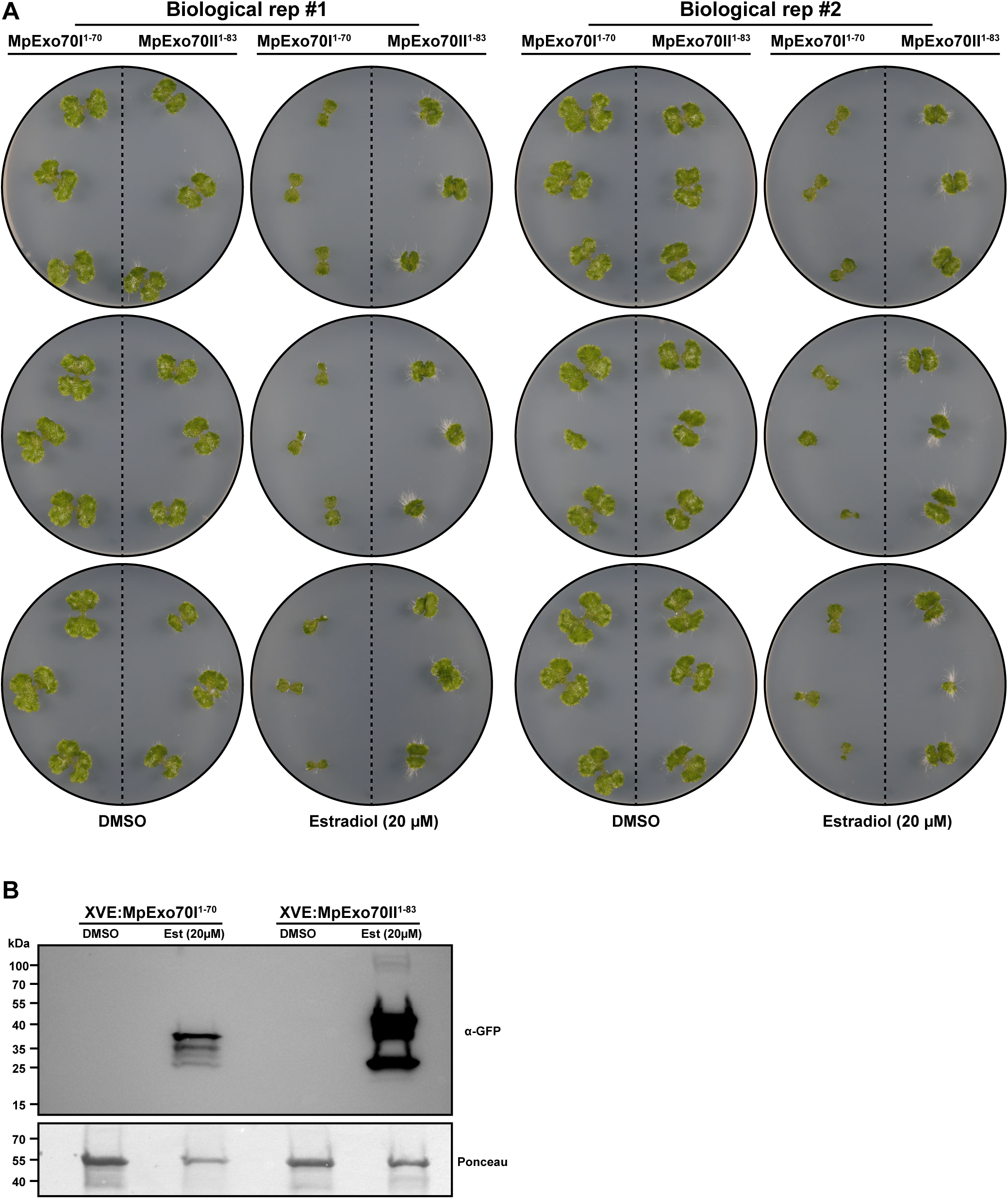
Experimental replicates of inducible expression of MpExo70 N-terminal domain expression in Marchantia. **(A)** Macroscopic phenotypes of Marchantia transgenic lines XVE:MpExo70I1-70:Clover and XVE:MpExo70II1-83 grown for 14 days on media containing B-estradiol (20 µM) or DMSO. **(B)** Accumulation of MpExo70I1-70:Clover and MpExo70II1-83:Clover after estradiol-mediated induction visualized by western blot.

**Fig. S32.**
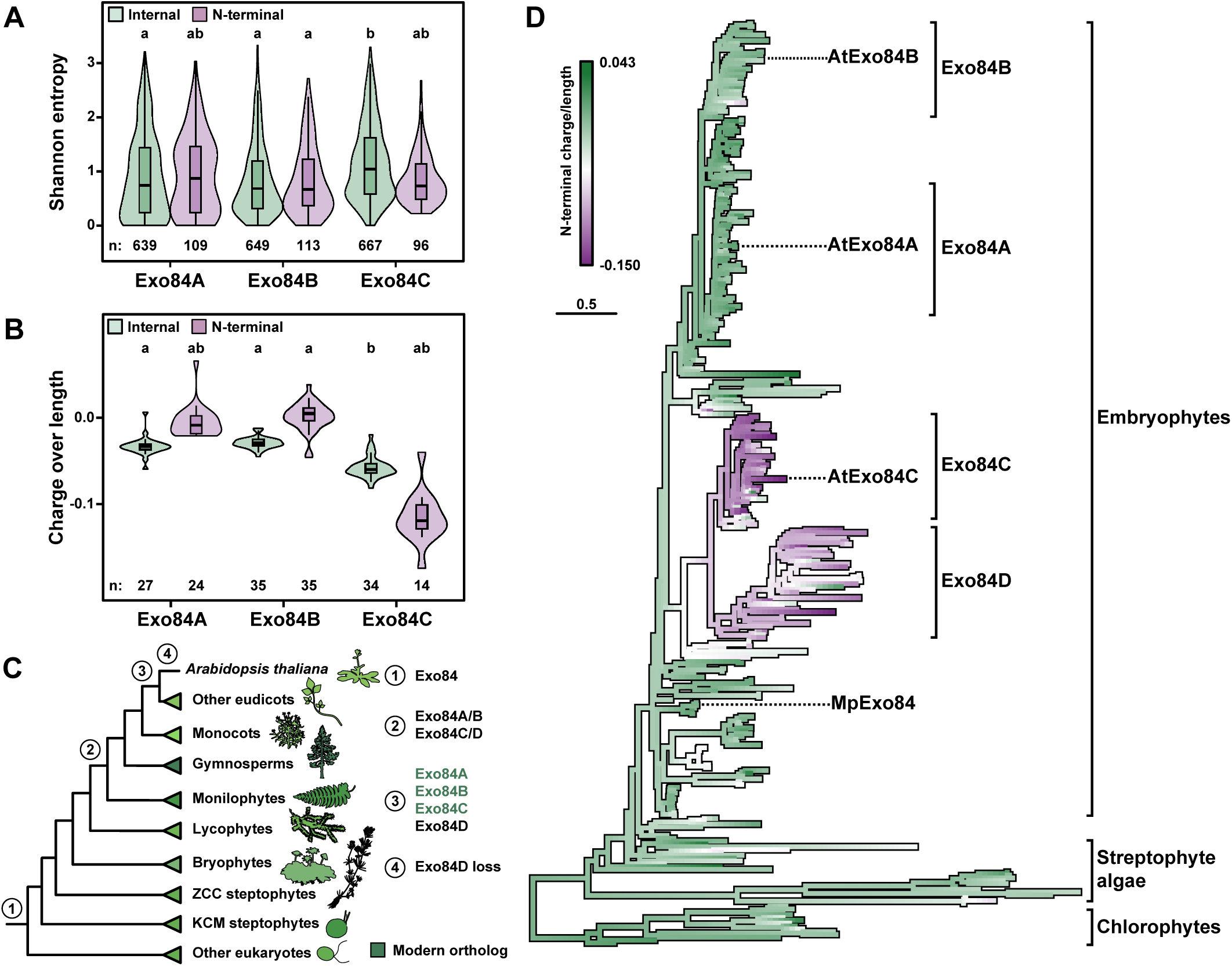
**(A)** Shannon entropy for aligned sites (*n*) in the N-terminal α-helix and the remainder of the protein (internal) of eudicot Exo84A, Exo84B, and Exo84C paralogs. The center line of the boxplots denotes the median and the upper and lower borders span from the first to the third quartiles, with whiskers extending 1.5 times the interquartile range. Distributions were compared using pairwise Tukey honestly significant difference (HSD) tests. Significance groups are denoted using compact letter display (P < 0.01 after Bonferroni multiple test correction). **(B)** Normalized electrostatic charge of the N-terminal α-helix compared to the rest of the protein for eudicot Exo84 paralogs. The number of analyzed proteins is noted (*n*). **(C)** Relative timing of the emergence of each of the eudicot Exo84 paralogs inferred using the Exo84 phylogeny in **Figure S37**. Taxonomic groups are noted with cartoons obtained from Phylopic.org. **(D)** A maximum likelihood phylogeny of the Exo84 family overlaid with ancestral state reconstructions of the normalized electrostatic charge of the N-terminal α-helices. *Arabidopsis thaliana* and *Marchantia polymorpha* paralogs as well as Exo84 subfamilies have been noted. The scale bar represents the average number of substitutions per site. Full phylogenies are available in **Figure S37** and from iTOL (*96*) (https://itol.embl.de/shared/OK75j4e8edHZ).

**Fig. S33.**
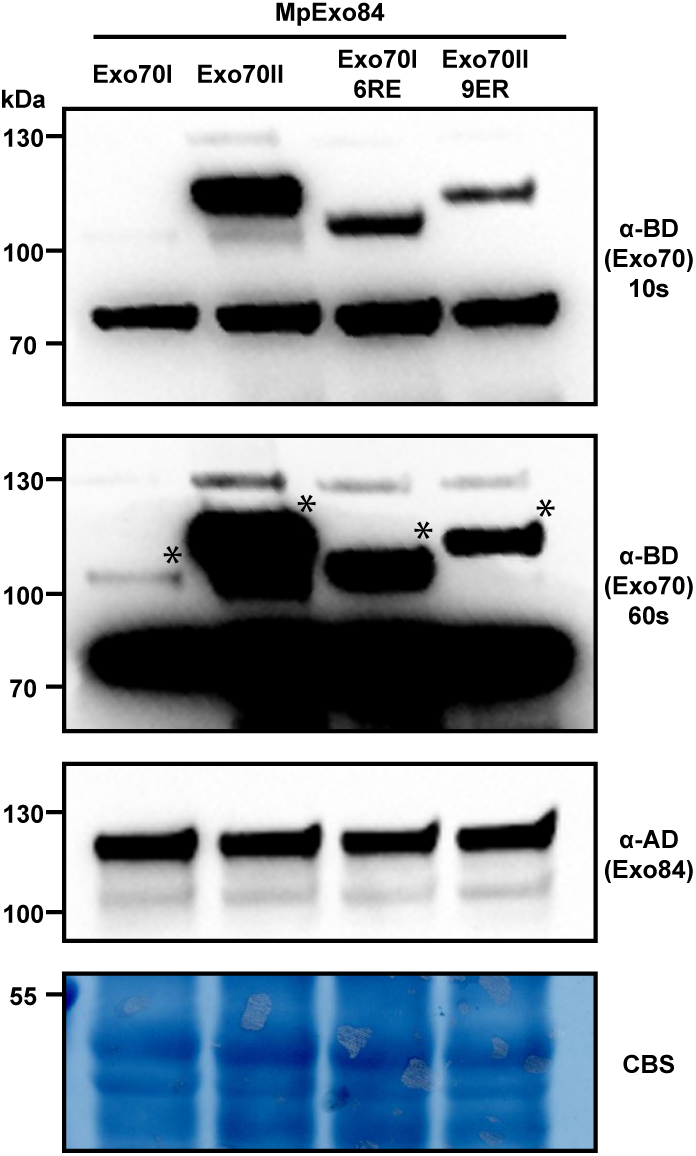
Protein accumulation in Yeast-Two-Hybrid assay analyzed by Western blot. Yeast lysate was probed for the presence of MpExo70I, MpExo70II, MpExo70I 6RE and MpExo70II 9ER using anti-GAL4 binding domain (BD); and MpExo84 using anti-GAL4 DNA activation domain (AD) antibodies. Total protein extracts were stained with Coomassie Blue Stain (CBS). Accumulation of MpExo70I was consistently lower and two panels with indicated exposure times are included. Asterisks indicate the bands corresponding to the proteins of interest.

**Fig. S34.**
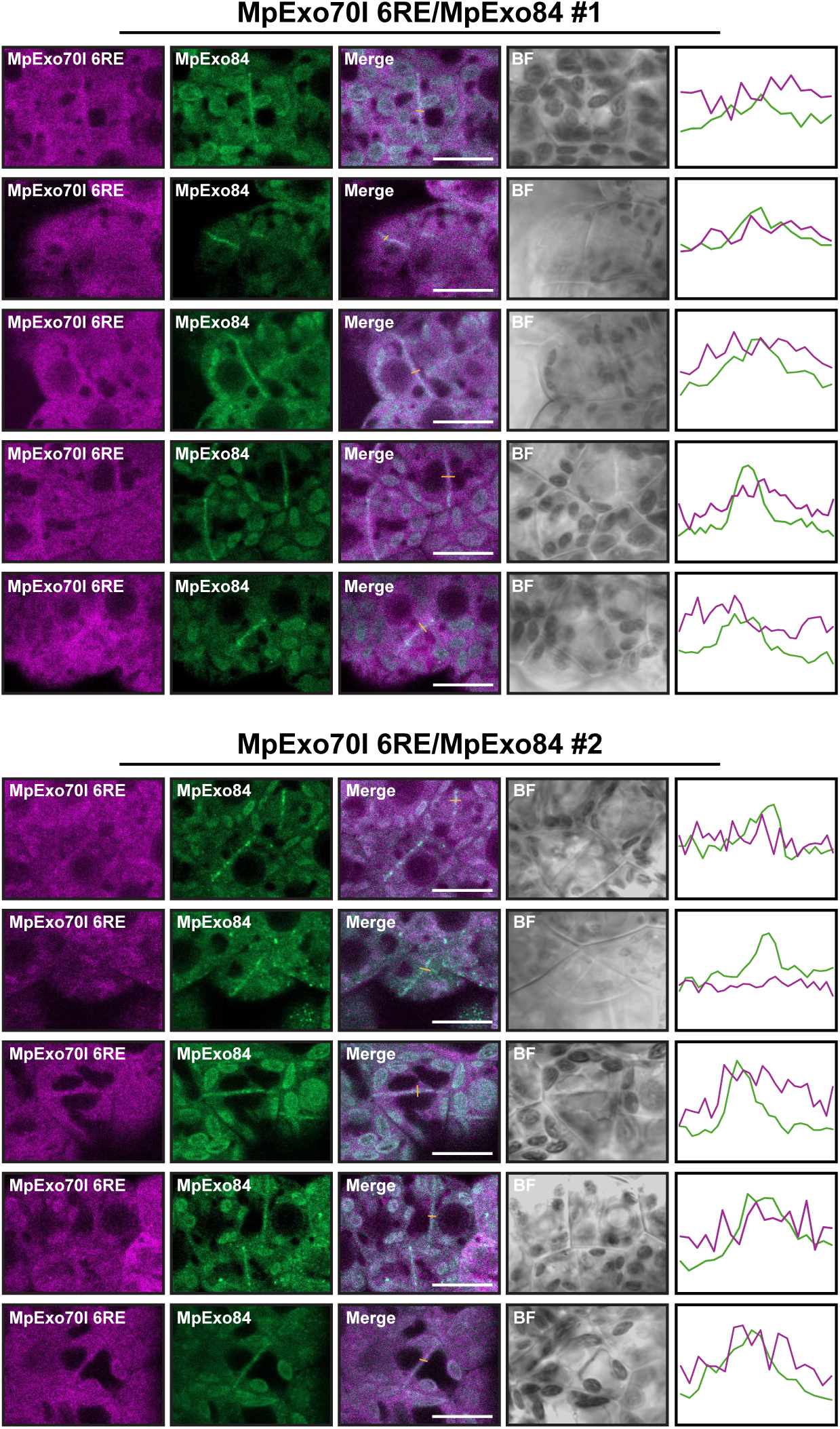
Image gallery for MpExo70I 6RE co-localization with MpExo84 in Marchantia. Micrograph collection of two independent Marchantia cell lines stably co-expressing MpExo84:Clover (green) with MpExo70I 6RE:mScarlet (magenta). The presence of the cell plate is depicted by accumulation of MpExo84:Clover. Right panels represent the fluorescence intensity profiles of Clover (green) and mScarlet (magenta) measured along the distance of the selected orange lines in merge channel. Scale bar is 10 µm. BF indicates Bright Field.

**Fig. S35.**
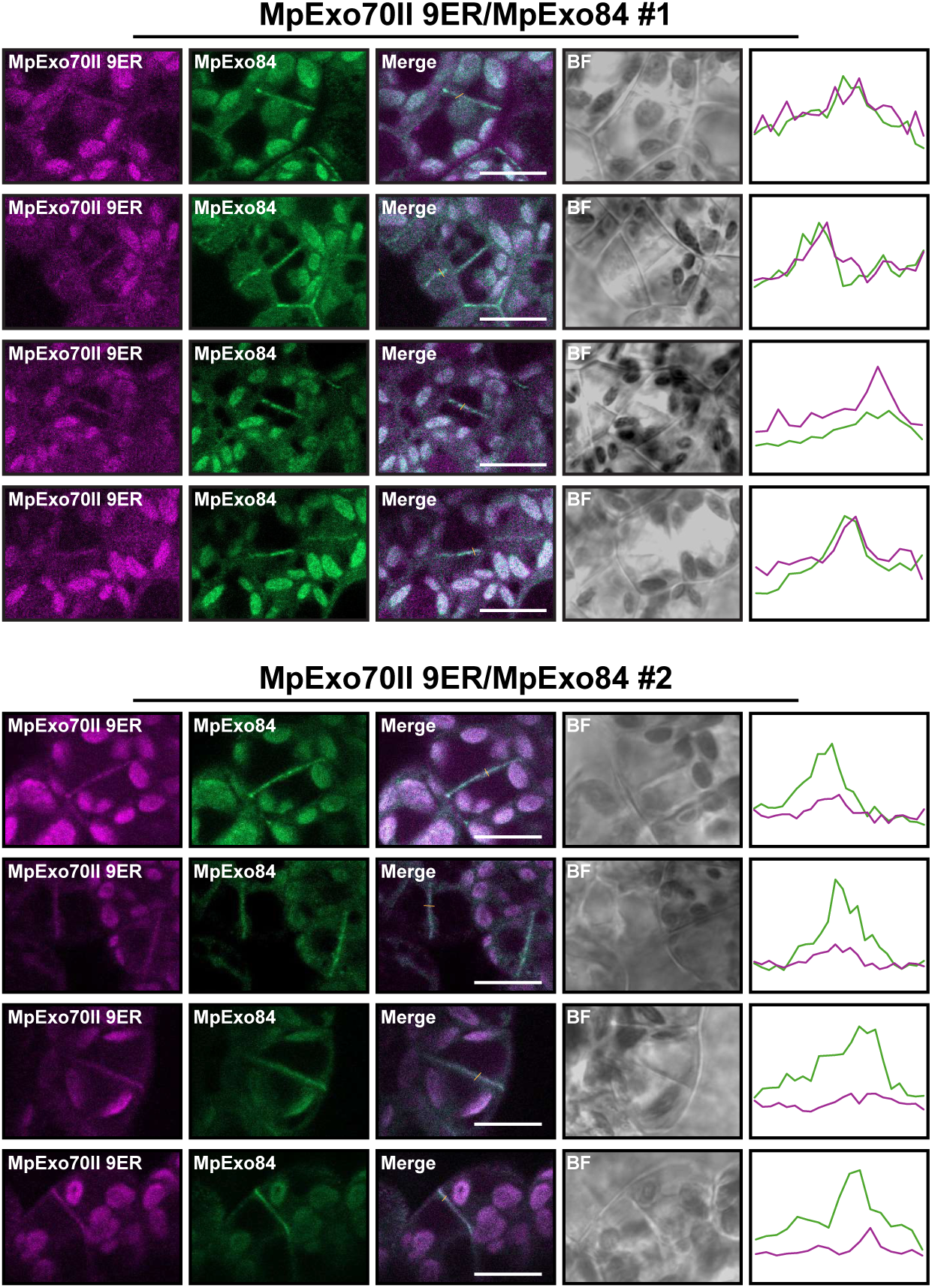
Image gallery for MpExo70II 9ER co-localization with MpExo84 in Marchantia. Micrograph collection of two independent Marchantia cell lines stably co-expressing MpExo84:Clover (green) with MpExo70I 9ER:mScarlet (magenta). The presence of the cell plate is depicted by accumulation of MpExo84:Clover. Right panels represent the fluorescence intensity profiles of Clover (green) and mScarlet (magenta) measured along the distance of the selected orange lines in merge channel. Scale bar is 10 µm. BF indicates Bright Field.

**Fig. S36.**
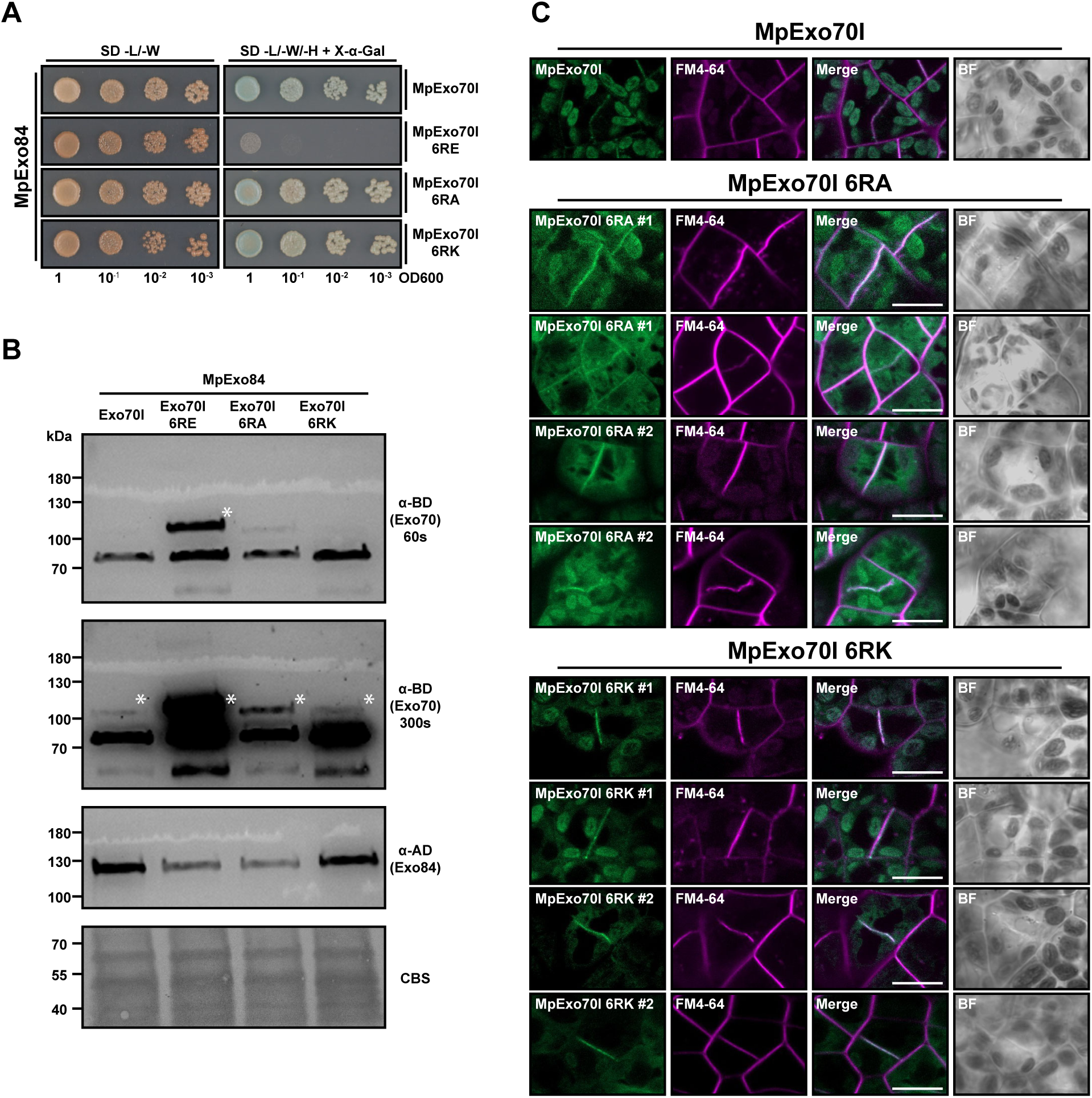
Substitution of MpExo70I Arg residues by Ala or Lys do not alter interaction with MpExo84 or cell plate localization. **(A)** Yeast two-hybrid assay of MpExo70I 6RA and MpExo70I 6RK interaction with MpExo84. For each combination, 5µl of yeast at indicated OD600 were spotted and incubated in double dropout plate for yeast growth control (left) and triple dropout media supplemented with X-α-gal (right). Growth and development of blue coloration in the right panel indicates protein-protein interactions. Wild-type MpExo70I and MpExo70I 6RE were included as positive and negative controls, respectively. MpExo70s were fused to the GAL4 DNA binding domain while MpExo84 was fused to the GAL4 activator domain. **(B)** Accumulation of each protein in yeast cells was probed for the presence of MpExo70I, MpExo70I 6RE, MpExo70I 6RA and MpExo70I 6RK using anti-GAL4 binding domain (BD); and MpExo84 using anti-GAL4 DNA activation domain (AD) antibodies. Total protein extracts were stained with Coomassie Blue Stain (CBS). Accumulation of MpExo70I 6RE was consistently higher and two panels with indicated exposure times are included. Asterisks indicate the bands corresponding to the proteins of interest. **(C)** Micrograph collection of two independent Marchantia cell lines stably expressing MpExo70I 6RA:Clover or MpExo70I 6RK: Clover stained with FM4-64 (magenta). A micrograph with Wild-type MpExo70I:Clover is included as control. The presence of the cell plate is depicted by accumulation of FM4-64 stain. Scale bar is 10 µm. BF indicates Bright Field.

**Fig. S37.**
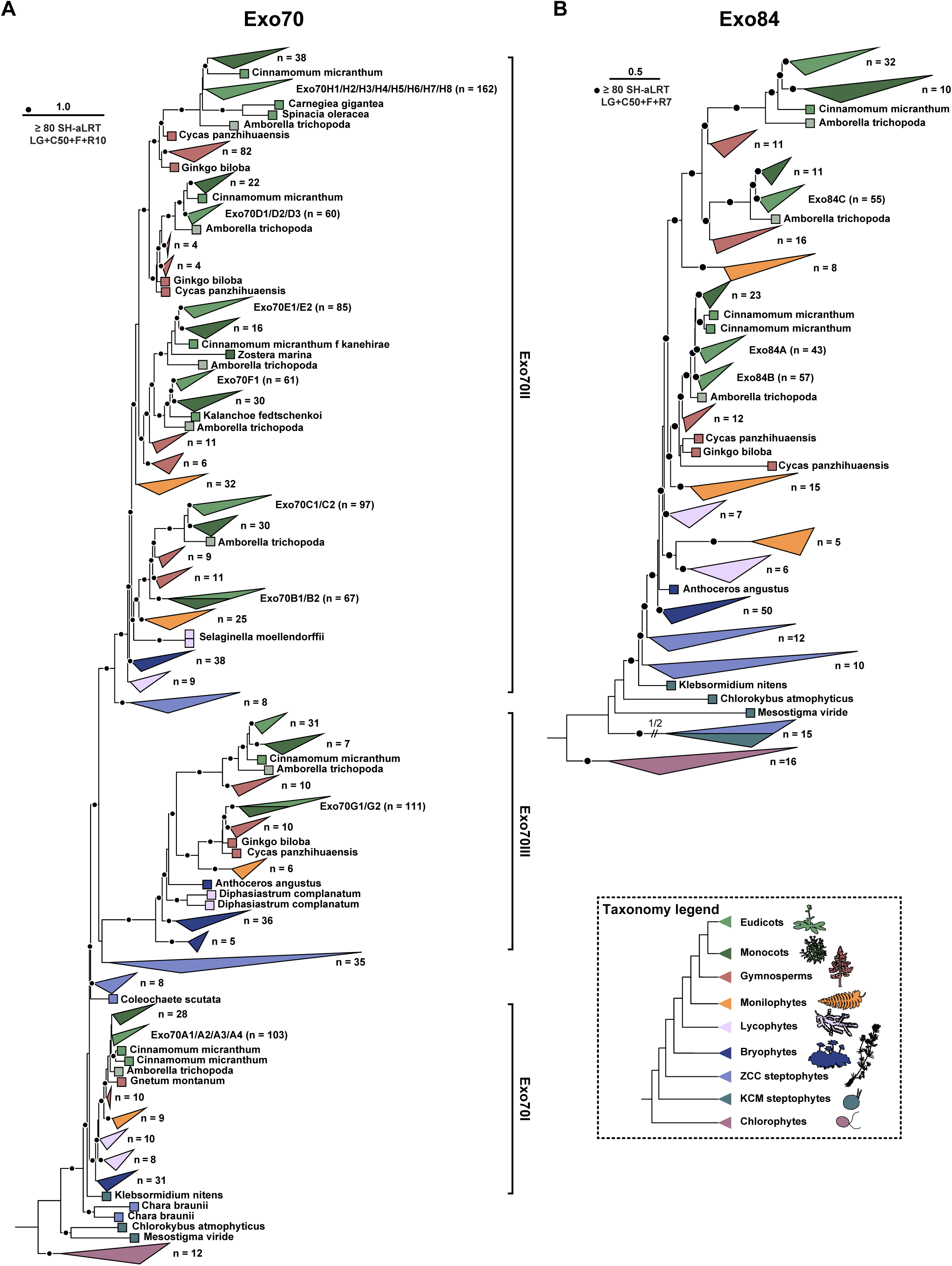
Maximum likelihood phylogenies of Exo70 **(A)** and Exo84 **(B)** in plants and green algae. Statistical support was inferred using ultrafast bootstrap (UFB) and Shimodaira-Hasegawa approximate likelihood ratio tests (SH-aLRT). The scale bars represent the average number of substitutions per site and clades have been colored taxonomically for clarity. Full phylogenies are available from iTOL (*96*) (https://itol.embl.de/shared/OK75j4e8edHZ).

**Fig. S38.**
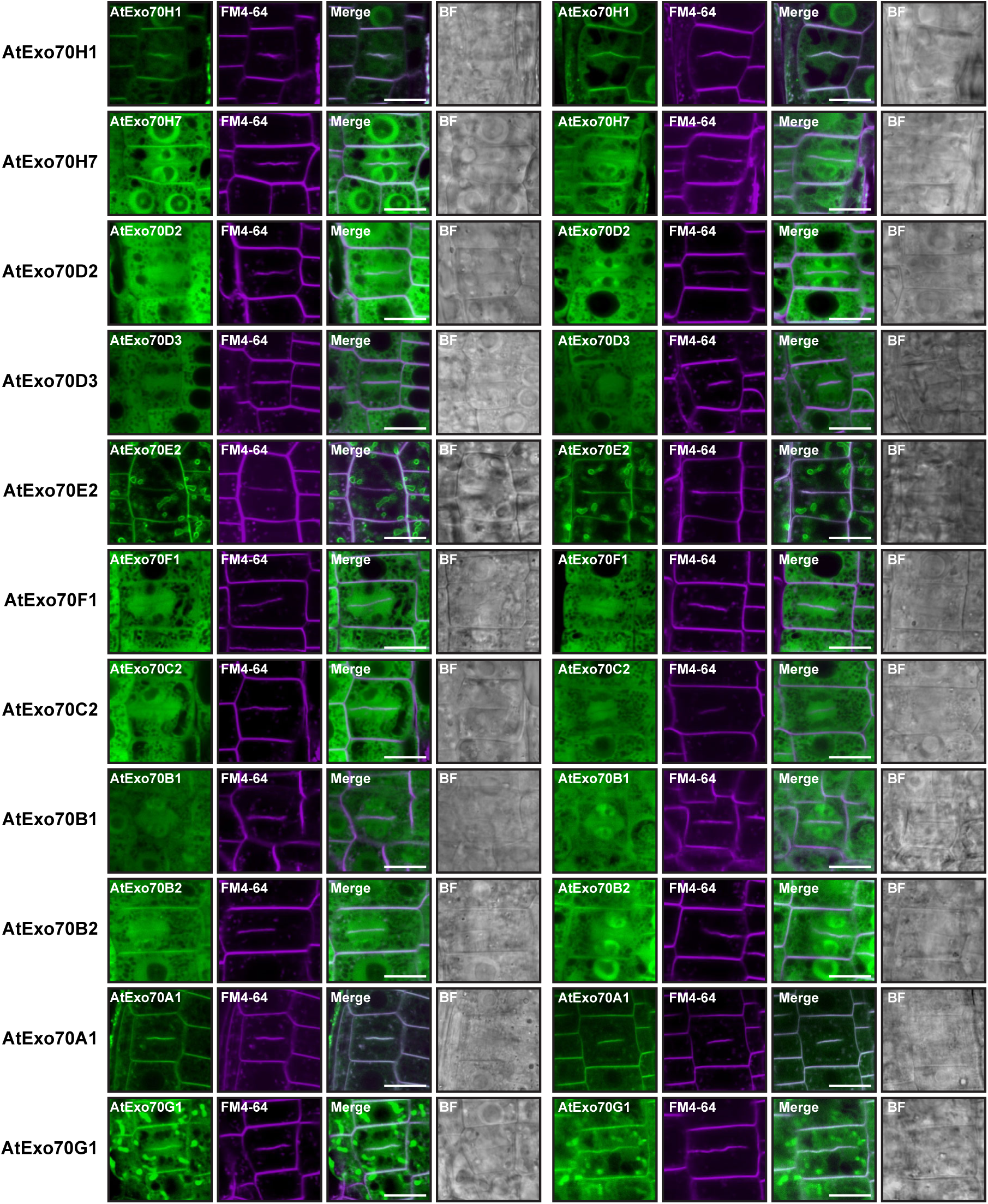
Image gallery for AtExo70 localization in Arabidopsis root cells. Micrograph collection of Arabidopsis root cells stably expressing C-terminally GFP tagged AtExo70 proteins (green) stained with FM4-64 (magenta). The presence of the cell plate is depicted by accumulation of FM4-64 stain. Scale bar is 10 µm. BF indicates Bright Field.

